# Variation in responses to temperature in admixed *Populus* genotypes predicts geographic shifts in regions where hybrids are favored

**DOI:** 10.1101/2025.05.16.654548

**Authors:** Alayna Mead, Joie R. Beasley-Bennett, Andrew Bleich, Dylan Fischer, Shelby Flint, Julie Golightly, Lee Kalcsits, Sara K. Klopf, Mason W. Kulbaba, Jesse R. Lasky, Jared M. LeBoldus, David B. Lowry, Nora Mitchell, Emily Moran, Jason P. Sexton, Kelsey L. Søndreli, Baxter Worthing, Michelle Zavala-Paez, Matthew C. Fitzpatrick, Jason Holliday, Stephen Keller, Jill A. Hamilton

## Abstract

- Plastic responses of plants to their environment vary as a result of genetic differentiation within and among species. To accurately predict rangewide responses to climate change, it is necessary to characterize genotype-specific reaction norms across the continuum of historic and future climate conditions comprising a species’ range.
- The North American hybrid zone of *Populus trichocarpa* and *P. balsamifera* represents a natural system that has been shaped by climate, geography, and introgression. We leverage a dataset containing 45 clonal genotypes from this natural hybrid zone, planted across 17 replicated common garden experiments spanning a broad climatic range. Growth and mortality were measured over two years, enabling us to model reaction norms for each genotype across these tested environments.
- Species ancestry and intraspecific genomic variation significantly influenced growth across environments, with genotypic variation in reaction norms reflecting a trade-off between cold tolerance and growth. Using modeled reaction norms for each genotype, we predicted that genotypes with more *P. trichocarpa* ancestry may gain an advantage under warmer climates.
- Spatial shifts of the hybrid zone could facilitate the spread of beneficial alleles into novel climates. These results highlight that genotypic variation in responses to temperature will have landscape-level effects.

## Introduction

Predicting the phenotypic responses of populations to changing climates and how they vary across species ranges is essential for conserving and restoring native ecosystems. Predicted changes in fitness between current and future climates can be used to identify populations most at risk, or to identify populations resilient to change which could be valuable seed sources for assisted gene flow and restoration (Aitken and Whitlock 2013; Aitken and Bemmels 2016). Phenotypes expressed in the field arise from the genotypic variation underlying phenotypic traits (G), the plastic response to the environment (E), and genotypic variation in response to the environment (GxE) (Des Marais et al. 2013). Each of these factors vary across complex and changing landscapes. By characterizing reaction norms, the set of phenotypes expressed across a range of environments, it is possible to predict organisms’ responses to the variable environments they inhabit and understand the selective forces that may be shaping plasticity (Via et al. 1995; Arnold et al. 2019).

Reaction norms can vary within species, resulting in part from selection imposed by spatially varying climates (Des Marais et al. 2013; Ikeda et al. 2017; Rehfeldt et al. 2018; Patsiou et al. 2020). Predicting genotype-specific responses will be particularly important for species with large ranges, which may exhibit variation in reaction norms due to the combined influence of geography, population demography, and climate (Cooper et al. 2019, 2022; Van Nuland et al. 2020). This is particularly true where glacial refugia have shaped species’ demographic history and connectivity (Woolbright et al. 2014; Love et al. 2023; Bolte et al. 2024). However, because of the difficulty of phenotyping multiple genotypes across many environments, many methods for modeling a species’ response to climate change ignore genotypic variation, and instead consider only the climate envelope of the extant species range (Capblancq et al. 2020). Furthermore, regions within a species’ range will vary in the rate, magnitude, and nature of climate change (e.g, increases or decreases in precipitation), making it important to understand localized climate responses. To accurately predict local phenotypic responses to changing climates, it is necessary to test whether reaction norms vary across genotypes, and if so, characterize genotype-specific reaction norms across the continuum of historic and future climate conditions comprising a species’ range (Arnold et al. 2019; VanWallendael et al. 2022).

Common garden experiments and provenance trials are invaluable tools for quantifying intraspecific variation in phenotypic plasticity and for predicting phenotypic responses to current and future environments (O’Neill et al. 2008; Wang et al. 2010; Leites et al. 2012; Fischer et al. 2014a; Grady et al. 2015; Browne et al. 2019; Leites and Benito Garzón 2023; Ye et al. 2023; Hord et al. 2025). Variation in reaction norms across genetic and climatic gradients can be quantified using repeated plantings of genotypes in multiple environments. Genotype-specific reaction norms modeled across continuous environments are used to identify genotypes or loci associated with optimized fitness or yield across varying environmental conditions; however, such studies have been limited to a handful of species (Gray et al. 2011; Arnold et al. 2019; Lowry et al. 2019; Patsiou et al. 2020; VanWallendael et al. 2022; Li et al. 2025). Because plasticity results from adaptation to climate as well as demographic history, it may be possible to predict the reaction norms of unmeasured genotypes if they vary as a function of climate of origin (O’Neill et al. 2008; Wang et al. 2010; Rehfeldt et al. 2018) or genetic variation.

When genomic data are available in addition to phenotypic and climate data collected in common garden experiments, predictions of phenotypic responses can be improved (Mahony et al. 2020; Archambeau et al. 2022; Putra et al. 2023; Li et al. 2025). Therefore, incorporating genetic information into characterization of reaction norms will enable more accurate predictions of rangewide climate responses.

Natural hybrid zones are ideal systems for disentangling the effects of adaptation and demographic history on phenotypes, including reaction norms, because multiple generations of backcrossing can result in novel recombinant genotypes associated with high phenotypic variability (Janes and Hamilton 2017). Hybrid zones can increase the genetic variation available for adaptation to rapidly changing climates and allow movement of adaptive loci from one species into another via introgression (Janes and Hamilton 2017; Suarez-Gonzalez et al. 2018b; Kremer and Hipp 2020; Buck et al. 2023; Hord et al. 2025). Comparing responses to the environment across admixed genotypes reveals how existing genetic variation within hybrid zones underlies fitness differences across environments.

Hybrid zones can also be used to monitor responses to climate change (Taylor et al. 2015), with documented geographic shifts in some zones in response to changing environments (Billerman et al. 2016; Wielstra 2019; Alexander et al. 2022). If reaction norms depart from the intraspecific pattern across a hybrid zone, climate change may alter regions where particular species, genotypes, or even genes are favored (Hord et al. 2025).

North American *Populus* is a model system for forest trees due to their small genomes, ease of clonal propagation, and development as a biofuel feedstock (Jansson and Douglas 2007; Sannigrahi et al. 2010; Porth and El-Kassaby 2015). Yet, like many non-model trees, its genetic variation is shaped by interactions between climate, geography, and interspecific introgression. *Populus* species occupy heterogeneous landscapes, often forming multi-species hybrid zones that are a source of novel recombinant genetic variation (Suarez-Gonzalez et al. 2016, 2018a; Chhatre et al. 2018; Bolte et al. 2024). These factors make *Populus* an ideal system for quantifying intraspecific variation in climate responses and predicting landscape-scale changes in fitness. Here, we focus on the North American hybrid zone between *P. trichocarpa*, a western species spanning latitudes from Alaska to California, and *P. balsamifera*, which occurs transcontinentally throughout the boreal regions of the contiguous United States, Canada and Alaska (Figure 1A). Populations within the hybrid zone exhibit genetic and phenotypic differentiation, likely resulting from varying demographic history as well as adaptation to wide-ranging biotic and abiotic environments (Keller et al. 2010, 2011, 2012; Slavov et al. 2012; Evans et al. 2014; Geraldes et al. 2014; McKown et al. 2014a, c,b; Zhou et al. 2014; Fitzpatrick and Keller 2015; Holliday et al. 2016; Chhetri et al. 2019; Zhang et al. 2019; Fitzpatrick et al. 2021). Gene flow rates across the hybrid zone vary in part due to geographic barriers, resulting in spatial differences in population differentiation (Bolte et al. 2024).

**Figure 1.**
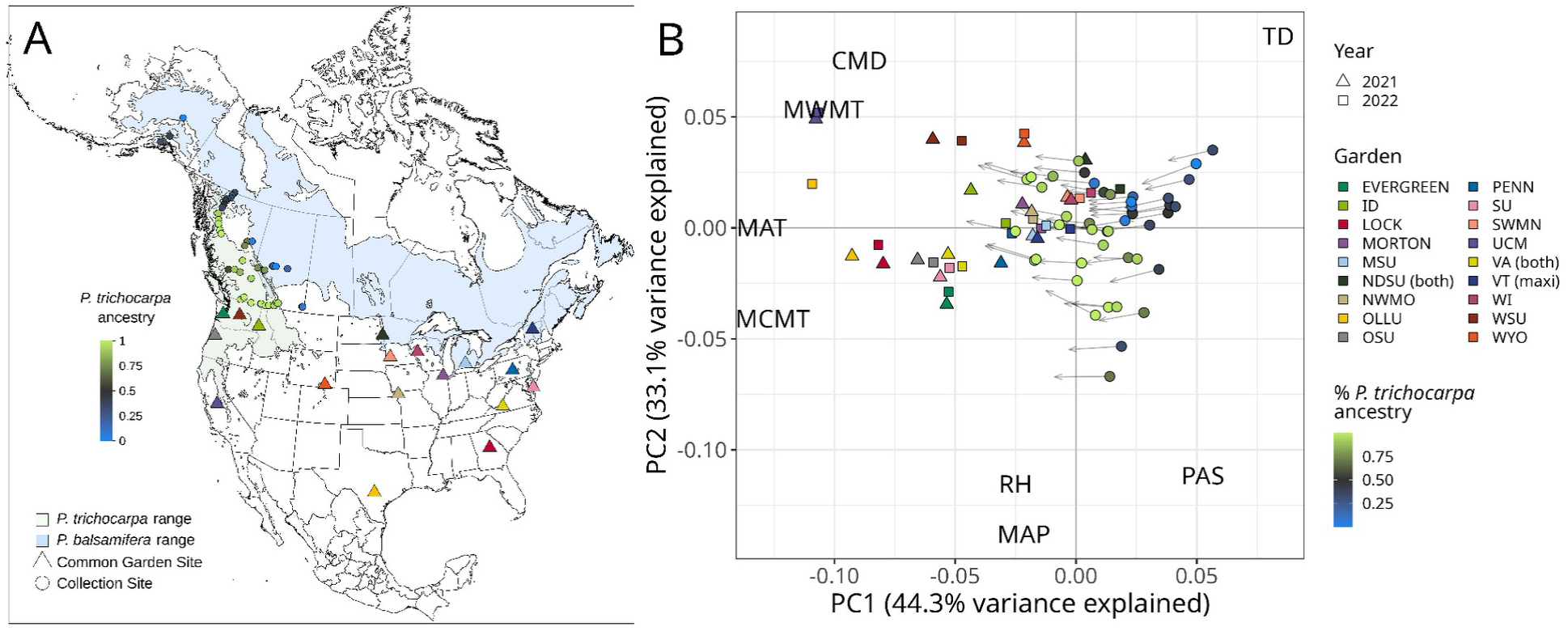
(A) Map of sampled genotype localities (circles) and common garden sites (triangles) in relation to the ranges of both species (Little 1971); color of sampled genotypes indicates species ancestry calculated from ADMIXTURE at K=2. Sites of maxi gardens and mini gardens are noted in the legend. (B) PCA of past and future climates across collection and garden sites. Gardens were generally warmer than home sites (lower PC1 loadings), and their position in multivariate climate space in PC1 and PC2 overlapped with the expected future climates for some genotypes. Circles represent the historic climate (1961-1990) of each genotype’s provenance, colored by species ancestry, and arrowheads represent the predicted climate for 2041-2070 under an ensemble of 13 GCMs.

We leveraged the *P. trichocarpa* × *P. balsamifera* hybrid zone to determine how genetic and environmental parameters interact to influence genotype-specific reaction norms, ultimately using our predictions to model changes to hybrid zone composition in the context of global change. We used a series of replicated provenance trials of clonal genotypes from the hybrid zone, which we planted in 17 gardens across the United States that span a wide range of environments, including many warmer than the climate of origin. We tested three hypotheses: Hypothesis 1:*Populus* hybrids display heritable differences in adaptation to climate, which manifest as genotype-specific responses to climatic variation. To test this, we measured fitness-related traits in each garden to determine how genetic background and the environment interact to determine fitness in this system, quantifying the effect of genetic structure (genotype effect, G), climate (environmental effect, E) and how the response to climate varies by genotype (G×E). A significant genotype × environment interaction could suggest that the response to climate is mediated by local adaptation to the climate of origin. Hypothesis 2: Variation in reaction norms are predictable based on a continuous gradient of genomic ancestry, enabling us to predict where the hybrid zone is favored to move under future climates. To test this, we used whole-genome sequence data to characterize multivariate genetic structure, including both species ancestry and intraspecific variation, and predicted its effect on phenotypic responses among genotypes. We developed a model to predict growth and mortality for a given garden environment using information about each genotype’s climate of origin and genetic data, which was validated using a combination of 500 novel genotypes and novel environments. Finally, we used genotype-specific reaction norms to predict phenotypic responses to climate change across the landscape of the hybrid zone. Hypothesis 3: Inclusion of genetic information in phenotypic response models improves their predictions due to inherent genetic structure contributing to variation in reaction norms. We tested whether climate of origin can serve as a proxy for adaptive genetic structure, circumventing the need for genetic data.

## Materials and Methods

### Field collections and propagation

In 2020 we established 17 common gardens containing 48 clonally replicated poplar genotypes, including *P. trichocarpa*, *P. balsamifera*, and admixed individuals. Clonal replicates were propagated from dormant vegetative cuttings taken from 48 different mature trees in the wild between October 2019 and March 2020. These genotypes originated from natural populations spanning five transects that traverse the natural hybrid zone of *P. trichocarpa* and *P. balsamifera*, including latitudes from Alaska to southern British Columbia and Alberta (Figure 1). Dormant vegetative cuttings were transported to Virginia Tech (Critz, VA, USA) for propagation, as described in Bolte et al. (2024).

Briefly, cuttings were exposed to a 30-second ZeroTol 2.0 fungal dip treatment, dipped in Garden Safe Take Root hormone (0.1% indole-3-butyric acid), rooted into a standard rooting mix, and placed on a mist bench for 38 days following which clonal genotypes were shipped and planted into common garden sites.

### Common garden design

Seventeen common gardens were established at colleges, universities, and arboreta across the United States in fall 2020 (Figure 1A and Table S1). We visualized variation in climate space among garden environments and past and future home environments using a PCA of environmental variables (Figure 1B-C). Garden sites span a range of environments and include the southern range of both *P. trichocarpa* and *P. balsamifera*, as well as warmer regions to the south of their native ranges, enabling us to predict responses to novel climates (Figure 1B). Each garden included two blocks, with one individual per genotype planted across each block in a randomized complete block design. In total, the design included 48 genotypes x 17 gardens x 2 blocks x 1 genotype/block. Each garden had a minimum of 94 trees, for a total of 1656 individuals included across all gardens. Phenotypic data were collected during the growing season for three years: 2021-2023. All 17 gardens were evaluated in the first year, but high mortality and changing local phenotyping capacity at some sites decreased the total number of gardens assessed to 15 and 12 sites in 2022 and 2023, respectively (Table S1).

As part of the same experiment, three additional common gardens were established in March 2020. These gardens included 544 genotypes, each with three clonally replicated individuals planted in three blocks (1 replicate per block), located at North Dakota State University (NDSU), Virginia Tech University (VA), and the University of Vermont (VT). We designated the 17 smaller gardens as “mini” gardens, and the three larger gardens as “maxi” gardens. All 48 genotypes included in the mini gardens were also planted in the maxi gardens. Here we focus on phenotypic variation across the mini garden environments and use the maxi gardens for model evaluation (see statistical methods below).

### Phenotypic data

Height was measured each year (2021-2023) prior to bud flush and after budset. Annual growth increment was calculated per year based on the height accumulated during the growing season (pre-bud flush height subtracted from post-budset height). Some gardens experienced herbivory, disease, and accidental mechanical damage leading to negative growth increments for some individuals. As these negative growth increments likely represent the consequences of measurement error or herbivory, they were removed prior to analyses (137 individuals in 2021 and 97 in 2022). Our goal was to isolate the effects of climate on growth. While impacts of herbivory will also likely shift with climate change, they may vary outside the native range, and herbivory also likely differed across gardens for non-climatic reasons (e.g., fencing and human presence), so we excluded its effects here. Given the frequency of herbivory, it is likely that some growth increment values are underestimates of potential yearly growth.

### Climate data

Climate data were extracted from ClimateNA rasters (Wang et al. 2016; AdaptWest Project 2022) using terra 1.8-21 (Hijmans 2025) in R version 4.4.2 (R Core Team 2024) for genotype provenances (locations of origin) and common garden environments. For provenances, we extracted historical climate data for the 30-year period from 1961-1990. We focused on average historical climate data because variation in climate response is partially the result of selection associated with climate of origin during establishment and over the lifespan of the collected adult trees, which average >30 years old.

For garden climates, we extracted the climate averages associated with each year of data collection, starting when the seedlings were planted in 2020. We also fit the model (described below) with 2021 data using 22 climate variables as the home and garden climates, dropping the block effect to allow all models to converge, and compared their AIC scores. We selected mean coldest month temperature (MCMT) for use in statistical modeling because it explained variation in performance across gardens and had the lowest AIC score (Table S2). Moreover, the range of MCMT values at the common garden sites encompassed climates for most natural populations (see Results). This enabled us to predict growth and mortality responses to MCMT within much of the native range of the two poplar species. We did not use precipitation variables in statistical modeling because most gardens were irrigated during the first year, and some drier gardens (OLLU, SWMN, and UCM) continued irrigation in later years to ensure survival, limiting the selective effect of precipitation variation across gardens.

### Genomic data

Whole-genome sequence data associated with each genotype and previously described in Bolte et al. (2024) was used in phenotypic prediction across common gardens. DNA extraction, sequencing, and variant filtration are described in Supplementary Methods and filtering scripts are available at https://github.com/alaynamead/poplar_hybrid_vcf_filtering. A total of 334,657 variable sites were used to characterize genetic variation and admixture. Bolte et al. (2024) identified three distinct lineages in this hybrid zone: *P. balsamifera*, coastal *P. trichocarpa*, and a separate interior *P. trichocarpa* lineage originating from an ancient admixture event between the two species. We incorporated this genetic variation into our predictions using the PC scores of each genotype from a PCA of SNP data created with vegan 2.6-8 in R (Oksanen et al. 2024), defining genetic structure as variation in genomic PC space. PC1 separates *P. balsamifera* and the two *P. trichocarpa* lineages, PC2 separates all three lineages, and PC3 separates the northern (Alaska and Cassiar) and southern (Chilcotin, Jasper, and Crowsnest) transects (**Figure S1**). To visualize how responses varied by species ancestry, we used ancestry proportions based on K=2 from the ancestry estimation software ADMIXTURE (Alexander et al. 2009). Across 48 genotypes, three genotypes were previously identified as genetic outliers, possibly as a result of mixed ancestry with additional *Populus* species (Bolte et al. 2024, Figure S2). These individuals were removed from the analyses, leaving a total of 45 genotypes for statistical modeling.

### Statistical model

We fit a generalized linear mixed model to evaluate the effects of garden climate, home climate, genetic structure and their interactions (G×E) on growth across the common gardens and to predict responses to future climates. We defined environment as the MCMT of each garden during the year of measurement, and we defined home climate as the historical average of MCMT (1961-1980) at the provenance origin for each genotype. Genotypes may vary in their responses to climate as a result of local adaptation to their climate of origin and as a result of neutral genetic variation. Given this, we accounted for these sources of variation using different model variables. Previous studies have tested how climate of origin determines genotypic variation in responses to the planting environment, assuming that responses are largely explained by local adaptation and modeling G×E using climate of origin as a proxy for genetic variation (O’Neill et al. 2008; Wang et al. 2010). However, this approach does not account for variation in responses that result from other processes, such as neutral evolution resulting from historic demographic processes, selection for recent but not long-term climate patterns, or novel recombinants originating from hybridization. We include both genetic structure and climate of origin in the model to account for these multiple interacting factors that may influence the response to the environment.

We included data from the first two years of measurement (2021 and 2022) across 17 and 14 gardens, respectively, analyzing 2768 measurements of 1610 individual trees. We excluded 2023 data because gardens at the upper and lower temperature extremes were lost due to high mortality (Figure S1), and climatic extremes are important to provide “anchor points” to produce biologically realistic response curves (Wang et al. 2006). However, we did compare the model fit for each year individually and observed similar effect directions across all three years (Supplementary Figure S10).

We included survival and yearly growth increment as two measures of performance within the same model by fitting a zero-inflated Gaussian model using glmmTMB 1.1.11 (Brooks et al. 2017) in R version 4.5.0 (R Core Team 2024). We set the growth of all trees marked dead at the end of each growing season as 0, resulting in a large number of zero values (381 trees in 2021 and 343 trees in 2022, 27% of the total). This model assumes that values of zero could have two origins, which were modeled as two separate components within one model. The first “conditional” component modeled “sampling” zeros as part of the continuous distribution of growth increment (i.e., some individuals survived but had growth at or near zero). The second “zero-inflated” component modeled “structural” zeros attributed to mortality. These are modeled by the zero-inflated component, taking binary mortality values as input to predict the probability of mortality using a logit link function (Hu et al. 2011; Brooks et al. 2017). This method allowed us to evaluate how two fitness-associated traits, growth and mortality, separately respond to MCMT. Together their combined effect can be considered a proxy for overall fitness, incorporating both the probability of mortality and the predicted growth for surviving individuals. We report phenotypic predictions for three model components: the zero-inflated component representing mortality, the conditional component representing growth, and the overall model combining the two.

The full model included the fixed effects of garden MCMT, home MCMT, their square terms (to account for the observed quadratic response associated with a fitness optimum, Figure S2 and S3), and all interactions between garden and home MCMT and their square terms (indicating that the response to garden temperature varies based on temperature of origin). We also included population genetic structure as a fixed effect using the values for genetic PC1, PC2, and PC3 (Figure S1). MCMT was negatively correlated with genetic PC1 (R = −0.69), consistent with the climate niches associated with the two species, but was only weakly correlated with PC2 (R=-0.22) and PC3 (−0.40), so including multiple PCs enables us to identify variation in responses resulting from neutral genetic structure or adaptation to other climate variables. We tested for G×E by including interaction effects between garden climate and the genetic PCs. We included the random intercepts of individual, genotype, year, and block nested within garden. The same formula was used for the zero-inflated portion of the model to test the effect of each factor on mortality. To improve model convergence, we scaled the numeric variables using the R function `scale` without centering and back-transformed scaled values to actual values for visualization. Growth increment was log-transformed to account for a distribution skewed towards lower values. Using each individual as a separate observation, we fit the following model using the `glmmTMB` function, accounting for home MCMT, garden MCMT and their interactions and as well as genetic PCs 1-3 and their interaction terms with garden MCMT. This modeled the *j*th individual at the *i*th garden, where μ represents the intercept, H_j_ is the home environment and G_i_ is the garden environment i; A_j_, B_j_, and C_j_ are respectively genetic PCs 1, 2, and 3 for individual j; block, genotype, year, and individual are random intercepts, and ɛ represents the error term.

## Model 1

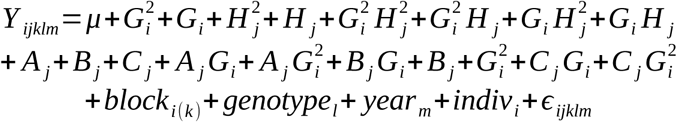

The significance of each term was evaluated using the `tab_model` function from the R package sjPlot 2.8.17 (Lüdecke 2024), which uses a type II Wald chi square test. Estimates for each factor were standardized by dividing by its standard deviation using tab_model and plot_model functions from sjPlot to facilitate comparisons between the relative effect sizes of linear and quadratic terms and their interactions.

### Model evaluation

To evaluate whether Model 1 could accurately predict poplar phenotypic response to environmental conditions, we compared its predictions of growth and mortality to actual measures in two ways. For both methods, we predicted the growth increment and mortality for each tree, using the garden MCMT and each genotype’s home MCMT values and genomic PC values as model parameters. First, we removed one of the garden environments from the dataset, trained the model on the remaining 16 gardens, and used the fitted model to predict growth increment in the garden that was excluded, repeating this process for each garden (hereafter referred to as leave-one-out cross validation). Second, we trained the model using all 17 gardens and then predicted growth increments of 544 genotypes in the three maxi gardens. Two maxi garden sites also have a mini garden site that was included in the training data (Virginia and North Dakota) and one is a novel environment without a mini garden (University of Vermont). 500 of these genotypes were novel genotypes not evaluated within the mini gardens, and we used their genomic PC values and home climate to predict their responses to garden climate. We predicted growth increments for individuals based on the full model using the `predict` function from the glmmTMB package and calculated the Pearson correlation between the actual values and the predicted values. We also tested whether the estimated probability of mortality corresponded to measured mortality rates using a generalized linear model with a binomial link function. Full details of methods used to predict phenotypes are included in the supplemental methods.

We used the same leave-one-out cross-validation method to compare the performance of three different models with different combinations of genetic and climatic information. If most of the variation among genotypes in response to garden MCMT is explained by local adaptation, provenance MCMT could serve as a proxy for genetic structure, enabling genotype-specific predictions of responses to temperature without the need for genetic data. To determine whether genetic PCs or home temperature increased the predictive power of the model, we tested simplified versions of the full model described above: Model 2, a genetics-only model with home temperature excluded to test the predictive power of the genetic PCs, and Model 3, a temperature-only model with genetic information excluded to test the predictive power of home MCMT (Table 1). We trained the three models on datasets with each garden excluded and predicted responses in the excluded garden for 2021 and 2022, evaluating models using the Pearson correlation between actual and predicted growth increments as described above. We tested whether the models differed in predictive ability using an ANOVA, with the 17 cross-validation tests used as replicates.

**Table 1.** Factors included in each model used for testing whether home climate can be a proxy for genetic structure in modeling the response to environment.

|  | **E:** Garden<br>MCMT<br>(quadratic) | **G:** Genetic<br>PCs 1-3 | Genotype x<br>garden MCMT<br>**(GxE)** | Home MCMT<br>x Garden<br>MCMT<br>(quadratic)<br>**(GxE proxy)** |
| --- | --- | --- | --- | --- |
| Model 1: Full | x | x | x | x |
| Model 2:<br>Genetics only | x | x | x |  |
| Model 3:<br>Climate only | x |  |  | x |

### Predicting the response to temperature

We used the parameters estimated from the full model (Model 1) to predict each genotype’s norm of reaction; i.e. the response to a continuous temperature gradient, allowing us to characterize responses to temperature across the hybrid zone. We predicted the growth increment and mortality probability of each genotype across a range of 100 equally-spaced MCMT values from −23.9 to 9.8 °C, encompassing both historic 30-year home climates (−23.9 to −3.8 °C) and yearly garden climates (−16.5 to 9.8 °C). Phenotypic predictions should be more reliable for the warmer climatic range included in the common gardens where phenotypic data was collected than for colder regions, enabling us to predict responses to warming climates. We predicted the phenotypic response of each genotype across this climatic range of MCMT, accounting for genotype-specific responses with the genetic PCs and the home MCMT of that genotype, using the parameters estimated from Model 1. The glmmTMB `predict` function was used, ignoring the random effects of genotype, block, and garden and predicting the overall response to temperature rather than the genotype-or garden-specific response (option re.form = NA). Phenotypic predictions were made for the two model components and for their combined effect: the conditional component reflecting growth increment, the zero-inflated component predicting the probability of mortality, and the combined model incorporating both. We also calculated the climate transfer distance as the difference between garden and home MCMT for each genotype, and plotted reaction norms for each genotype against this value to visualize where maximum growth occurred relative to the genotype’s home climate.

We estimated fitness changes under future values of MCMT using the norm of reaction generated from the model to predict the growth, mortality, and combined fitness of each genotype at its provenance under historic (1961-1980) and future values of MCMT. Future MCMT values were predicted from an ensemble containing 13 general circulation models (GCMs) from the CMIP6 database, generated from ClimateNA (Wang et al. 2016) and available at AdaptWest (AdaptWest Project 2022). We used the time period of 2041-2070 under the shared socioeconomic pathway (SSP) 2-4.5, an intermediate scenario in which emissions rise until mid-century, then decline (IPCC 2023). To predict how the relative fitness of *P. balsamifera*, *P. trichocarpa*, and hybrid genotypes may change under future values of MCMT, we predicted which genotypes would have the best performance (highest overall fitness combining growth and survival) across the species ranges and regions with common gardens. For each grid cell, we extracted MCMT values for past and future climates, and identified the genotype with the highest predicted fitness at that temperature.

## Results

### Variation in growth among gardens

Growth increment varied widely among gardens (Figure S2), with the highest growth occurring at WSU (Washington) and SU (Maryland) gardens, and the lowest growth occurring at the upper and lower winter temperature extremes, particularly at OLLU (Texas), UCM (California), OSU (Oregon), and NDSU (North Dakota). Genotypes typically reached their greatest height gain at intermediate temperatures warmer than their climate of origin, with growth increment decreasing at temperature extremes (**Figure S3** and **S4**), suggesting that MCMT exerts selection pressure on poplars, as previously found in high-latitude tree species (Leites et al. 2012, 2019; Yeaman et al. 2016; Rehfeldt et al. 2018; Mahony et al. 2020).

### Factors predicting growth and mortality

We selected MCMT as the climate variable used in each model, because it had the lowest AIC score when predicting growth increment for 2021 (Table S2), and MCMT values in gardens included MCMT values from most historic home environments. We evaluated the effect size and significance of each model factor considering the two model components: the conditional component, representing growth, and the zero-inflated component, representing the probability of mortality (**Figure 2** and **Table S3**). Here we report effect sizes as standardized beta coefficients to enable comparison of the relative importance of each factor. Intercepts of random effects are presented in Figure S5, and effect sizes are in Table S4. The two model components had similar effects, indicating that the same factors associated with increased growth were also associated with low mortality. MCMT of the common garden site and its square term had the greatest effect on growth (slope = −0.53 and −0.46, p < 0.001 and = 0.001 respectively, **Figure 2** and **Table S2**). Both had negative effects, indicating decreased growth in climates warmer or colder than the optimum temperature. Intra- and interspecific genetic structure represented by PC1 and PC3 both had significant effects on growth, although with a smaller effect size than garden temperature (slope = −0.3, p < 0.001 for PC1 and slope=-0.15, p=0.003 for PC3, Tables S2). Individuals with greater *P. trichocarpa* ancestry (PC1) and from the three southernmost transects (PC3) had higher growth on average. PC2, which explains genetic structure within *P. trichocarpa* (**Figure S1**), was not statistically significant, suggesting that genetic differences between interior and coastal *P. trichocarpa* did not significantly affect growth. Home temperature and its interaction with garden temperature did not significantly affect growth, suggesting that responses to MCMT were better explained by genetic structure than by MCMT of origin. While no genotype × environment interaction terms were significant at the p<0.05 level across both years, Garden MCMT^2^ × Genetic PC2 was significant for a model including only measurements from 2021 (**Figure S6**), suggesting a weak effect of variation in the temperature response curve between two *P. trichocarpa* lineages during first year growth (**Figure S1**). Overall, our results show that, of the factors tested, temperature effects had the largest effect on growth. Genetics also contributed to variation in growth, which suggests the response to temperature is partially determined by species ancestry and latitudinal variation within both species. Similarly, in the zero-inflated model, garden MCMT significantly affected the probability of mortality, with increased mortality in sites at the higher and lower temperature extremes. Genetic structure, home MCMT, and their interactions with garden MCMT were not significant, suggesting that among the tested factors, mortality probability was primarily driven by planting site temperature, with limited variation among genotypes.

**Figure 2.**
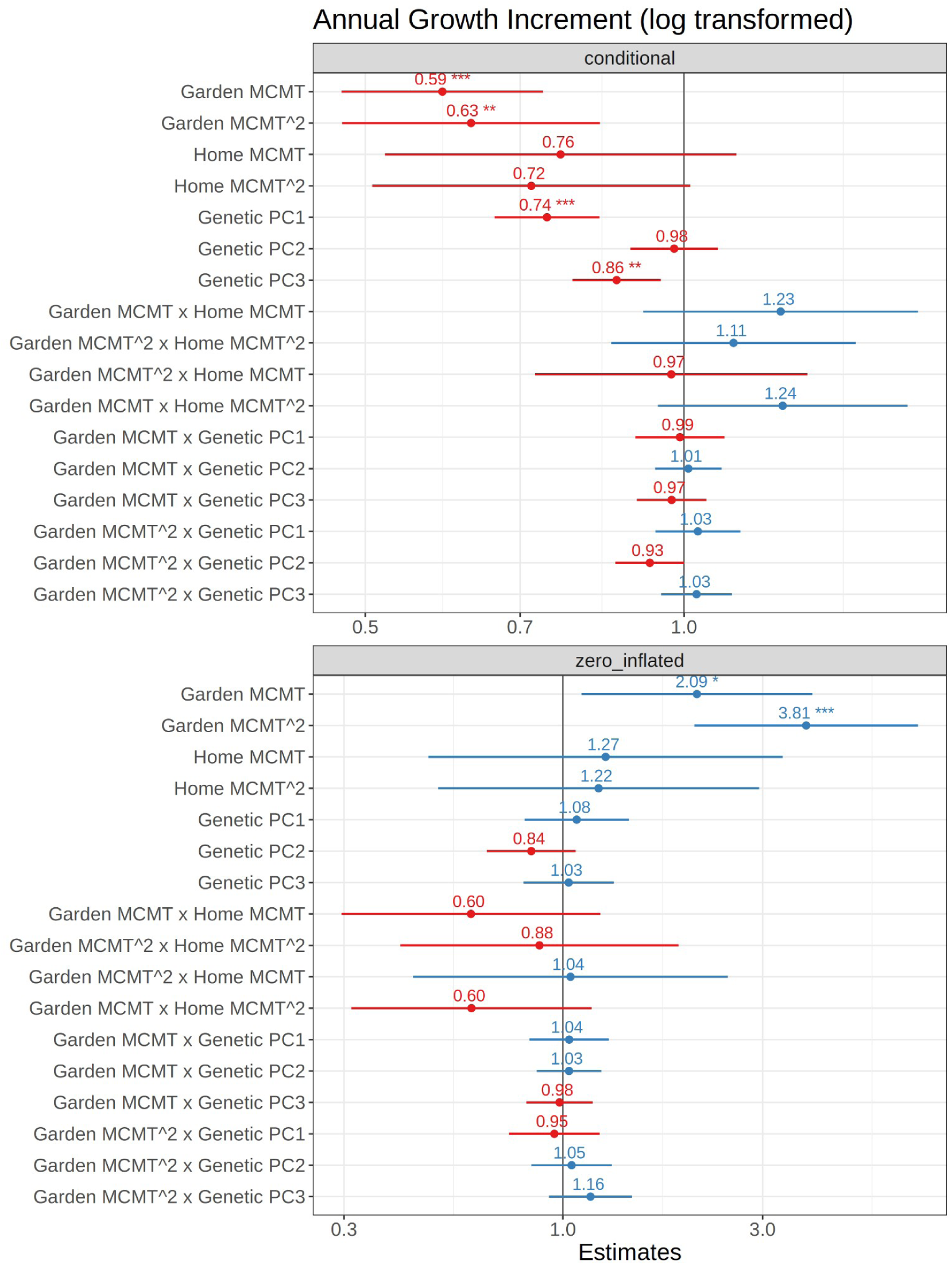
Forest plot of model coefficients using sjPlot (Lüdecke 2024); with an estimate of one indicating no effect, and estimates significantly greater or less than one indicating a positive and negative effect, respectively. Coefficients are standardized to allow comparison when interaction effects are included. Top plot shows results for the conditional model (growth increment) and bottom plot shows the estimates for the zero-inflated model (mortality); the effects for the two models are reversed because the zero-inflated model predicts the probability of mortality, which is negatively correlated with growth. P-values are indicated as follows: *<0.05, **<0.01, ***<0.001.

### Evaluation of model predictive ability

Both conditional (growth) and overall (growth and survival) components of Model 1 generally performed well in predicting the response to climate. The correlation between actual and predicted growth increments when random effects were included in the prediction was 0.795 for the overall model, and 0.809 for the conditional component (**Figure S7**). Similarly, the prediction of mortality from the zero-inflated component was significantly associated with actual mortality (P<0.001, Figure S8). When random effect estimates from Model 1 were not incorporated, predictive ability decreased to 0.631 and 0.538 for the overall and conditional components, respectively (**Figure S7**), indicating there were garden-specific effects on plant response not accounted for by garden MCMT, and/or genotype-specific effects that were not accounted for by home MCMT or the genetic PCs, and year-specific effects. However, the model consistently underpredicted actual growth increment values, particularly for the tallest trees. For this reason we focus on the relative differences in growth among genotypes and gardens rather than predicting specific yearly growth increments.

Prediction ability (Pearson’s correlation between observed and predicted growth increments) evaluated using leave-one-out cross-validation varied widely among gardens (**Figure S9**), with the average being 0.49 for 2021 and 0.38 for 2022, the highest correlation being 0.67 for OSU in 2022 (Oregon) and the lowest correlation being −0.8 for NDSU in 2022 (North Dakota). Most sites with poor predictions were sites with high mortality (NDSU, OLLU, LOCK, UCM, and OSU had >40% mortality in the first year). Other sites with low prediction ability included SWMN (r=0.058 in 2021 but increasing to 0.47 in 2022) and WI (r=0.397 in 2021, −0.092 in 2022), which are two sites that have low MCMT (**Figure S2**) and therefore may provide unique information on responses to lower temperatures. The relationship between predicted probability of mortality and actual mortality was not significant for most gardens and years (**Figure S10**), reflecting a low predictive ability for mortality alone.

We also predicted growth for our three ‘maxi’ common garden experiments, which included 500 novel genotypes and one novel site (**Figure 3**). Prediction ability was low for NDSU, likely due to high mortality (Pearson’s r=0.04 for the conditional model). However, prediction ability for the VA and VT modelwas relatively high. For the conditional model thecorrelation between actual and predicted values for VA was 0.45 and VT 0.65 (**Figure 3**), and when dead individuals were included in the overall model VA had an prediction ability of 0.48 and VT 0.53 (**Figure S11**). For VA, the predicted probability of mortality was significantly associated with actual mortality (Figure S12). However, as with the leave-one-garden-out tests, the model underpredicted growth. Taken together, these results suggest that for environments with low mortality, the model can predict relative performance for both novel genotypes and novel environments outside of the training dataset, but is limited in its prediction of specific growth increments.

**Figure 3.**
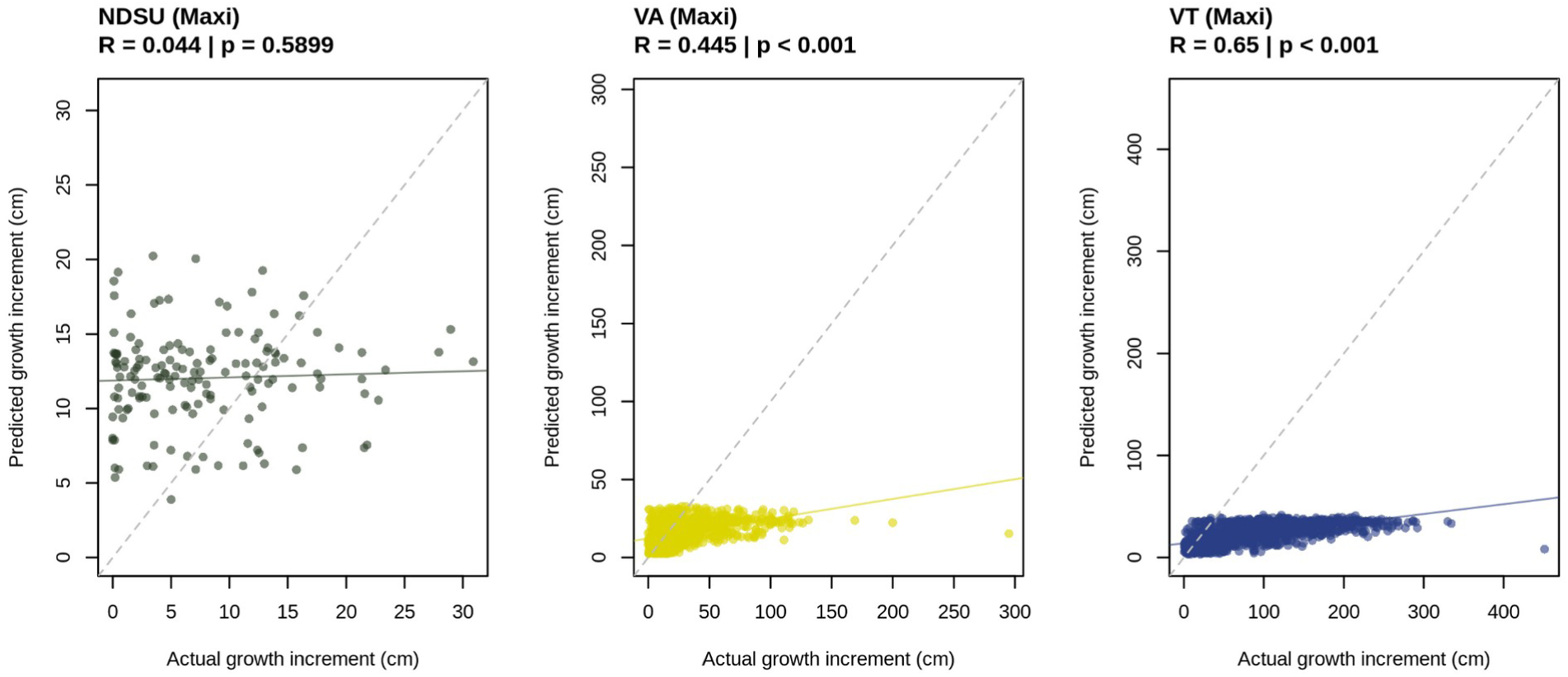
Performance of model predictions in the maxi gardens, predicting growth across 544 genotypes (500 novel genotypes) and one novel environment (VT) using a model trained on all 17 mini gardens. Models were evaluated by comparing predicted and actual growth increment for the conditional model with no random effects for garden and genotype. As the conditional portion of the model predicts growth rather than mortality, individuals that did not survive were removed from the model. R values and p-values are given for the Pearson correlation between actual and predicted growth. The dotted grey line indicates the 1:1 line; and the solid colored line indicates the best fit.

### The role of genetics in phenotypic prediction across environments

When predicting growth and overall performance for each garden and year using leave-one-out cross validation, the climate-only model (Model 3) that excluded genetic information had lower performance than the genetics-only model (Model 2; ANOVA of predictive ability from 17 leave-one-out cross-validation tests, p=0.0039). On average Model 3 had lower performance than the full model (Model 1), but the two were not significantly different (p = 0.061) (**Table 1**, **Figure 4**). These results show that genetic information improved predictions when climate data was excluded, but that models excluding either genetics or climate data performed similarly to the full model. When random effects of garden and genotype were included in the predictions, all models performed equally well (**Figure S13**). This is likely because all genotypes were included in the training datasets allowing the intercept of each genotype to be estimated, improving predictions of their growth in a novel environment without explicitly including genetic or climate information.

**Figure 4.**
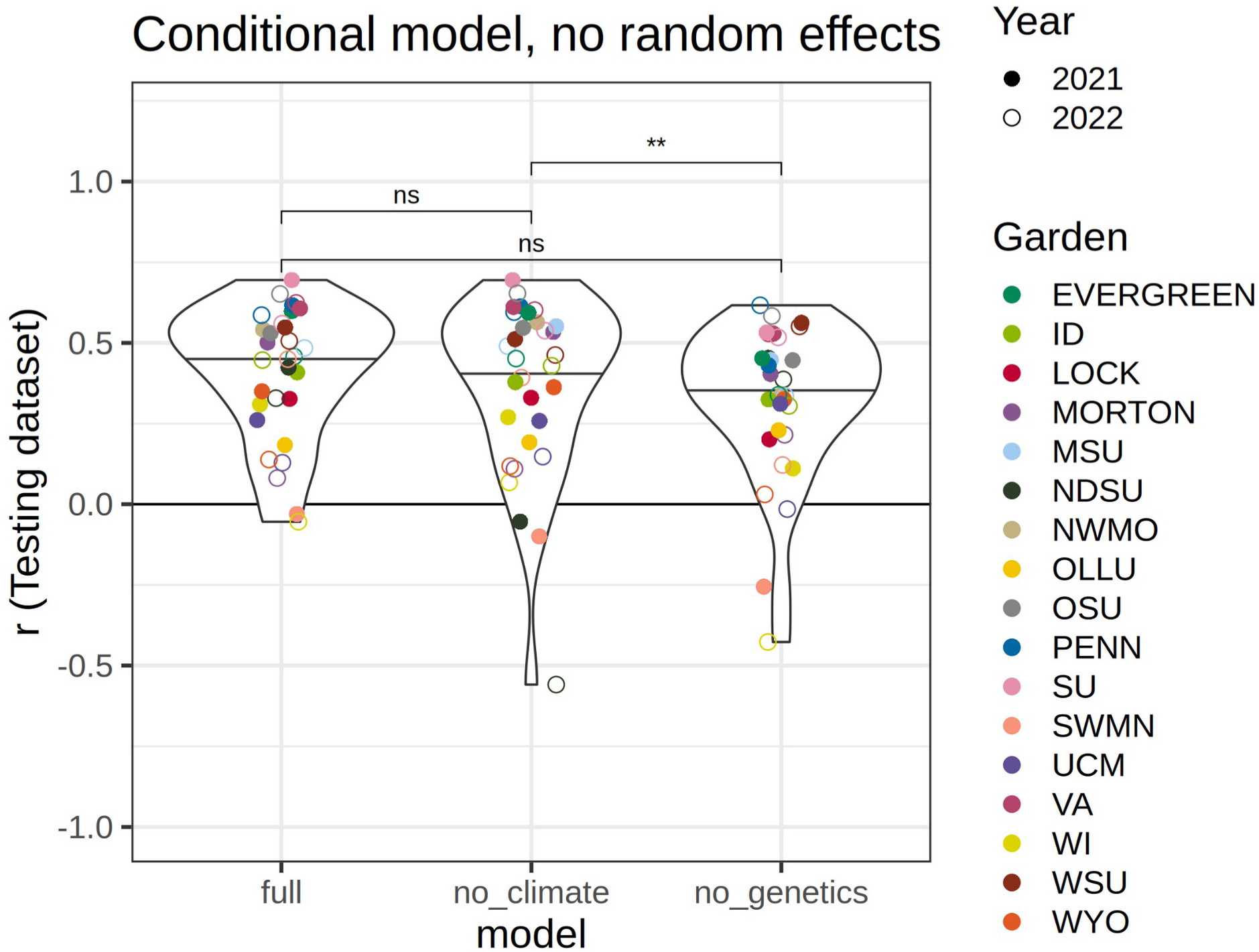
Comparison of model performance among the full model and those excluding provenance MCMT or genetic PCs (Table 1), estimated as the correlation between predicted and observed growth increments for each garden and year predicted using the leave-one-out models, in which growth increments were predicted for a single year and garden using a model trained on other gardens. The model including genetic data without home temperature data resulted in better predictive ability than the model with temperature data alone. Other comparisons with the overall model and with random effects included are shown in Figure S13. ** indicates p ≤0.01, “ns” indicates not significant.

The full model (Model 1) trained on all 17 gardens predicts that genotypes with more *P. trichocarpa* ancestry have higher growth and overall performance in warmer environments (MCMT ≈ −14 to 10 °C) compared to *P. balsamifera* genotypes, but that genotypes with more *P. balsamifera* ancestry have higher growth in colder climates, consistent with the climatic preferences of the two species (**Figure 5A-B**). Biologically unrealistic reaction norms were predicted for two majority-*P. balsamifera* genotypes, with growth increasing exponentially with decreased winter temperatures (**Figure 5B**).

**Figure 5.**
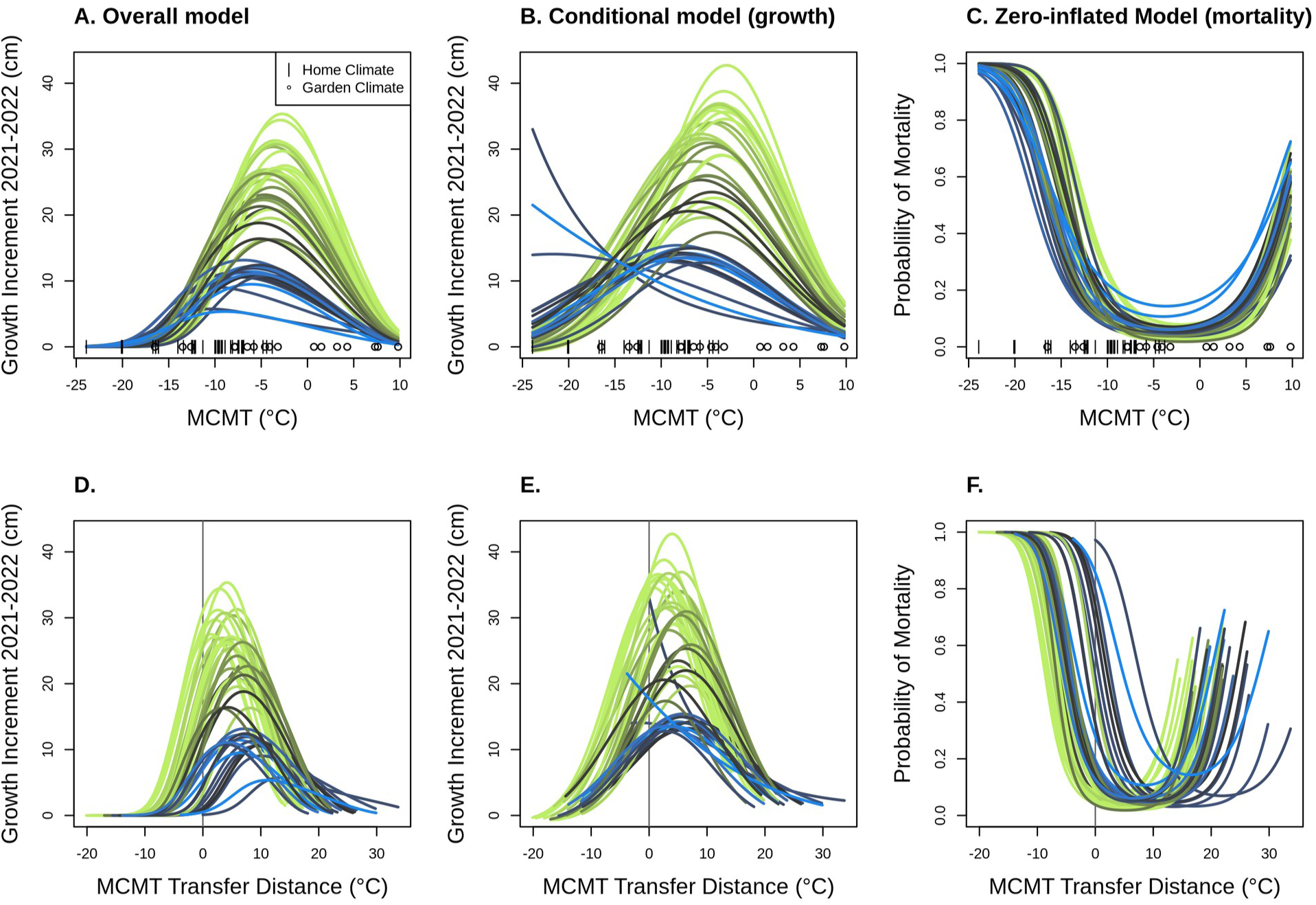
Model predictions for each genotype’s growth and mortality response to temperature, with each genotype indicated as a separate line colored by species ancestry at K=2, with green indicating *P. trichocarpa* and blue indicating *P. balsamifera*. Predictions for the overall model (A and D) incorporate both growth and the probability of mortality, predictions for the conditional model (B and E) only predict growth, ignoring the probability of zeros arising from other processes (mortality), and predictions for the zero-inflated model (C and F) predict the probability of zeros arising from mortality. A-C: Responses across the range of mean coldest month temperature (MCMT) values at garden and home climates. Actual values of home climates (|) and garden climates (circle) used for model training are shown on the x-axis. D-E. Responses based on distance from climate of origin (garden - home MCMT); positive values indicate a warmer climate (higher coldest month temperature) and negative values indicate a colder climate. If genotypes perform best in environments similar to their home environment, growth should be highest and mortality should be lowest when the transfer distance is 0.

These modeled growth responses likely reflect a preference for very cold environments that are not well-represented in our gardens, and when the effects of growth and mortality are combined, these genotypes show realistic response curves with optima at relatively low temperatures. Across species ancestries, mortality is predicted to increase at the colder and warmer extremes, with a greater probability of mortality under extreme cold climates compared to warmer climates (**Figure 5C**). As with predictions of growth, mortality responses predict that *P. balsamifera* is more cold tolerant, with increased probability of mortality in *P. trichocarpa* genotypes occurring at less extreme cold temperatures than for *P. balsamifera* (**Figure 5C**). Likewise, the probability of mortality for *P. balsamifera* generally increases with warmer temperatures more than *P. trichocarpa*. Combining the growth and mortality predictions as an overall fitness proxy, genotypes with a majority *P. trichocarpa* ancestry have consistently higher fitness except in the coldest climates, where majority-ancestry *P. balsamifera* genotypes are predicted to outperform them (**Figure 5A**). The overall maximum aboveground growth of *P. balsamifera* genotypes is lower than that of *P. trichocarpa*, which has previously been reported to be a faster-growing species (Larchevêque et al. 2011, Suarez-Gonzalez et al. 2017). Admixed genotypes with majority *P. trichocarpa* ancestry had maximum heights intermediate to the two parental species, while admixed majority-*P. balsamifera* genotypes had responses more similar to parental *P. balsamifera* genotypes.

### Rangewide projections of future fitness changes

Our full model (Model 1) predicts increased growth and survival under climates with MCMT temperatures warmer than the climate of origin (**Figure 5D-F** and **Figure S14**). However, we are limited in our ability to predict responses to multivariate changes in climate. Here, we focus on relative performance of genotypes rather than absolute changes in performance metrics. Predictions of the best-performing genotype under historic values of MCMT generally follows previously-described species ranges; with *P. trichocarpa* occurring along the west coast of North America and *P. balsamifera* occurring in the more northern and interior regions of the continent (**Figure 6 and S15**). Hybrid genotypes are predicted to outcompete parental species in intermediate regions of the contact zone, particularly in central British Columbia, where sampled genotypes exhibited high levels of admixture. Under future climates (2041-2070 for a “middle-of-the-road” emissions scenario), the model predicts shifts in the best-performing genotype following the two species’ climatic preferences: the regions suitable for *P. trichocarpa* and its backcrosses will shift northward as climate warms (**Figure 6**).

**Figure 6.**
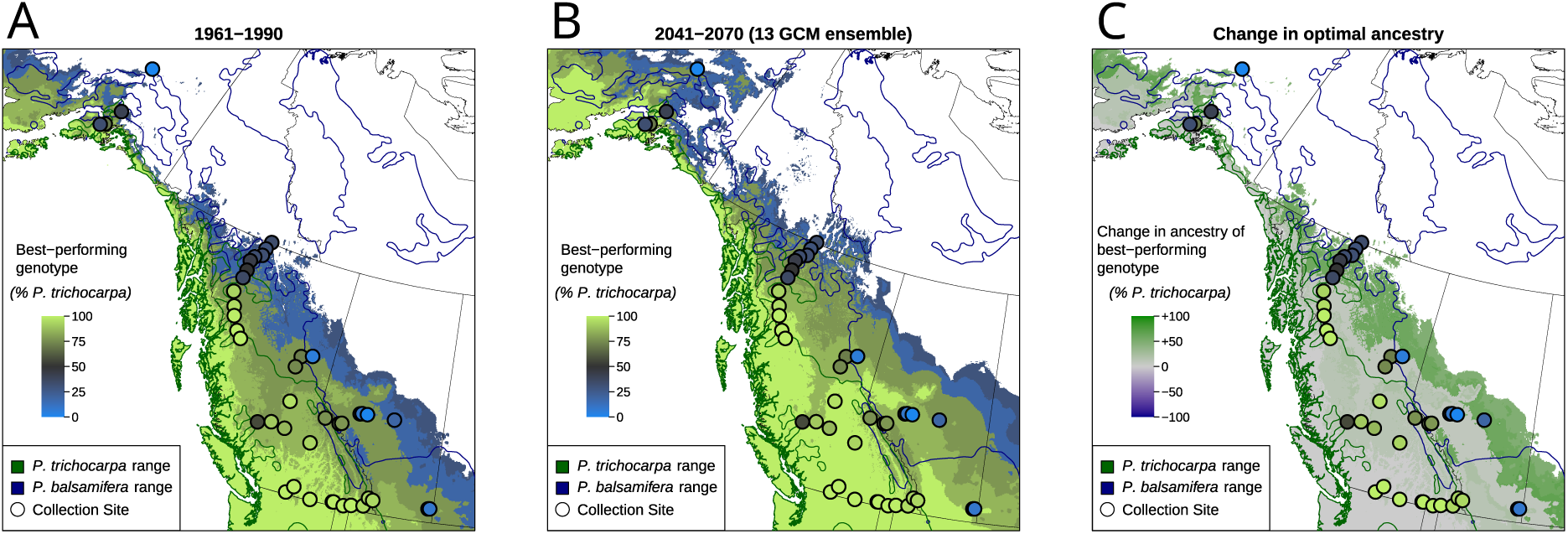
Maps showing the species ancestry of the studied genotype which is predicted to have highest fitness (as measured by growth and mortality) in that location under historic (A) and future climates (B) indicated by the color of the base layer, within the sampled hybrid zone region. Figure C indicates the change in optimal species ancestry between future (B) and historic (A) climates, indicating regions where increased *P. trichocarpa* ancestry is expected to be beneficial. Regions with MCMT values outside of the range measured in the common gardens (−13.05 to 10.85 °C) are masked and colored white. Actual ancestry of collected genotypes are shown as circles. Species ranges are shown as dark blue and green outlines (Little 1971). As MCMT increases, we predict that genotypes with higher *P. trichocarpa* ancestry may be able to outcompete genotypes with higher *P. balsamifera* ancestry in some portions of the *P. balsamifera range*, favoring a northeastern shift of the *P. trichocarpa* range and the hybrid zone and into historically colder, more continental regions. The same predictions are mapped across North America, including the common garden sites, in Figure S15.

Triangles and squares represent the climate at common garden sites for 2021 and 2022. Text indicates the loadings of climate and geographic variables on each axis, with abbreviations as follows: CMD, climatic moisture deficit; MAP, mean annual precipitation, MAT, mean annual temperature; MCMT, mean coldest month temperature; MWMT, mean warmest month temperature, PAS, precipitation as snow; RH, relative humidity; TD, temperature difference, or continentality. Loadings of climate variables have been downscaled to allow visibility of sites.

## Discussion

Using 17 common gardens with *Populus* genotypes which originated from a hybrid zone spanning a broad temperature range, we evaluated the genotype-specific response to warming winter temperatures and predicted future responses across the hybrid zone. We found that 1) fitness metrics and reaction norms varied among genotypes due to species ancestry and region of origin, consistent with a trade-off between cold tolerance and growth potential in warmer environments, 2) hybrids displayed reaction norms and temperature optima intermediate between their parental species, suggesting that warming temperatures could favor movement of the hybrid zone, and 3) a significant variance in growth explained by genetic structure illustrates the importance of genetic variation in predicting responses to climate change.

### Question 1: Do genotypes from different climates vary in fitness responses across environments?

Local adaptation to climate can shape not only traits, but their reaction norms across environments, contributing to variation in plastic responses within and across species (Patsiou et al. 2020). We found that poplar growth and survival across common garden environments was predicted by species ancestry and by genetic differences associated with a northern and southern region within each species (**Figure 2**), illustrating that genetic variation will contribute to spatially varying responses to warming temperatures (Patsiou et al. 2020; Martinez del Castillo et al. 2024). Performance traits varied along a continuous ancestry gradient between the two parental species, consistent with their climatic niches. In general, individuals with greater *P. trichocarpa* ancestry had greater overall growth, but these genotypes experienced lower growth and increased mortality in cold, continental environments typical of the *P. balsamifera* range (**Figure 5**). In addition, genotypes from the three southernmost transects had higher maximum yearly growth than those from the two northern transects (as described by genetic PC3). This is consistent with a tradeoff between growth potential and cold tolerance often observed in temperate or boreal tree species, likely enabling *P. balsamifera* to outcompete *P. trichocarpa* in colder regions (Leites et al. 2012; Menon et al. 2015; Rehfeldt et al. 2018). While no G×E terms were significant in the model based on two years of data, the effect of garden MCMT^2^ × genetic PC2 and was significant when only measurements from 2021 were included in the model (Figure S10). This result suggests a weak effect of genetic variation between two *P. trichocarpa* lineages on the plastic response to winter temperatures, which may be partially explained by maternal effects expressed in the first year of growth. Additionally, optimum temperatures varied for each genotype, with *P. trichocarpa* individuals reaching maximum growth in warmer environments than *P. balsamifera* **(Figure 5)**. This suggests that selection associated with minimum temperatures may be acting to produce different plastic responses across the range of both species and their hybrid zone.

While the majority of *P. balsamifera* genotypes had lower overall growth, their climatic range appeared to be wider (**Figure 5**), perhaps reflecting adaptation to more continental climates with larger annual temperature ranges (**Figure 1**). Generally, the response of hybrid genotypes was intermediate between the parental species; however, some hybrid genotypes have reduced probability of mortality at lower temperatures (**Figure 5**), suggesting that introgression could promote increased cold hardiness (Hamilton et al. 2013). However, the effects of species ancestry and geography may be difficult to disentangle, as genotypes from the coldest part of the sampled range (Alaska and northern British Columbia) are admixed (**Figure S1**).

In this study, we found that most genotypes reached their fitness optima in environments that were warmer than their climate of origin (**Figure 6**), and that genotypes planted at sites having MCMT values similar to their home site did not always perform better than other, nonlocal genotypes (**Figure S16**). Taken together, these results suggest genotypes may not be locally adapted to winter temperatures alone when considering either the “home vs away” or “local vs foreign” criteria (Kawecki and Ebert 2004). This decoupling between the physiological optimum, the climate in which genotypes reach their maximum growth rate, and the ecological optimum, the climate in which they exist in the ecosystem, has been previously observed for temperate tree species (Rehfeldt et al. 2018). Higher growth rates in non-local environments does not necessarily indicate a lack of local adaptation (Kawecki and Ebert 2004), but provides important context for predicting responses to climate change, which often assume optimal performance in local climates. Instead, we found that most genotypes, when in gardens with temperatures similar to their climate of origin, were outperformed by a small number of primarily *P. trichocarpa* genotypes. These genotypes may be ideal for planting as part of restoration projects using assisted gene flow or for biofuel production across diverse environments, particularly under increasingly variable and unpredictable climate futures (Sannigrahi et al. 2010; Porth and El-Kassaby 2015; Mahoney et al. 2019). However, if increased growth corresponds to a lack of cold tolerance or lower survival and fecundity (Leites et al. 2012, 2019; Yeaman et al. 2016; Rehfeldt et al. 2018; Mahony et al. 2020), caution is warranted when planting in locations susceptible to cold stress. Further work comparing the cold tolerance of these genotypes could determine the lower temperature limits where they are likely to be successful, while reciprocal transplants could be used to test whether local genotypes outperform nonlocal genotypes. Regardless of the mechanisms underlying genetic variation in fitness proxies, the large differences in responses to temperature illustrate the importance and benefit of measuring reaction norms across wide geographic regions within and among species.

Our predictions of increased growth under warming climates for these *Populus* genotypes raises the question: is active conservation and management of poplar in this region necessary, or should we prioritize species more vulnerable to increased temperatures? Predictions of increased growth should be treated with caution, as they only represent aboveground height and mortality in the first two years of growth and exclude reproductive fitness measures and belowground biomass (Fischer et al. 2007). Additionally, we only modeled growth over two growing seasons, in relatively warm climates and with irrigation during the first year, limiting the selective response to cold or drought stresses. In natural environments, major selective weather events may occur only rarely; for example, an unusually cold winter could favor slower growing but cold-tolerant genotypes over longer timescales than studied here (Rehfeldt et al. 2018; Lowry et al. 2019). In this case, our dataset may underestimate the benefit that *P. balsamifera* has in colder regions. Similarly, if warming winters are accompanied by summer heat waves, the benefits of warming we observe may not persist long-term in wild populations.

Precipitation levels and timing will also shift with climate change, and these changes can compound with temperature stresses (Arend et al., 2013; Gantois, 2022; Zandalinas & Mittler, 2022). However, we were unable to assess the effects of precipitation because most gardens were irrigated in the first year to allow seedling establishment. We also did not consider biotic factors such as competition with more cold-hardy conifers or pathogen pressure (e.g. *Melampsora* spp.) that may exclude *Populus* from warmer climates (La Mantia et al. 2013; McKown et al. 2014; Suarez-Gonzalez et al. 2018), interactions with belowground communities (Fischer et al. 2014b), or shifts in herbivore presence. If biotic interactions shift along with climate change, overall fitness could decrease even if higher temperatures favor faster aboveground growth. Lastly, our predictions of the response to climate are limited by the training data used to create the model – in this case, the climatic range of the common gardens (Rogers and Holland 2021). Because genotypes were largely planted in climates warmer than their climate of origin, most temperature variables (including multivariate PC axes) had little overlap between home and garden climates, limiting our ability to make predictions for growth and mortality in the colder historic climates of origin. We choose to use MCMT as the predictor variable because of its biological importance in past studies (Leites et al. 2012, 2019; Yeaman et al. 2016; Rehfeldt et al. 2018; Mahony et al. 2020) and because garden climates included temperatures matching historic home climates for both *P. balsamifera* and *P. trichocarpa* genotypes (Figure 1B-C, Figure 5). Given the limitation of our predictions for the coldest part of the hybrid zone and the potential effects of precipitation, testing how native poplar populations are responding to climate change *in situ* could complement common garden studies (Sharma et al. 2022; Astigarraga et al. 2024; Martinez del Castillo et al. 2024).

### Question 2: How will warming temperatures alter the geography of the two species and their hybrid zone?

Geographic variation in the magnitude of climate change and differences in phenotypic plasticity among genotypes could alter local competition dynamics among genotypes and lead to range shifts for the two species and their hybrid zone. While we predict that local poplar populations across the sampled region could experience increased growth and decreased mortality from warmer winter temperatures, they must contend with competition from non-local genotypes that may have greater fitness than local genotypes under novel climates. Species ranges are already beginning to shift in response to climate change (Parmesan 2006; Bell et al. 2014; Astigarraga et al. 2024) and multi-species hybrid zones may also shift (Taylor et al. 2014, 2015; Billerman et al. 2016; Hamilton and Miller 2016; Wielstra 2019; Alexander et al. 2022). In this poplar hybrid zone, regions where particular genomic backgrounds dominate may shift under climate change. For example, under higher minimum winter temperatures, *P. balsamifera* may be outcompeted by less cold-tolerant but faster growing genotypes with higher *P. trichocarpa* ancestry (**Figure 6**). However, such shifts in genetic composition would rely on gene flow or migration being fast enough to track climate change, either through dispersal of genotypes or seeds or wind-dispersed pollen (Corlett and Westcott 2013). While *Populus* can likely disperse over long distances via wind dispersal or river-dispersed vegetative material (Kling and Ackerly 2021), geographic barriers to gene flow persist across the hybrid zone, particularly in mountainous regions (Bolte et al. 2024). Our model predicts that parental *P. trichocarpa* genotypes may begin to outcompete *P. trichocarpa* backcrosses in central British Columbia, a region with high gene flow (Bolte et al. 2024), so this region may see shifts in the genetic composition of poplars (**Figure 6**). Conversely, while the regions where *P. trichocarpa* ancestry is favored are predicted to shift to the northeast in Alberta, the Rocky Mountains are a barrier to gene flow that may prevent their dispersal (Bolte et al. 2024). Our model also predicts that backcrossed *P. balsamifera* genotypes would perform better than parental *P. balsamifera* in northern, interior North America within the core of the species distribution (**Figure 6**). However, the reaction norms of parental and backcrossed *P. balsamifera* are very similar (**Figure 5A**) and because we have limited data for parental *P. balsamifera* genotypes and their growth in cold climates, we interpret these as regions where increased *P. balsamifera* ancestry is beneficial, not necessarily that backcrosses will be significantly more successful than parental genotypes.

Spatial changes in the location of the hybrid zone could provide an opportunity for adaptive introgression to occur, enabling individual alleles and novel allelic combinations to track climate change. Ancient and contemporary introgression can allow species to persist as environments change (Kremer and Hipp 2020; Leroy et al. 2020; O’Donnell et al. 2021; Buck et al. 2023; Yang et al. 2023). The presence of admixed genotypes in intermediate climates in this *Populus* hybrid zone and the reaction norms that are intermediate between the two parental species are consistent with the bounded hybrid superiority model, in which hybrids have higher fitness than either parental species in intermediate climates (Hamilton et al. 2013; De La Torre et al. 2014; Bolte et al. 2024). Movement of the *P. trichocarpa* x *P. balsamifera* hybrid zone could allow genes to move across species boundaries at new contact zones, increasing the levels of standing genetic variation in these regions and facilitating introgression of alleles that are beneficial under novel climates.

### Question 3: What information is needed to predict climate change responses in trees?

Like generations of provenance studies, we find that the temperature of the planting site had the greatest effect in determining growth, illustrating the continued value that common garden experiments will have in predicting organisms’ responses to climate change (O’Neill et al. 2008; Wang et al. 2010; Leites et al. 2012; Browne et al. 2019; Leites and Benito Garzón 2023; Ye et al. 2023).

Understanding how growth and survival varies across species, populations, and environments will be essential to predicting shifts in the performance and carbon sequestration abilities of trees under changing climates. We show that a model including genetic PCs but not climate of origin significantly improved predictions of growth compared to a model containing only home and garden climates (Figure 5), suggesting that universal transfer functions for predicting tree growth (O’Neill et al. 2008; Wang et al. 2010) could benefit from genetic information, as found by other studies (Mahony et al. 2020; Archambeau et al. 2022; Putra et al. 2023; Li et al. 2025). Furthermore, the model that included genetic information but not climate of origin predicted growth equally as well as the full model, suggesting genetic structure may be used to effectively model responses to climate in this system.

Genetic isolation-by-environment is stronger than isolation-by-distance in this system (Bolte et al. 2024), suggesting environmental adaptation drives much of the observed genetic structure.

Conversely, in species with high gene flow and less structured populations, adaptation to climate may be the primary factor contributing to intraspecific variation, and genetic structure may not be as effective in predicting responses.

## Conclusions

By characterizing genotype-specific reaction norms across 17 common garden environments, we were able to predict which genotypes should have the highest performance in historic and future climates across the hybrid zone, identifying regions where the contact zone is favored to shift under warming winter temperatures. If migration and gene flow is able to track climate change, a moving hybrid zone could facilitate adaptive introgression of alleles beneficial under novel climates, enabling poplar populations to persist in their ecosystems even as their genetic makeup may change.

## Supporting information

Rmarkdown output files

Common garden data (maxi gardens)

Common garden data (mini gardens)

## Acknowledgments

This research was supported by NSF PGR 1856450, the Schatz Center for Tree Molecular Genetics, National Institute of Food and Agriculture Award #PEN04809 to JAH, NIH award R35GM138300 to JRL, and the Life and Environmental Sciences Dept. at UCM. AM was supported by NSF PRFP 2209410. NM was supported by University of Wisconsin– Eau Claire Office of Research and Sponsored Programs through the Biology Research Scholars Program and Summer Research Experience for Undergraduate awards.

We would like to thank everyone involved in with sampling, garden establishment and maintenance, and data collection, including Shane Baumgart, Joseph Braasch, Schaefer Buchanan, Ian Clancy-Mallue, Emma Collins, Kevin Cowden, Lionel Di Santo, Marcia Fairbanks, Alice Fischer, Abby Ferson-Mitchell, Linnea Fraser, Carter Guffey, Carri LeRoy, David Hainlen, Alex Hass, Christian Hernandez, Kristina Hufford, Daria Hutchinson, Erika Jones, Lee Kalcsits, Ben Levitt, Eli Levitt, James Levitt, Jessica Lindstrom, Abigail Marshall, Luke McCormack, Albert Medel, Chloe Meyer, Katie Nelson, George Newcombe, Thu Nguyen, Amanda Penn, Jennifer Pollard, Steven Quick, Michelle Reid, Brendon Reidy, Christy Rollinson, Seth Townsend, Paul Warnick, Rebekah Wells, Dean Wu, Kate Volk, Lindsey Zakopal, and Thomas Zambiasi. We would also like to thank Tongli Wang for discussions of climate response functions.

## Author Contributions

JAH, JH, SRK, and MCF designed the study and contributed to data analysis and interpretation. AM performed analyses and wrote the first draft of the manuscript. AM, JRBB, ACB, DF, SF, JK, LK, SKK, MWK, JL, JML, DL, MN, EVM, JS, KLS, BWW and MZP contributed to fieldwork and data collection and provided input on garden-specific conditions for data analysis and interpretation. All authors contributed to writing the manuscript.

## Data Availability

Scripts for filtering genomic data are available at github.com/alaynamead/poplar_hybrid_vcf_filtering.

Analysis scripts are available at <u>github.com/alaynamead/popup_poplar_reaction_norms</u> and corresponding R markdown files with script outputs are provided as supplementary material. Github repositories will be archived at Zenodo upon manuscript acceptance.

Data from this study is included as supplementary material and will be archived in Dryad upon manuscript acceptance.

