## Supplemental Material for "Variation in responses to temperature in admixed *Populus* genotypes predicts geographic shifts in regions where hybrids are favored"

### Supplementary Methods

*Genomic data*

For each genotype, approximately 100 mg of young leaf tissue was used for genomic DNA extraction with the Qiagen Plant DNeasy kit. Genomic libraries were sequenced using an Illumina NovaSeq 6000, with 64 samples per lane, using paired-end 150 bp reads. Reads were aligned to the *P. trichocarpa* reference genome (v4.0) and variant calling was performed using GATK Haplotype Caller. Within GATK, variants were filtered for quality-by-depth (QD<2), mapping quality (MQ<40), elevated strand bias (FS>40, SOR>3), and differential map quality and positional bias between reference and alternate alleles (MQRankSum<-12.5, ReadPosRankSum<-8). Using bcftools, variants were subset to biallelic SNPs (-m2 -M2 snps) and variants with a minor allele count of 1 (--include ‘MAC>1’). VCFtools was used to remove all SNPs with missing data across individuals (max-missing 1.0). Finally, variants were LD-pruned in PLINK using a 10000 bp window size, shifted by 1000 bp, removing variants with a pairwise $R^2$ value >0.1. After filtering and LD-pruning, a total of 334,657 variable sites were used to characterize genetic variation and admixture.

*Model function*

We implemented Model 1 (main text) using the glmmTMB function from the glmmTMB package version 1.1.11 (Brooks et al. 2017) in R version 4.5.0 (R Core Team 2024) using the following formula. `log(growth increment + 1) ~ garden MCMT × home MCMT + garden MCMT`$^2$ `× home MCMT`$^2$ `+ garden MCMT`$^2$ `× home MCMT + garden MCMT × home MCMT`$^2$ `+ genetic PC1 × garden MCMT + genetic PC2 × garden MCMT + genetic PC3 × garden MCMT + genetic PC1 × garden MCMT`$^2$ `+ genetic PC2 × garden MCMT`$^2$ `+ genetic PC3 × garden MCMT`$^2$ `+ (1 | genotype) + (1 | garden/block) + (1 | year) + (1 | individual)`

We used a gaussian model for the conditional component (family = gaussian()) and used the same formula for the zero-inflated component of the model (ziformula = ~.). For complete code used, see the script “transfer_function_multiyear_linear_mixed_effects_model.Rmd.”

*Model evaluation*

We predicted heights for individuals based on the full model using the `predict` function from the glmmTMB package and calculated the Pearson correlation between the actual heights and the predicted heights. To quantify the model’s predictive ability for each year, we calculated the Pearson correlation between the actual heights and the predicted heights (conditional component ) as well as the predicted heights when the probability of mortality was included (overall model). To calculate predictive ability for the conditional component, we removed the individuals with a measured height of zero (which is accounted for in the zero-inflated component of the model). We also evaluated model predictive ability with and without the random effects of garden, block, and genotype. Including random effects (using option re.form=NULL in the predict function) accounts for the varying intercepts associated with each group, while excluding them and setting all random effects to zero (re.form = NA) makes predictions based on the fixed effects of climate and genetics only. When the model was used to predict growth in genotypes or gardens not included in the training set, a new random effect was predicted.

We also quantified the predictive ability for the zero-inflated model component estimating mortality rate by testing for a significant relationship between the probability of mortality predicted by the model, and the actual mortality, measured as a binary (yes/no) variable. We fit a generalized linear model with a binomial link function in R using the `glm` function. We evaluated the model fit using the p-value; however, $R^2$ values are not estimated for GLMs and are not reported here. Instead, we focus on the predictive ability of the overall model, which incorporates both the conditional and zero-inflated components of the model.

### Supplementary Figures

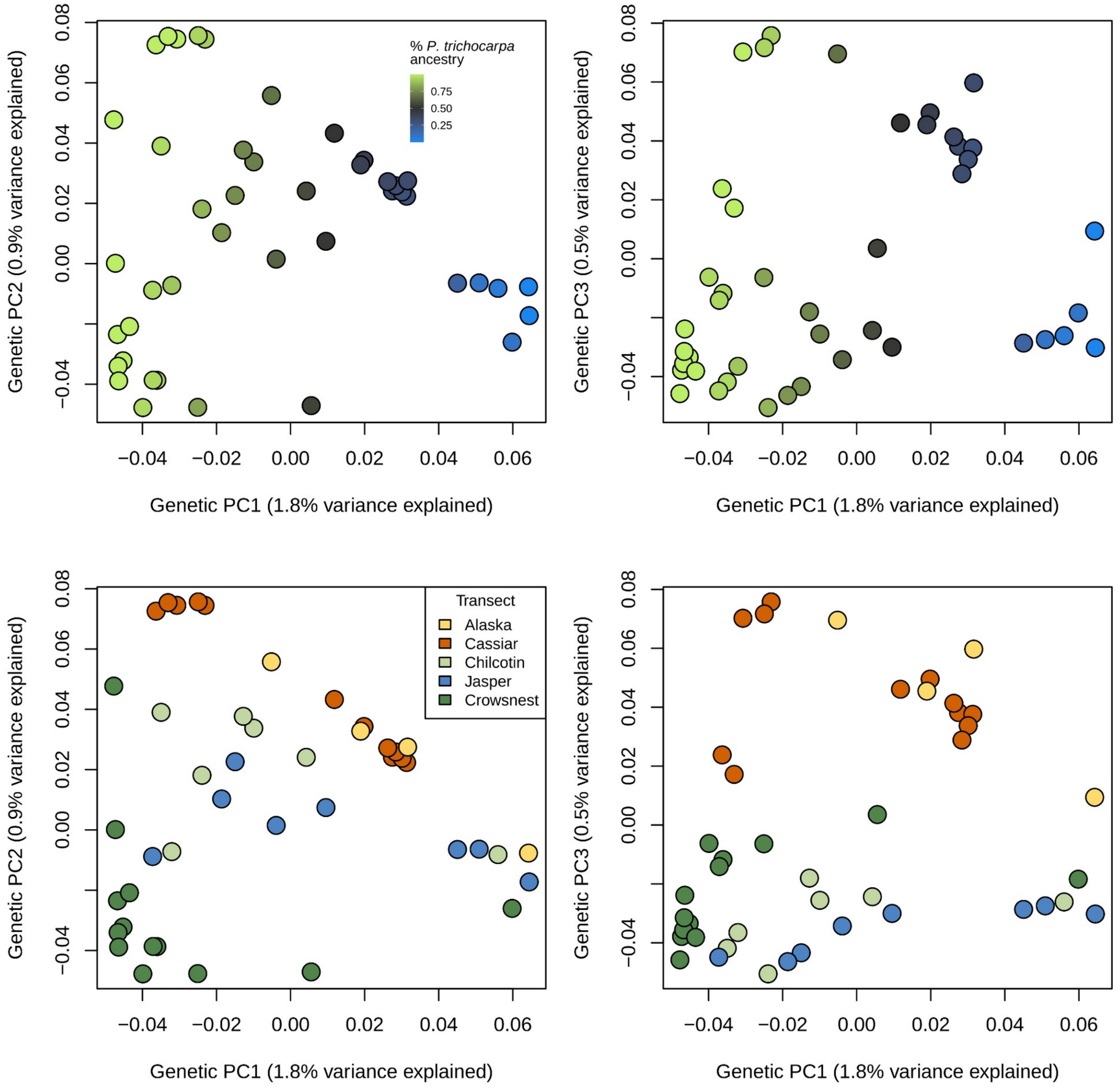


**Figure S1.** Genetic PCs 1-3 for each genotype, used to estimate the effect of genetic structure on phenotypic responses to climate. Top: colors represent species ancestry at K=2. Bottom: colors represent transect, listed from northernmost (Alaska) to southernmost (Crowsnest).

**Figure S2.**

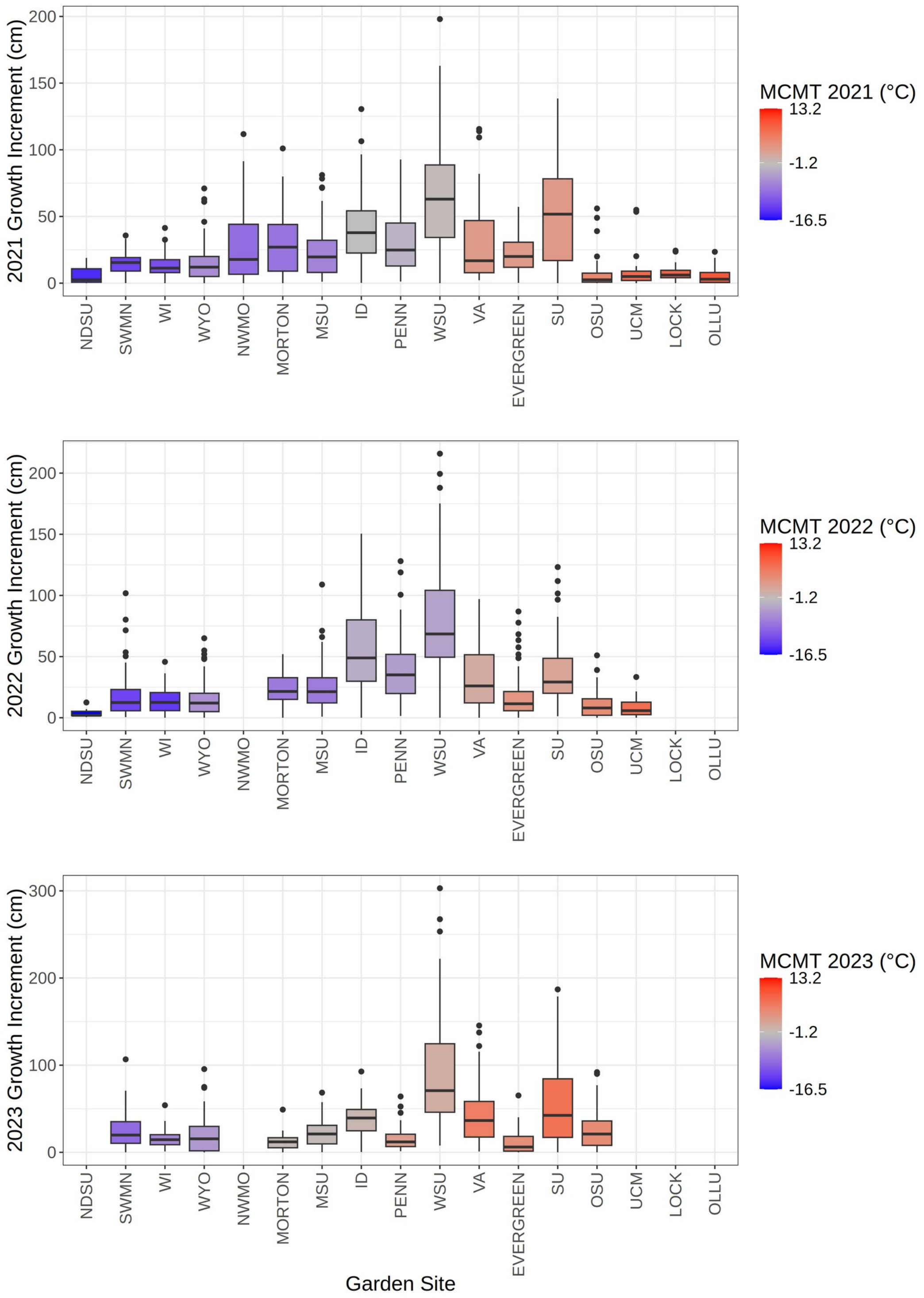


Variation in growth increment across gardens for each year. Gardens are ordered along the x-axis by their mean coldest month temperature (MCMT) averaged from 2020-2023. Garden boxplots are colored by the MCMT (°C) of that year.

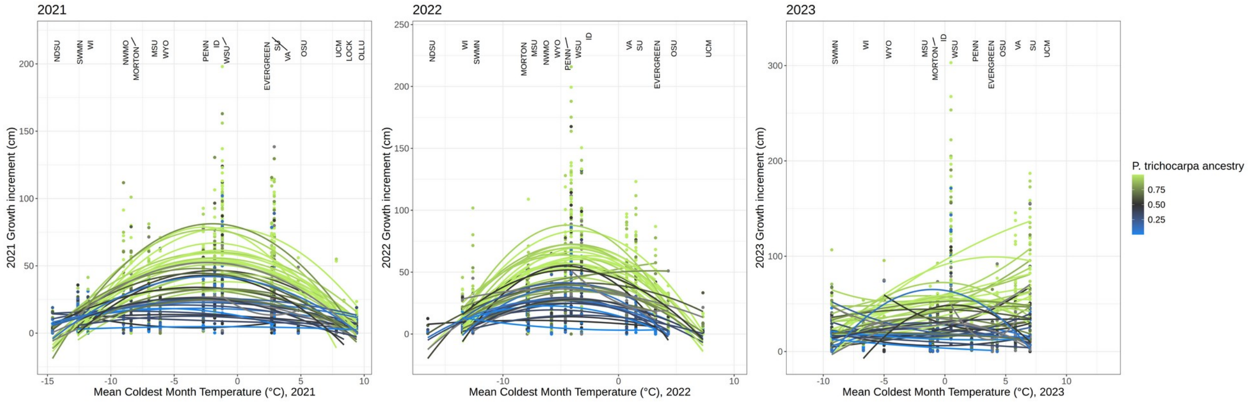


**Figure S3.** The genotype-specific response of yearly growth increment to garden MCMT for three separate years (2021-2023), fit using a quadratic model of garden MCMT. Text labels indicate the MCMT value of each garden for that year.

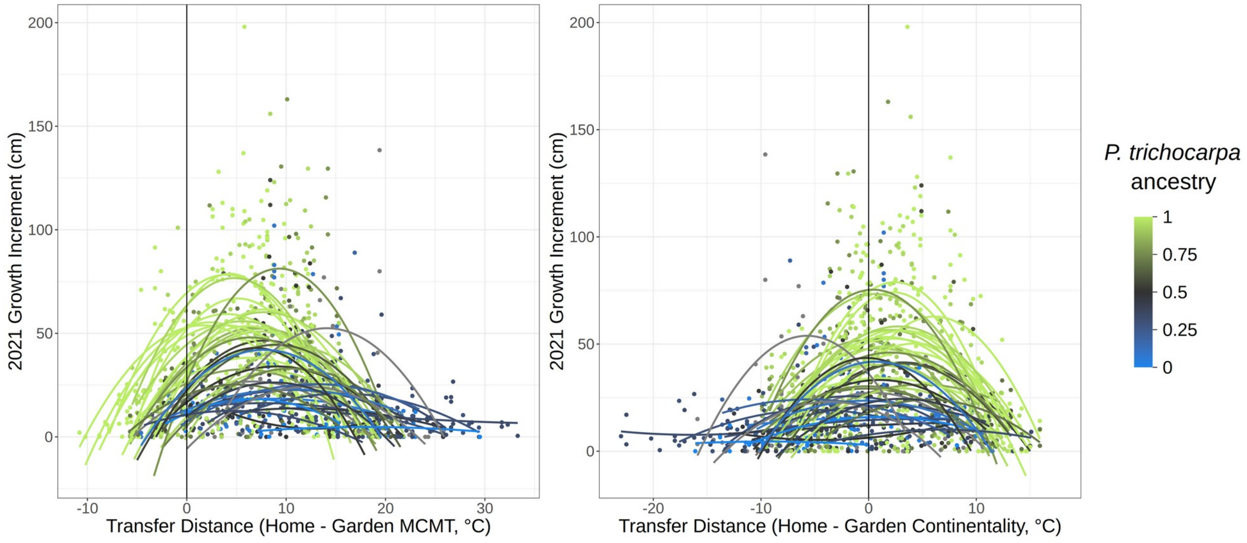


**Figure S4.** Response of growth increment to transfer distance, or the difference between home and common garden climate in 2021. Lines show the response of each genotype and are fitted using a simple quadratic model in the lm function in R. Line colors indicate the genotype's species ancestry at K=2. The response to continentality transfer distance indicates genotypes generally have the highest growth in environments similar to their climate of origin (transfer distance = 0) while the response to mean coldest month temperature transfer distance indicates genotypes have higher growth in environments that are warmer than their climate of origin.

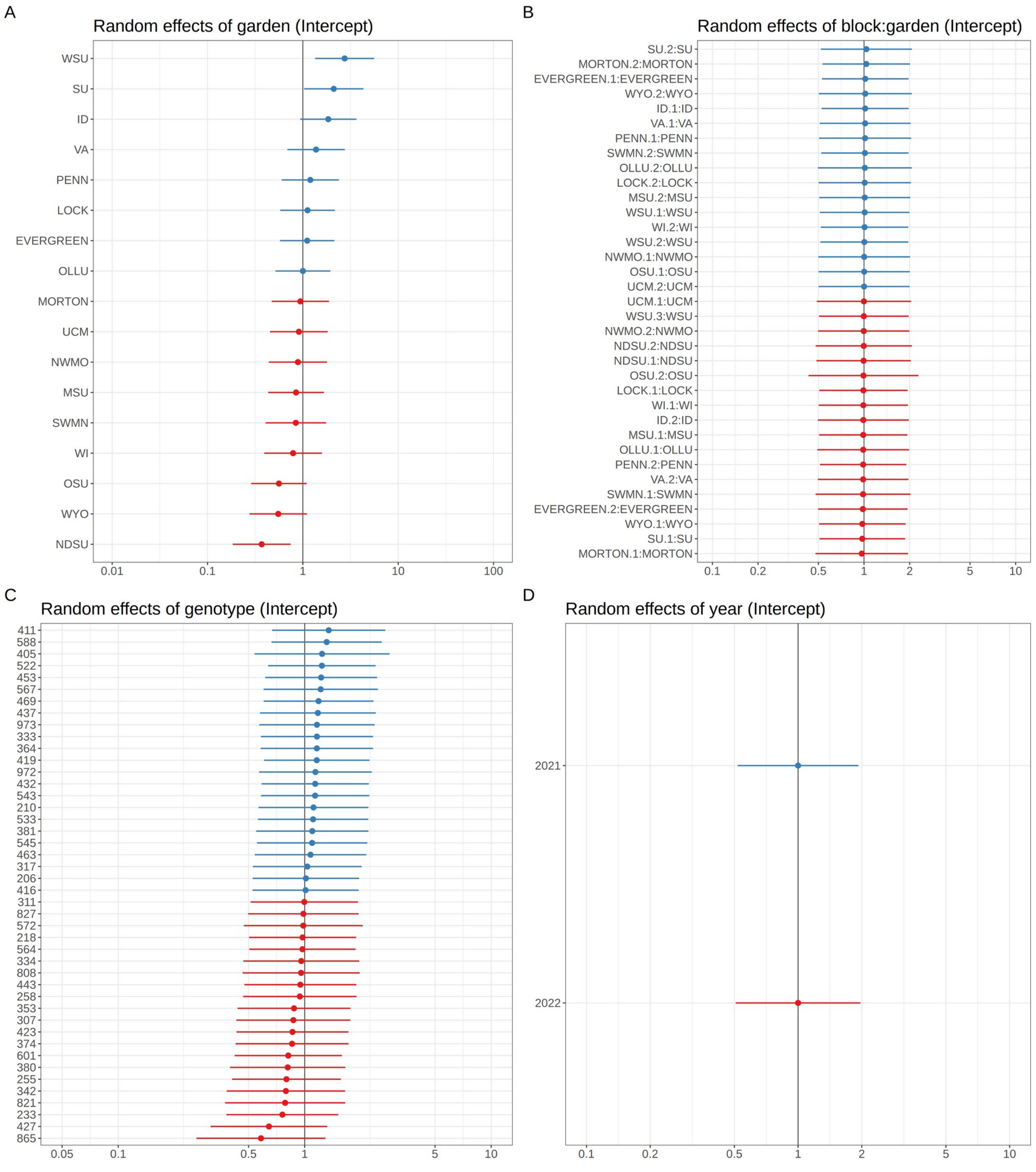


**Figure S5.** Estimates of model random effects for A) garden site, B) block nested within garden, C) genotype, and D) year.

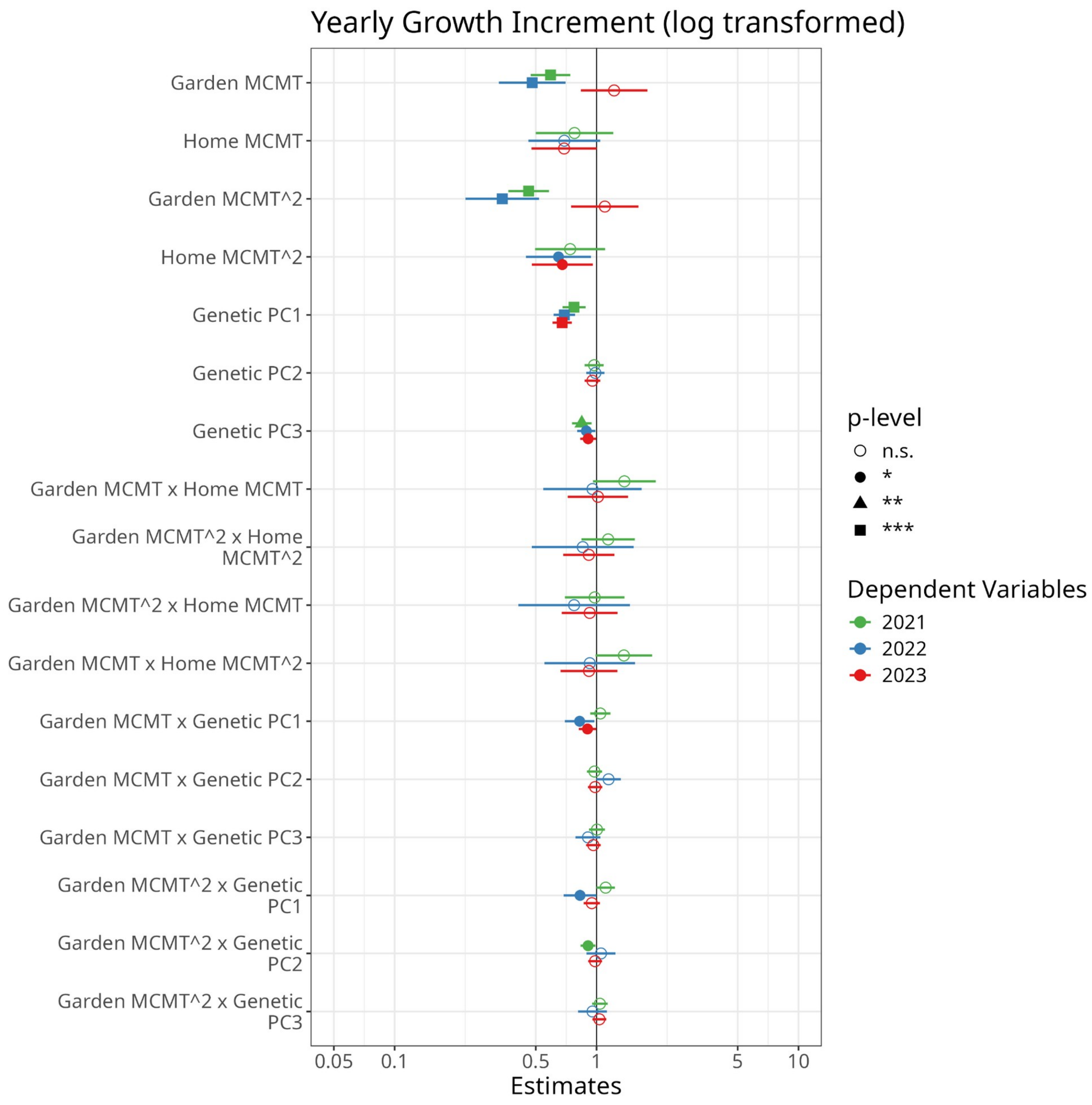


**Figure S6.** Comparison of model effects across three years of data collection, as in Figure 2. A model was fit to the subset of data from each year by modifying Model 1 to drop the random effects of year and individual, as there is one measurement per year per individual.

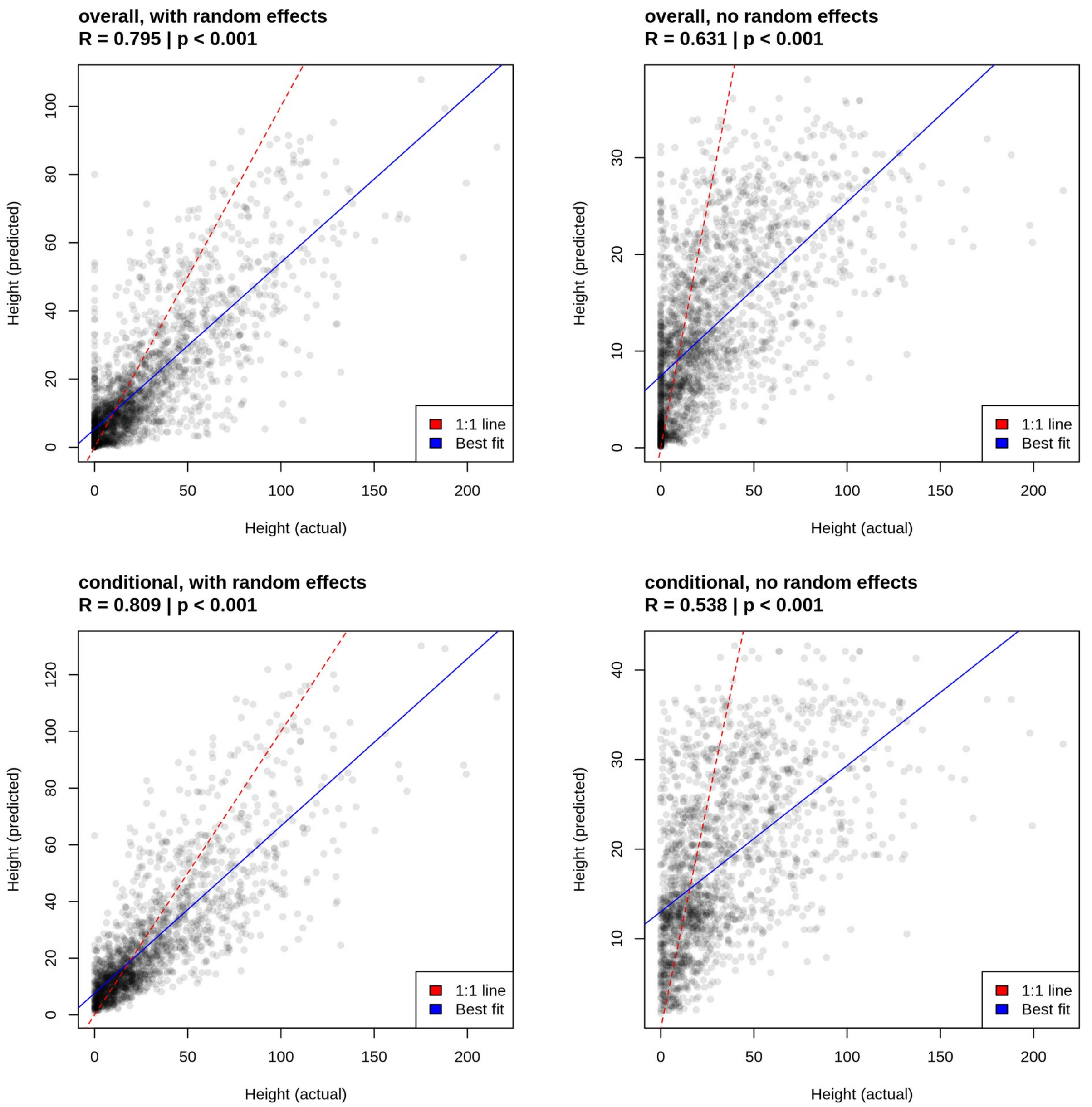


**Figure S7.** Correlation between actual and predicted growth for individuals across all 17 garden sites and two years, with the Pearson correlation and its p-value, across four categories of predictions from the model. Results are shown for heights predicted from the overall model, including growth and mortality, with dead individuals having a height of zero (top) and for the conditional model representing growth in the surviving individuals (bottom). When the random effects of genotype, garden, and block are included (left) predictions are more closely correlated with actual heights compared to when they were excluded (right). Best fit lines have a lower slope than the 1:1 line, indicating that the model underpredicts height for taller individuals.

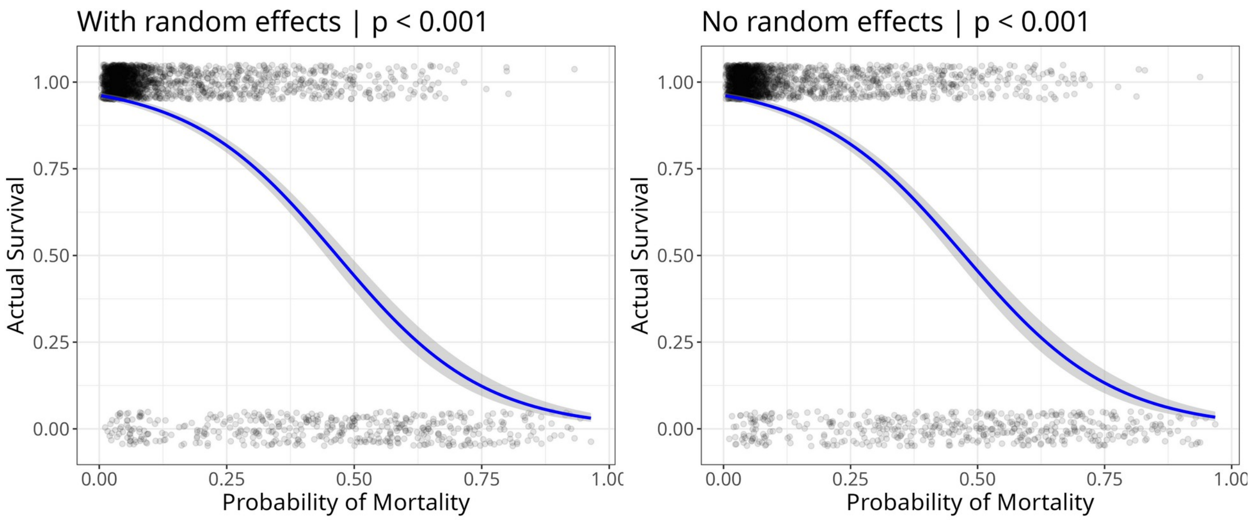


**Figure S8.** Relationship between actual survival and the probability of mortality predicted from the full model, including all individuals across 17 garden sites and two years. The logistic relationship was modeled using a generalized linear model with a binomial link function, and the p-values for each model are shown.

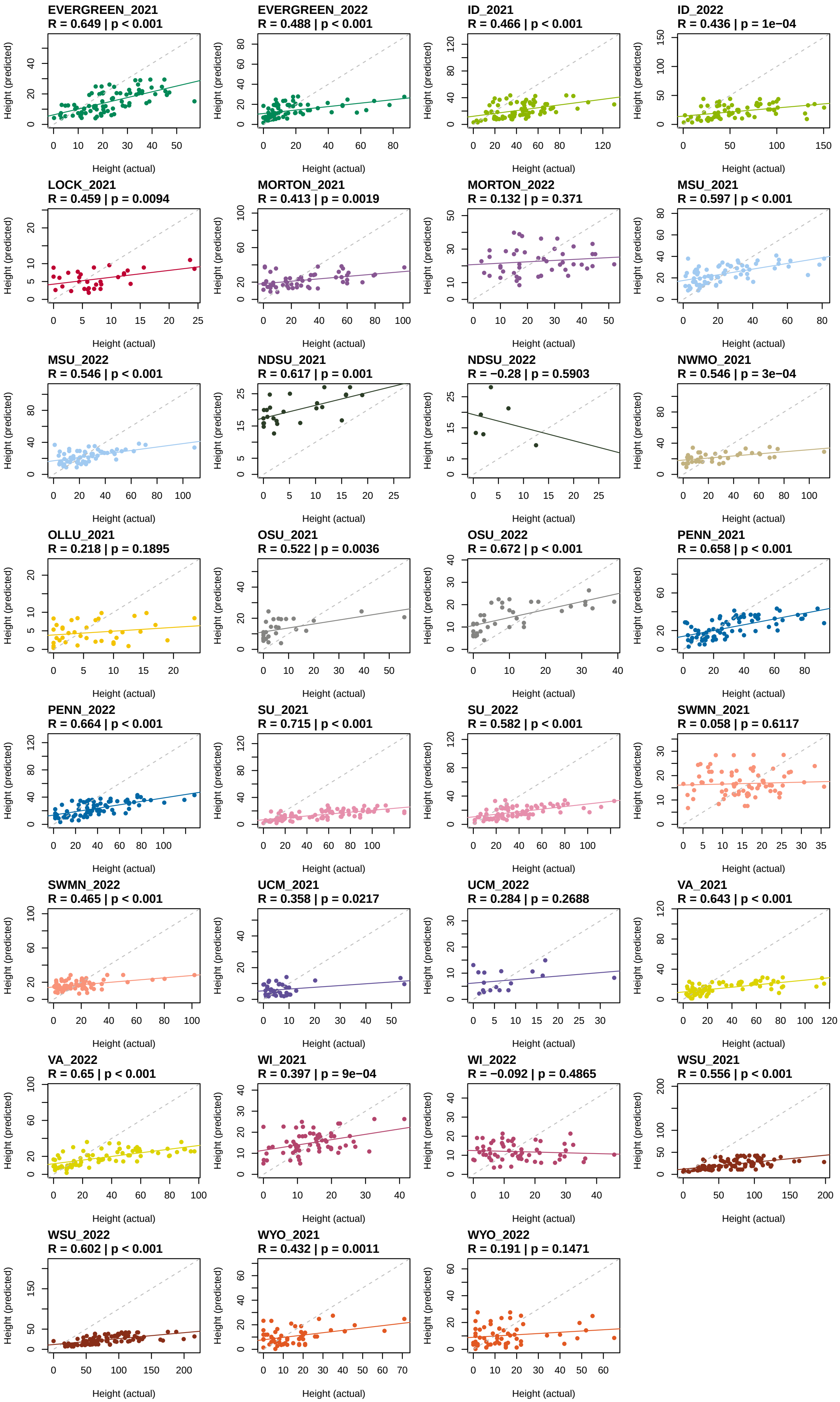
EVERGREEN_2021
R = 0.649 | p < 0.001
EVERGREEN_2022
R = 0.488 | p < 0.001
ID_2021
R = 0.466 | p < 0.001
ID_2022
R = 0.436 | p = 1e−04
LOCK_2021
R = 0.459 | p = 0.0094
MORTON_2021
R = 0.413 | p = 0.0019
MORTON_2022
R = 0.132 | p = 0.371
MSU_2021
R = 0.597 | p < 0.001
MSU_2022
R = 0.546 | p < 0.001
NDSU_2021
R = 0.617 | p = 0.001
NDSU_2022
R = −0.28 | p = 0.5903
NWMO_2021
R = 0.546 | p = 3e−04
OLLU_2021
R = 0.218 | p = 0.1895
OSU_2021
R = 0.522 | p = 0.0036
OSU_2022
R = 0.672 | p < 0.001
PENN_2021
R = 0.658 | p < 0.001
PENN_2022
R = 0.664 | p < 0.001
SU_2021
R = 0.715 | p < 0.001
SU_2022
R = 0.582 | p < 0.001
SWMN_2021
R = 0.058 | p = 0.6117
SWMN_2022
R = 0.465 | p < 0.001
UCM_2021
R = 0.358 | p = 0.0217
UCM_2022
R = 0.284 | p = 0.2688
VA_2021
R = 0.643 | p < 0.001
VA_2022
R = 0.65 | p < 0.001
WI_2021
R = 0.397 | p = 9e−04
WI_2022
R = −0.092 | p = 0.4865
WSU_2021
R = 0.556 | p < 0.001
WSU_2022
R = 0.602 | p < 0.001
WYO_2021
R = 0.432 | p = 0.0011
WYO_2022
R = 0.191 | p = 0.1471
Height (predicted)
Height (actual)

**Figure S9.** Correlation between actual and predicted growth for each garden and year from leave-one-out cross validation predictions in gardens, in which predictions were made for the garden using a model trained on the other gardens. Pearson R values and p-values for the correlation are shown. Solid lines indicate the best fit line; grey dotted lines indicate the one-to-one line.

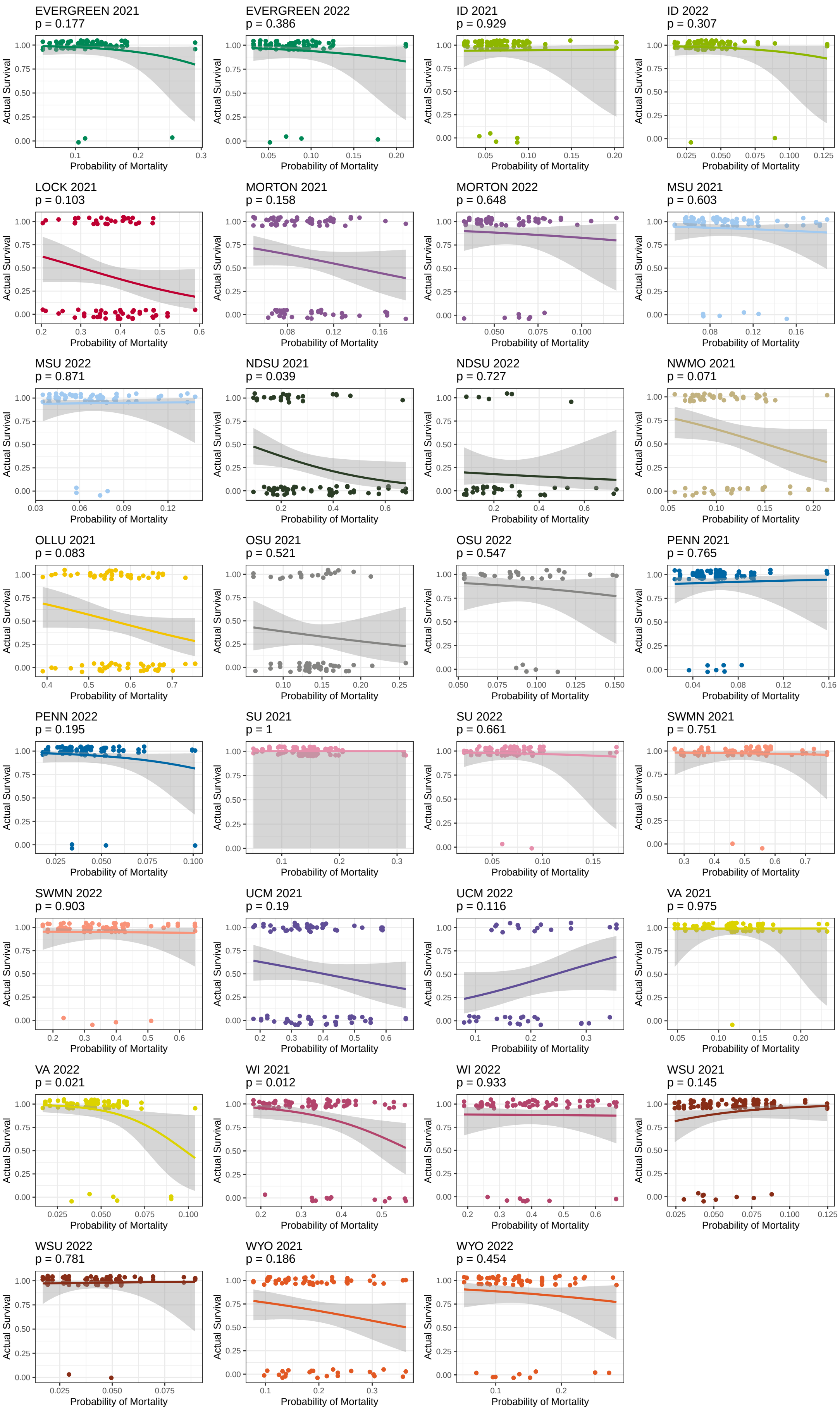
EVERGREEN 2021
p = 0.177
EVERGREEN 2022
p = 0.386
ID 2021
p = 0.929
ID 2022
p = 0.307
LOCK 2021
p = 0.103
MORTON 2021
p = 0.158
MORTON 2022
p = 0.648
MSU 2021
p = 0.603
MSU 2022
p = 0.871
NDSU 2021
p = 0.039
NDSU 2022
p = 0.727
NWMO 2021
p = 0.071
OLLU 2021
p = 0.083
OSU 2021
p = 0.521
OSU 2022
p = 0.547
PENN 2021
p = 0.765
PENN 2022
p = 0.195
SU 2021
p = 1
SU 2022
p = 0.661
SWMN 2021
p = 0.751
SWMN 2022
p = 0.903
UCM 2021
p = 0.19
UCM 2022
p = 0.116
VA 2021
p = 0.975
VA 2022
p = 0.021
WI 2021
p = 0.012
WI 2022
p = 0.933
WSU 2021
p = 0.145
WSU 2022
p = 0.781
WYO 2021
p = 0.186
WYO 2022
p = 0.454
Actual Survival
Probability of Mortality

**Figure S10.** Relationship between predicted probability of mortality and actual survival for each garden and year from leave-one-out cross validation predictions in gardens, in which predictions were made for the garden using a model trained on the other gardens. The logistic relationship was modeled using a generalized linear model with a binomial link function, and the p-values for each model are shown.

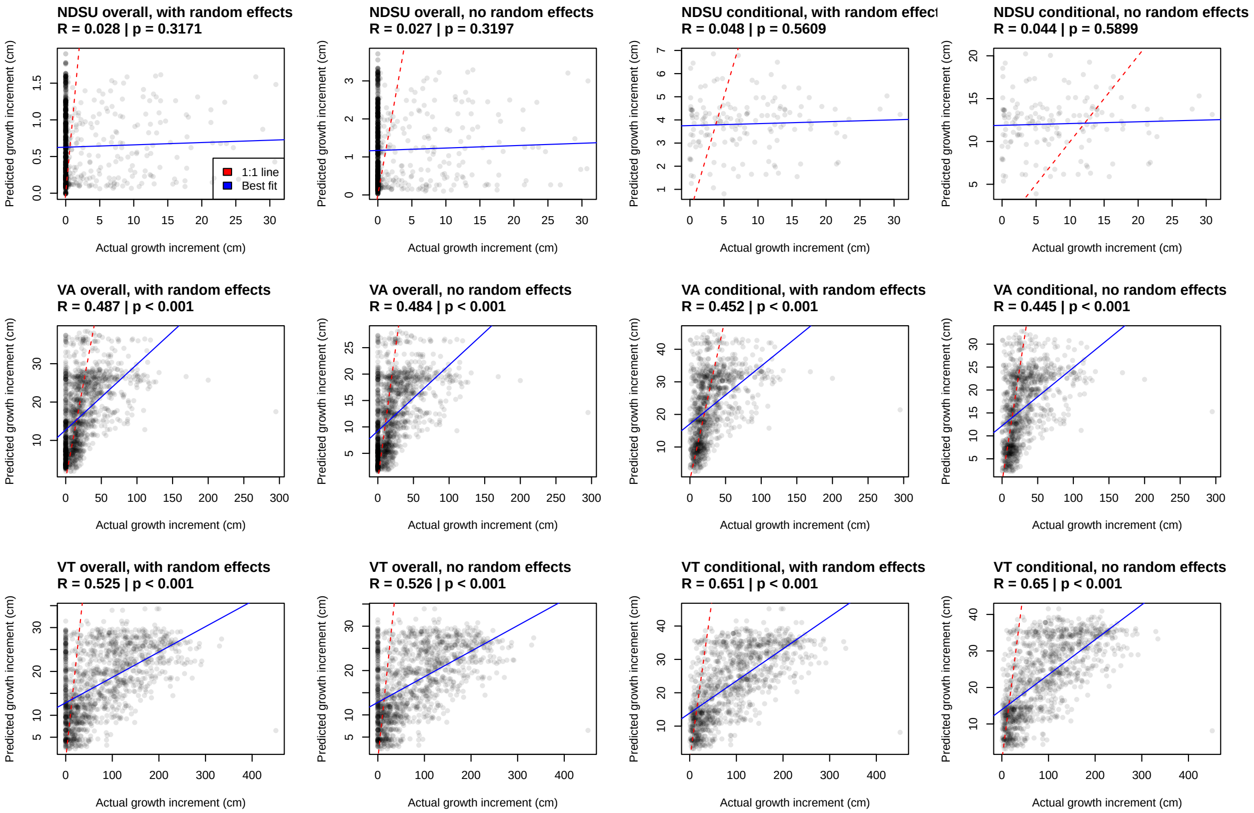


**Figure S11.** Correlation between actual and predicted growth for individuals for the three maxi garden sites, with the Pearson correlation and its p-value, across four categories of predictions from the model (as described in Figure S7).

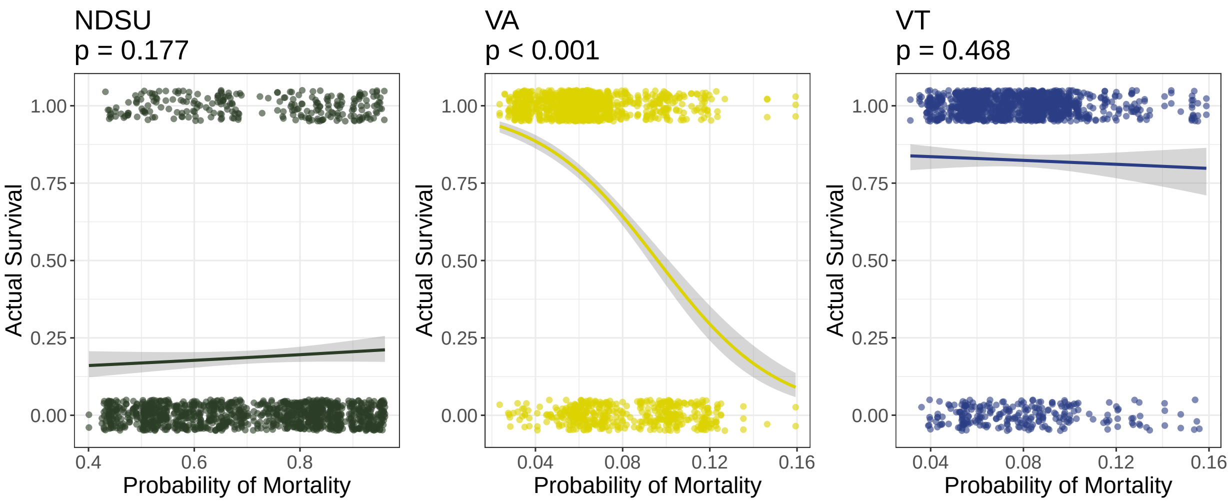


**Figure S12.** Relationship between predicted probability of mortality and actual survival for the three maxi garden sites. The logistic relationship was modeled using a generalized linear model with a binomial link function, and the p-values for each model are shown.

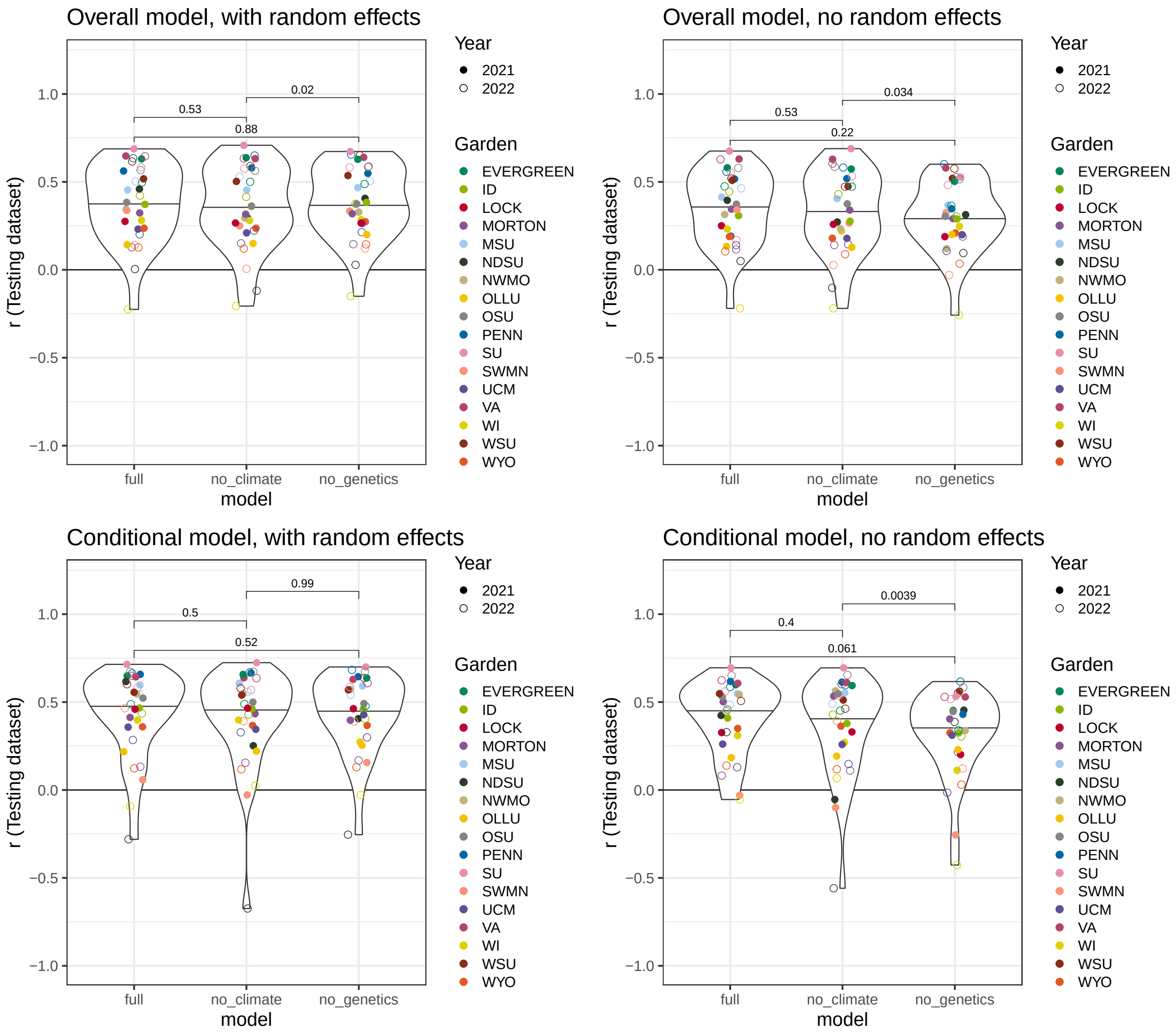


**Figure S13.** Comparisons of Pearson's correlation r values with and without genetic and home climate information. P-values are given for each pairwise model comparison using a paired Wilcoxon test.

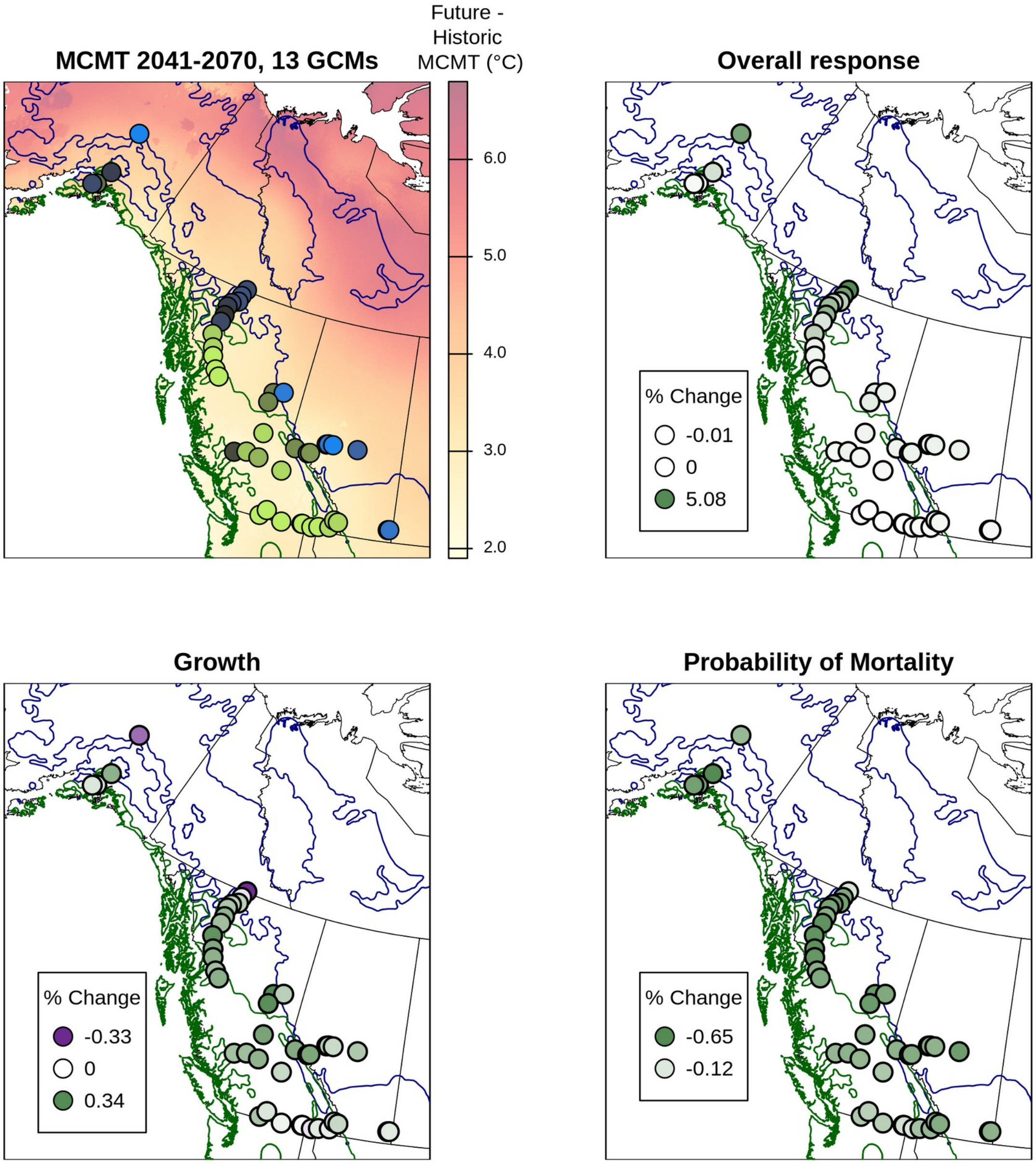


**Figure S14**. Spatial changes in MCMT and predicted changes in fitness metrics for each genotype as a result of changing climate at its home site, based on their norm of reaction (Figure 5). Green indicates increased fitness, purple indicates decreased fitness, and white indicates no change; minimum and maximum values for each metric are shown in the legend. Predictions are shown for changes in MCMT between the periods of 1961-1990 and 2041-2070 under SSP 2-45.

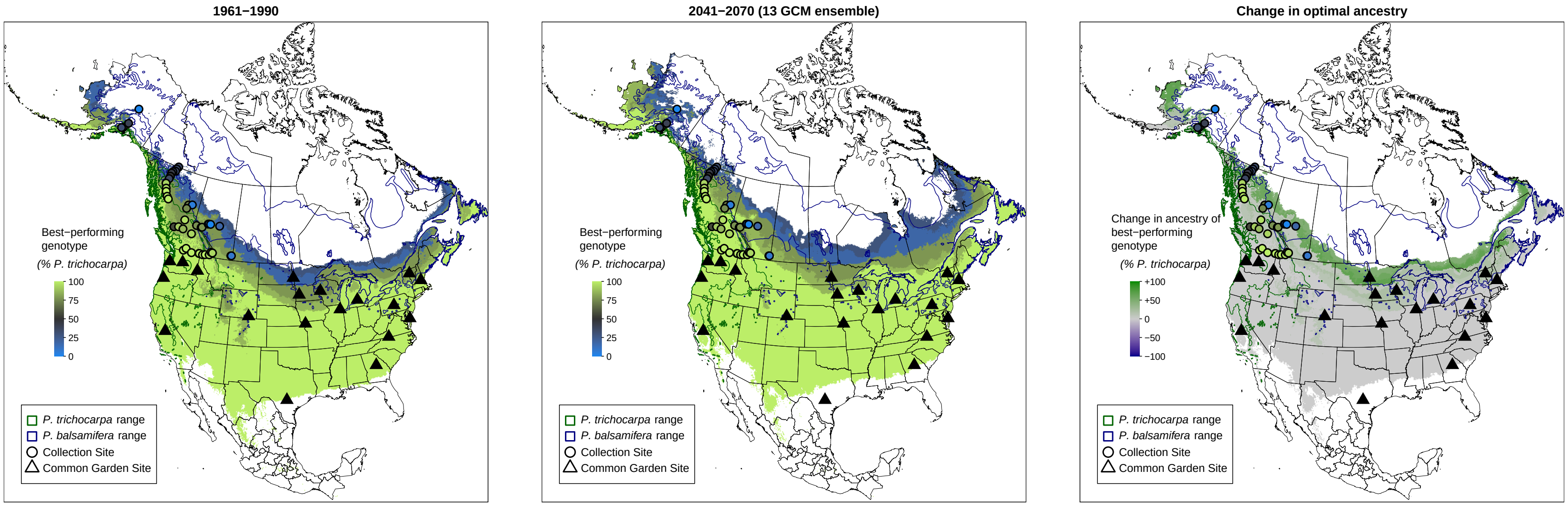


**Figure S15.** Maps showing the species ancestry of the studied genotype which is predicted to have highest fitness (as measured by growth and mortality) in that location under historic and future climate, indicated by the color of the base layer, across the ranges of both species and the common garden sites included in this study. "Change in optimal ancestry" indicates the change in optimal species ancestry between future and historic climates, indicating regions where increased *P. trichocarpa* ancestry is expected to be beneficial. See Figure 6 for corresponding figures inset to the sampled hybrid zone.

**Figure S16** (attached as a separate file). Predicted reaction norms across values of MCMT as in Figure 5a, with each genotype plotted separately (solid curve) with its MCMT of origin (vertical dotted line), allowing performance to be compared among the "local" genotype (MCMT of origin is the same as MCMT at planting site) and all other genotypes. Under the "local vs foreign" criterion of local adaptation, the genotype originating from a particular environment should outperform other genotypes in that environment (Kawecki and Ebert 2004).

**Supplementary Tables**

**Table S1.** List of common garden sites, abbreviations, and years measured. Sites that have a maxi garden or both a maxi and mini garden are indicated by parentheses ("maxi" and "both", respectively. All other sites are mini gardens.

| Common Garden Site | Abbreviation | City | State | Years Measured |
|---|---|---|---|---|
| Evergreen State | EVERGREEN | Olympia | WA | 2021-2023 |
| Lockerly Arboretum | LOCK | Milledgeville | GA | 2021 |
| Michigan State University | MSU | East Lansing | MI | 2021-2023 |
| Missouri Arboretum | NWMO | Maryville | MO | 2021-2022 |
| Morton Arboretum | MORTON | Lisle | IL | 2021-2023 |
| North Dakota State University (both) | NDSU | Fargo | ND | 2021-2022 |
| Oregon State University | OSU | Corvallis | OR | 2021-2023 |
| Our Lady of the Lake University | OLLU | San Antonio | TX | 2021 |
| Pennsylvania State University | PENN | State College | PA | 2021-2023 |
| Salisbury University Arboretum | SU | Salisbury | MD | 2021-2023 |
| Southwest Minnesota State University | SWMN | Marshall | MN | 2021-2023 |
| UC Merced | UCM | Merced | CA | 2021-2022 |
| University of Idaho | ID | Moscow | ID | 2021-2023 |
| University of Wisconsin - Eau Claire | WI | Eau Claire | WI | 2021-2023 |
| University of Wyoming | WYO | Laramie | WY | 2021-2023 |
| University of Vermont (maxi) | VT | Burlington | VT | 2021 |
| Virginia Tech (both) | VA | Critz | VA | 2021-2023 |
| Washington State - Wenatchee | WSU | Wenatchee | WA | 2021-2023 |

**Table S2.** Differences between AIC scores when different climate variables are used. MCMT has the lowest AIC score, and the dAIC column shows the increase in AIC score from the best model. Produced by the AICtab function from R package bbmle (version 1.0.25.1).

| | dAIC | df |
|---|---|---|
| MCMT | 0 | 41 |
| DD_0 | 3.3 | 41 |
| DD_18 | 6.1 | 41 |
| EMT | 12.3 | 41 |
| MAT | 23.6 | 41 |
| NFFD | 37.2 | 41 |
| Eref | 48.5 | 41 |
| TD | 54.9 | 41 |
| eFFP | 81.1 | 41 |
| DD5 | 88.6 | 41 |
| MAP | 88.7 | 41 |
| SHM | 91.9 | 41 |
| FFP | 93.1 | 41 |
| CMD | 97.6 | 41 |
| EXT | 99.4 | 41 |
| RH | 99.4 | 41 |
| bFFP | 102 | 41 |
| DD18 | 105.2 | 41 |
| AHM | 105.7 | 41 |
| MSP | 107 | 41 |
| PAS | 109.6 | 41 |
| MWMT | 114 | 41 |

**Table S3.** Effects of the linear mixed-effect model predicting yearly growth increment: home and garden MCMT, the square terms of MCMT, genetic PCs 1-3, and their interactions. Effects are shown for the conditional component testing each factor's effect on growth, and for the zero-inflated component testing the effect on the probability of mortality. Standardized beta coefficients are calculated by dividing the estimate for each predictor by its standard deviation to enable the relative effect sizes of predictors to be compared.

| **Conditional Model** | | | | | | |
|---|---|---|---|---|---|---|
| **Predictors** | **Estimates** | **std. Beta** | **CI** | **standardized CI** | **p** | **std. p** |
| (Intercept) | 2.85 | 2.73 | 2.13 – 3.57 | 2.48 – 2.98 | **<0.001** | **<0.001** |
| Garden MCMT | -0.28 | -0.53 | -0.83 – 0.28 | -0.74 – -0.31 | 0.33 | **<0.001** |
| Home MCMT | -0.42 | -0.27 | -1.57 – 0.72 | -0.65 – 0.11 | 0.468 | 0.169 |
| Garden $MCMT^2$ | -0.83 | -0.46 | -1.53 – -0.13 | -0.74 – -0.18 | **0.02** | **0.001** |
| Home $MCMT^2$ | -0.54 | -0.33 | -1.14 – 0.06 | -0.68 – 0.01 | 0.077 | 0.06 |
| Genetic PC1 | -0.33 | -0.3 | -0.46 – -0.21 | -0.41 – -0.18 | **<0.001** | **<0.001** |
| Genetic PC2 | 0.05 | -0.02 | -0.06 – 0.16 | -0.12 – 0.07 | 0.35 | 0.66 |
| Genetic PC3 | -0.18 | -0.15 | -0.29 – -0.08 | -0.24 – -0.05 | **0.001** | **0.003** |
| Garden MCMT × Home MCMT | 0.64 | 0.21 | -0.27 – 1.55 | -0.09 – 0.51 | 0.168 | 0.168 |
| Garden $MCMT^2$ × Home $MCMT^2$ | 0.23 | 0.11 | -0.33 – 0.79 | -0.16 – 0.37 | 0.427 | 0.427 |
| Home MCMT × Garden $MCMT^2$ | -0.1 | -0.03 | -1.19 – 0.98 | -0.32 – 0.27 | 0.855 | 0.855 |
| Garden MCMT × Home $MCMT^2$ | 0.38 | 0.21 | -0.10 – 0.85 | -0.06 – 0.49 | 0.121 | 0.121 |
| Garden MCMT × Genetic PC1 | -0.01 | -0.01 | -0.12 – 0.10 | -0.11 – 0.09 | 0.86 | 0.86 |
| Garden MCMT × Genetic PC2 | 0.01 | 0.01 | -0.07 – 0.09 | -0.06 – 0.08 | 0.796 | 0.796 |
| Garden MCMT × Genetic PC3 | -0.03 | -0.03 | -0.11 – 0.05 | -0.10 – 0.05 | 0.487 | 0.487 |
| Garden $MCMT^2$ × Genetic PC1 | 0.04 | 0.03 | -0.08 – 0.16 | -0.06 – 0.12 | 0.521 | 0.521 |
| Garden $MCMT^2$ × Genetic PC2 | -0.1 | -0.07 | -0.20 – 0.00 | -0.15 – 0.00 | 0.052 | 0.052 |
| Garden $MCMT^2$ × Genetic PC3 | 0.04 | 0.03 | -0.07 – 0.14 | -0.05 – 0.10 | 0.487 | 0.487 |
| (Intercept) | 0.69 | 0.69 | 0.65 – 0.73 | 0.65 – 0.73 | | |
| **Zero-Inflated Model** | | | | | | |
| **Predictors** | **Estimates** | **std. Beta** | **CI** | **standardized CI** | **p** | **std. p** |
| (Intercept) | -2.99 | -2.03 | -5.18 – -0.80 | -2.81 – -1.26 | **0.008** | **<0.001** |
| Garden MCMT | 0.05 | 0.74 | -1.31 – 1.42 | 0.10 – 1.37 | 0.94 | **0.023** |
| Home MCMT | -0.07 | 0.24 | -3.61 – 3.47 | -0.74 – 1.21 | 0.97 | 0.636 |
| Garden $MCMT^2$ | 2.08 | 1.34 | 0.11 – 4.05 | 0.72 – 1.95 | **0.039** | **<0.001** |
| Home $MCMT^2$ | 0.15 | 0.2 | -1.68 – 1.97 | -0.69 – 1.08 | 0.876 | 0.663 |

| | | | | | | |
|---|---|---|---|---|---|---|
| Genetic PC1 | 0.14 | 0.08 | -0.24 – 0.51 | -0.21 – 0.36 | 0.475 | 0.604 |
| Genetic PC2 | -0.21 | -0.17 | -0.54 – 0.12 | -0.42 – 0.07 | 0.217 | 0.163 |
| Genetic PC3 | -0.11 | 0.03 | -0.44 – 0.22 | -0.22 – 0.28 | 0.506 | 0.803 |
| Garden MCMT × Home MCMT | -1.54 | -0.51 | -3.69 – 0.62 | -1.22 – 0.21 | 0.164 | 0.164 |
| Garden $MCMT^2$ × Home $MCMT^2$ | -0.27 | -0.13 | -1.89 – 1.35 | -0.89 – 0.64 | 0.741 | 0.741 |
| Home MCMT × Garden $MCMT^2$ | 0.15 | 0.04 | -3.02 – 3.33 | -0.82 – 0.91 | 0.925 | 0.925 |
| Garden MCMT × Home $MCMT^2$ | -0.88 | -0.5 | -2.04 – 0.28 | -1.16 – 0.16 | 0.136 | 0.136 |
| Garden MCMT × Genetic PC1 | 0.04 | 0.03 | -0.20 – 0.28 | -0.18 – 0.25 | 0.756 | 0.756 |
| Garden MCMT × Genetic PC2 | 0.04 | 0.03 | -0.16 – 0.24 | -0.14 – 0.21 | 0.71 | 0.71 |
| Garden MCMT × Genetic PC3 | -0.02 | -0.02 | -0.22 – 0.18 | -0.20 – 0.16 | 0.843 | 0.843 |
| Garden $MCMT^2$ × Genetic PC1 | -0.06 | -0.05 | -0.39 – 0.27 | -0.30 – 0.20 | 0.711 | 0.711 |
| Garden $MCMT^2$ × Genetic PC2 | 0.07 | 0.05 | -0.24 – 0.37 | -0.17 – 0.27 | 0.671 | 0.671 |
| Garden $MCMT^2$ × Genetic PC3 | 0.2 | 0.15 | -0.10 – 0.50 | -0.08 – 0.38 | 0.192 | 0.192 |

**Table S4.** Effect sizes of the random effects of genotype, garden, block nested within garden, year, and individual and their sample sizes.

| **Random Effects** | |
|---|---|
| $\sigma2$ | 0.69 |
| $\tau00$ genotype | 0.05 |
| $\tau00$ block:Garden | 0 |
| $\tau00$ Garden | 0.24 |
| $\tau00$ year | 0 |
| $\tau00$ indiv | 0.06 |
| N genotype | 44 |
| N block | 35 |
| N Garden | 17 |
| N year | 2 |
| N indiv | 1448 |
| Observations | 2308 |
| Marginal R2 / Conditional R2 | 0.348 / NA |
| AIC | Inf |
