## Supplementary material for "Variation in responses to temperature in admixed *Populus* genotypes predicts geographic shifts in regions where hybrids are favored": Rmarkdown output files: climate_PCA_garden_provenance_historic_future.html


### Climate PCA

###### Alayna Mead

#### 2025-09-23

- 1 Setup
- 2 Reduce climate variables
- 3 PCA
- 4 Plots
  - 4.1 Setup
  - 4.2 Plot with base R
  - 4.3 Plot with ggplot
- 5 PCA - no lat/lon

### 1 Setup

```
library(vegan) # rda() function
```

```
## Loading required package: permute
```

```
## Loading required package: lattice
```

```
library(viridis) # color palette
```

```
## Loading required package: viridisLite
```

```
library(pals) # color palette
```

```
## 
## Attaching package: 'pals'
```

```
## The following objects are masked from 'package:viridis':
## 
##     cividis, inferno, magma, plasma, turbo, viridis
```

```
## The following objects are masked from 'package:viridisLite':
## 
##     cividis, inferno, magma, plasma, turbo, viridis
```

```
library(psych) # pairs.panels
```

```
## 
## Attaching package: 'psych'
```

```
## The following object is masked from 'package:vegan':
## 
##     pca
```

```
library(ggplot2)
```

```
## 
## Attaching package: 'ggplot2'
```

```
## The following objects are masked from 'package:psych':
## 
##     %+%, alpha
```

```
library(ggnewscale) # multiple color scales
library(ggtext) # for using markdown in legend title 'legend.title = element_markdown()'

# climates in long format - named 'all'
load(file = 'data/clean/546_genotypes_provenance_future_historic_climate_with_gardens_long_format.Rda')
clim <- all
rm(all)

# only need one set of future climates - remove 8 GCMs
clim <- clim[clim$period != "2041_2070_8GCMs",]

# remove SNHU, which doesn't get analyzed
clim <- clim[clim$site_name != 'SNHU',]

# ggplot theme
theme_set(theme_bw(base_size = 18))

sessionInfo()
```

```
## R version 4.5.1 (2025-06-13)
## Platform: x86_64-pc-linux-gnu
## Running under: Arch Linux
## 
## Matrix products: default
## BLAS:   /usr/lib/libblas.so.3.12.0 
## LAPACK: /usr/lib/liblapack.so.3.12.0  LAPACK version 3.12.0
## 
## locale:
##  [1] LC_CTYPE=en_US.UTF-8       LC_NUMERIC=C              
##  [3] LC_TIME=en_US.UTF-8        LC_COLLATE=en_US.UTF-8    
##  [5] LC_MONETARY=en_US.UTF-8    LC_MESSAGES=en_US.UTF-8   
##  [7] LC_PAPER=en_US.UTF-8       LC_NAME=C                 
##  [9] LC_ADDRESS=C               LC_TELEPHONE=C            
## [11] LC_MEASUREMENT=en_US.UTF-8 LC_IDENTIFICATION=C       
## 
## time zone: US/Eastern
## tzcode source: system (glibc)
## 
## attached base packages:
## [1] stats     graphics  grDevices datasets  utils     methods   base     
## 
## other attached packages:
##  [1] ggtext_0.1.2      ggnewscale_0.5.1  ggplot2_3.5.2     psych_2.5.3      
##  [5] pals_1.10         viridis_0.6.5     viridisLite_0.4.2 vegan_2.6-10     
##  [9] lattice_0.22-7    permute_0.9-7    
## 
## loaded via a namespace (and not attached):
##  [1] sass_0.4.10       generics_0.1.4    renv_0.17.3       xml2_1.3.8       
##  [5] digest_0.6.37     magrittr_2.0.3    evaluate_1.0.3    grid_4.5.1       
##  [9] fastmap_1.2.0     maps_3.4.2.1      jsonlite_2.0.0    Matrix_1.7-3     
## [13] gridExtra_2.3     mgcv_1.9-3        scales_1.3.0      jquerylib_0.1.4  
## [17] mnormt_2.1.1      cli_3.6.5         rlang_1.1.6       munsell_0.5.1    
## [21] splines_4.5.1     withr_3.0.2       cachem_1.1.0      yaml_2.3.10      
## [25] tools_4.5.1       parallel_4.5.1    dplyr_1.1.4       colorspace_2.1-1 
## [29] vctrs_0.6.5       mapproj_1.2.11    R6_2.6.1          lifecycle_1.0.4  
## [33] MASS_7.3-65       cluster_2.1.8.1   pkgconfig_2.0.3   pillar_1.10.2    
## [37] bslib_0.9.0       gtable_0.3.6      glue_1.8.0        Rcpp_1.0.14      
## [41] xfun_0.52         tibble_3.2.1      tidyselect_1.2.1  rstudioapi_0.17.1
## [45] knitr_1.50        dichromat_2.0-0.1 htmltools_0.5.8.1 nlme_3.1-168     
## [49] rmarkdown_2.29    compiler_4.5.1    gridtext_0.1.5
```

```
knitr::opts_chunk$set(fig.width = 10, fig.height = 8)
```

### 2 Reduce climate variables

```
# Derived annual variables:
# DD<0              degree-days below 0°C, chilling degree-days
# DD>5              degree-days above 5°C, growing degree-days
# DD<18            degree-days below 18°C, heating degree-days
# DD>18            degree-days above 18°C, cooling degree-days
# NFFD              the number of frost-free days
# FFP                 frost-free period
# bFFP               the day of the year on which FFP begins
# eFFP                the day of the year on which FFP ends
# PAS                 precipitation as snow (mm) between August in previous year and July in current year
# EMT                extreme minimum temperature over 30 years
# EXT                extreme maximum temperature over 30 years
# Eref                 Hargreaves reference evaporation (mm)
# CMD               Hargreaves climatic moisture deficit (mm)
# MAR               mean annual solar radiation (MJ m‐2 d‐1)
# RH                  mean annual relative humidity (%)
# CMI                Hogg’s climate moisture index (mm)
# DD1040 (10<DD<40)    degree-days above 10°C and below 40°C

vars <- c("lat", "lon", "elev", "AHM", "bFFP", "CMD", "CMI", "DD_0", "DD_18", "DD1040", "DD18", "DD5", "eFFP", "EMT", "Eref", "EXT", "FFP", "MAP", "MAR", "MAT", "MCMT", "MSP", "MWMT", "NFFD", "PAS", "RH", "SHM", "TD")

pairs.panels(clim[,vars], scale = T)
```

```
vars <- c("MWMT", "MCMT", "TD", "MAP", "MSP", "SHM", "PAS", "EMT", "CMD", "RH")

pairs.panels(clim[,vars], scale = T)
```

```
# remove EMT

vars <- c("MWMT", "MCMT", "TD", "MAP", "MSP", "SHM", "PAS", "CMD", "RH")

pairs.panels(clim[,vars], scale = T)
```

### 3 PCA

PCA of climate across all home sites and garden sites (both minis and
maxis), 544 genotypes, historic (1961-1990) and future (2041-2070, 13GCM
ensemble) climates, and yearly garden climates for 2020-2023.

Uses rda() funcion from vegan package

```
# reorder so gardens are plotted last
clim <- clim[order(clim$site_type, decreasing = T),]

# add site-period column
clim$site_period <- paste(clim$site_name, clim$period, sep = '_')


# climate variables to use

vars <- c("CMD", "MAP", "MAT", "MCMT", 'MWMT', "PAS", "RH", "TD", "lat", "lon")

# run PCA
rda <- rda(clim[,vars], scale = T)
summary(rda)
```

```
## 
## Call:
## rda(X = clim[, vars], scale = T) 
## 
## Partitioning of correlations:
##               Inertia Proportion
## Total              10          1
## Unconstrained      10          1
## 
## Eigenvalues, and their contribution to the correlations 
## 
## Importance of components:
##                          PC1    PC2    PC3     PC4     PC5     PC6      PC7
## Eigenvalue            4.5456 2.7689 1.0151 0.85760 0.48130 0.19717 0.086563
## Proportion Explained  0.4546 0.2769 0.1015 0.08576 0.04813 0.01972 0.008656
## Cumulative Proportion 0.4546 0.7314 0.8330 0.91872 0.96685 0.98656 0.995220
##                            PC8      PC9      PC10
## Eigenvalue            0.035955 0.011808 3.524e-05
## Proportion Explained  0.003595 0.001181 3.524e-06
## Cumulative Proportion 0.998816 0.999996 1.000e+00
```

```
biplot(rda)
```

```
plot(rda)
```

```
info <- summary(rda)
barplot(info$cont$importance[2,])
```

```
# look at loadings
rda$CA$v
```

```
##              PC1         PC2         PC3         PC4        PC5         PC6
## CMD  -0.33139551  0.18088495  0.44785296 -0.25043699 -0.4693566 -0.37784105
## MAP  -0.01351538 -0.52954680 -0.37386638 -0.02565451 -0.1978312  0.37395129
## MAT  -0.44706018 -0.12238613  0.03777786  0.17074528 -0.1080018  0.13919373
## MCMT -0.39860717 -0.28373756  0.19301183 -0.04391572  0.1052342  0.14721954
## MWMT -0.39347457  0.10994474 -0.10457306  0.40683537 -0.4300201  0.25159488
## PAS   0.23291213 -0.36841131 -0.19248702 -0.44044295 -0.5469254 -0.22259937
## RH    0.01619919 -0.47465443  0.03356597  0.58397176  0.0490824 -0.63275152
## TD    0.22442145  0.42294694 -0.30791895  0.32920463 -0.4194876 -0.01235377
## lat   0.42307778 -0.03218338  0.27616796  0.29463577 -0.1227027  0.10973278
## lon  -0.30422256  0.19581623 -0.63306764 -0.09912957  0.2008707 -0.38849614
##              PC7         PC8          PC9          PC10
## CMD  -0.45378789 -0.14463894  0.065294061 -8.703826e-05
## MAP  -0.62103105 -0.11186594  0.042340865 -4.761358e-04
## MAT   0.20728170 -0.31507416 -0.762429646  1.424927e-03
## MCMT  0.22065401 -0.02357206  0.359368578 -7.143429e-01
## MWMT  0.25030420  0.14977286  0.412733714  3.911379e-01
## PAS   0.47905796  0.01445529 -0.024503793  1.609758e-04
## RH   -0.06562230  0.15860554 -0.006718519  6.049217e-04
## TD   -0.10176971  0.12948359 -0.166111133 -5.802781e-01
## lat   0.09611579 -0.75822989  0.213640823 -6.466238e-05
## lon   0.02632042 -0.47672116  0.198164564  1.008468e-04
```

```
barplot(rda$CA$v[,1])
```

```
barplot(rda$CA$v[,2])
```

```
barplot(rda$CA$v[,3])
```

```
# PC1 mostly temperatures/latitude
# PC2 mostly precipitation/continentality
# PC3 mostly CMD/continentality
```

### 4 Plots

#### 4.1 Setup

```
# setup colors and shapes for nice plot

# color based on transect or garden
# provenances are colored by genotype ancestry, garden sites each have their own color

clim$gards <-  NA
clim$gards[clim$site_type == 'garden'] <- clim$site_name[clim$site_type == 'garden']

clim$gards <- factor(clim$gards, levels = c("EVERGREEN", "ID", "LOCK", "MORTON", "MSU", "NDSU","NWMO", "OLLU", "OSU", "PENN", "SU", "SWMN", "UCM", "VA", "VT", "WI", "WSU", "WYO"))


# colors for gardens - picking distinguishable colors from transects
kelly(22)
```

```
##  [1] "#F2F3F4" "#222222" "#F3C300" "#875692" "#F38400" "#A1CAF1" "#BE0032"
##  [8] "#C2B280" "#848482" "#008856" "#E68FAC" "#0067A5" "#F99379" "#604E97"
## [15] "#F6A600" "#B3446C" "#DCD300" "#882D17" "#8DB600" "#654522" "#E25822"
## [22] "#2B3D26"
```

```
# remove some colors from output
col_pal <- c("#008856",  "#8DB600", "#BE0032", "#875692", "#A1CAF1", "#2B3D26", "#C2B280", "#F3C300", "#848482", "#0067A5", "#E68FAC", "#F99379", "#604E97", "#DCD300", "#2b3e85",  "#B3446C", "#882D17", "#E25822")


names(col_pal) <- levels(clim$gards)

dput(col_pal)
```

```
## c(EVERGREEN = "#008856", ID = "#8DB600", LOCK = "#BE0032", MORTON = "#875692", 
## MSU = "#A1CAF1", NDSU = "#2B3D26", NWMO = "#C2B280", OLLU = "#F3C300", 
## OSU = "#848482", PENN = "#0067A5", SU = "#E68FAC", SWMN = "#F99379", 
## UCM = "#604E97", VA = "#DCD300", VT = "#2b3e85", WI = "#B3446C", 
## WSU = "#882D17", WYO = "#E25822")
```

```
# c(EVERGREEN = "#008856", ID = "#8DB600", LOCK = "#BE0032", MORTON = "#875692", 
# MSU = "#A1CAF1", NDSU = "#2B3D26", NWMO = "#C2B280", OLLU = "#F3C300", 
# OSU = "#848482", PENN = "#0067A5", SU = "#E68FAC", SWMN = "#F99379", 
# UCM = "#604E97", VA = "#DCD300", VT = "#2b3e85", WI = "#B3446C", 
# WSU = "#882D17", WYO = "#E25822")

# vector with colors
# start with genotype colors
clim$cols <- clim$color_Pt

# add garden colors

for(n in 1:nrow(clim)){
  
  if(clim$site_type[n] == 'garden'){
    clim$cols[n] <- col_pal[clim$gards[n]]
  }
  
}

# shape by year
shapes <- clim$period
shapes[shapes == '2020'] <- 22
shapes[shapes == '2021'] <- 23
shapes[shapes == '2022'] <- 24
shapes[shapes == '2023'] <- 25
shapes[shapes == '1961_1990'] <- 21
shapes[shapes == '2041_2070_13GCMs'] <- 1
shapes <- as.numeric(shapes)
```

#### 4.2 Plot with base R

```
# PC1 and PC2

# png(file = 'results/climate/climate_PCA_1-2_transects_and_gardens_4yrs.png', height = 8, width = 10, res = 300, units = 'in')

par(cex.lab = 1.5, mar = c(5,5,3,1))

choices = c(1,2)

plot(rda, choices = choices, type = 'none', xlim = c(-3.5, 4),
     xlab = paste('PC', choices[1], ' (', round(info$cont$importance[2,choices[1]]*100, 1), '% variance explained)', sep = ''),
     ylab = paste('PC', choices[2], ' (', round(info$cont$importance[2,choices[2]]*100, 1), '% variance explained)', sep = ''))
points(rda, choices = choices, display = 'sites', col = clim$cols,  pch = shapes, cex = ifelse(clim$site_type == 'garden', 1.5, 1), lwd = 2)
#text(rda, choices = choices, display = 'sites', col = col, cex = 0.5)
text(rda, choices = choices, display = 'species', col = 'black', cex = 1.5)

legend('topright', pch = c(1,2,3,0,16, 2), col = c(rep('black', 6)), pt.cex = 2, legend = c('2020', '2021', '2022', '2023','1961-1990', '2041-2070'))
```

```
#dev.off()

# PC1 and PC3
# png(file = 'results/climate/climate_PCA_1-3_transects_and_gardens_4yrs.png', height = 8, width = 10, res = 300, units = 'in')

par(cex.lab = 1.5, mar = c(5,5,3,1))

choices = c(1,3)

plot(rda, choices = choices, type = 'none', xlim = c(-3.5, 4),
     xlab = paste('PC', choices[1], ' (', round(info$cont$importance[2,choices[1]]*100, 1), '% variance explained)', sep = ''),
     ylab = paste('PC', choices[2], ' (', round(info$cont$importance[2,choices[2]]*100, 1), '% variance explained)', sep = ''))
points(rda, choices = choices, display = 'sites', col = clim$cols,  pch = shapes, cex = ifelse(clim$site_type == 'garden', 1, 1), lwd = 2)
#text(rda, choices = choices, display = 'sites', col = col, cex = 0.5)
text(rda, choices = choices, display = 'species', col = 'black', cex = 1.5)

legend('topright', pch = c(1,2,3,0,16, 2), col = c(rep('black', 6)), pt.cex = 2, legend = c('2020', '2021', '2022', '2023','1961-1990', '2041-2070'))
```

```
#dev.off()

# PC1 and PC3, version without legend
#png(file = 'results/climate/climate_PCA_1-3_transects_and_gardens_4yrs_nolegend.png', height = 8, width = 10, res = 300, units = 'in')

par(cex.lab = 1.5, mar = c(5,5,3,1))

choices = c(1,3)

plot(rda, choices = choices, type = 'none', xlim = c(-3.5, 4),
     xlab = paste('PC', choices[1], ' (', round(info$cont$importance[2,choices[1]]*100, 1), '% variance explained)', sep = ''),
     ylab = paste('PC', choices[2], ' (', round(info$cont$importance[2,choices[2]]*100, 1), '% variance explained)', sep = ''))
points(rda, choices = choices, display = 'sites', col = clim$cols,  pch = shapes, cex = ifelse(clim$site_type == 'garden', 1, 1), lwd = 2)
#text(rda, choices = choices, display = 'sites', col = col, cex = 0.5)
text(rda, choices = choices, display = 'species', col = 'black', cex = 1.5)
```

```
#dev.off()
```

#### 4.3 Plot with ggplot

```
#####################
# ggplot

# merge PCA data for each individual with information dataframe
to_plot <- cbind.data.frame(clim, rda$CA$u)

clims <- rda$CA$v

no_fut <- which(to_plot$period != '2041_2070_13GCMs')

# only minis
no_fut_minis <- which(to_plot$period != '2041_2070_13GCMs' & (to_plot$in_minis == TRUE | to_plot$site_type == 'garden'))

ggplot(dat = to_plot, aes(x = PC1, y = PC2)) +
  geom_line(dat = subset(to_plot, site_type == 'provenance' & in_minis == TRUE), aes(x = PC1, y = PC2, group = site_name), arrow = arrow(ends = 'first', angle = 20, length = unit(0.1, "inches")), col = rgb(0,0,0, alpha = 0.3)) + # arrows
  geom_point(dat = to_plot[no_fut_minis,], aes(x = PC1, y = PC2), bg = clim$cols[no_fut_minis], pch = shapes[no_fut_minis], cex = 2.5, show.legend = T) + # sites
  geom_text(data = clims, label = rownames(clims), aes(x = PC1/3, y = PC2/3), size = 8) + # climate loadings
  scale_color_manual(name = 'sites', breaks = names(col_pal), values = col_pal)
```

```
## Warning: No shared levels found between `names(values)` of the manual scale and the
## data's colour values.
```

```
ggplot(dat = to_plot, aes(x = PC1, y = PC2)) +
  geom_line(dat = subset(to_plot, site_type == 'provenance' & in_minis == TRUE), 
            aes(x = PC1, y = PC2, 
                group = site_name), 
            arrow = arrow(ends = 'first', angle = 20, length = unit(0.1, "inches")), col = rgb(0,0,0, alpha = 0.3)) + # arrows
  geom_point(dat = to_plot[no_fut_minis,], 
             aes(x = PC1, y = PC2), 
             bg = clim$cols[no_fut_minis], 
             pch = shapes[no_fut_minis], 
             cex = 2.5, 
             show.legend = T) + # sites
  geom_text(data = clims, 
            label = rownames(clims), 
            aes(x = PC1/3, y = PC2/3), 
            size = 8) + # climate loadings
  scale_color_manual(name = 'sites', breaks = names(col_pal), values = col_pal)
```

```
## Warning: No shared levels found between `names(values)` of the manual scale and the
## data's colour values.
```

```
# all years
ggplot(dat = to_plot, aes(x = PC1, y = PC2)) +
  geom_line(dat = subset(to_plot, site_type == 'provenance' & in_minis == TRUE), 
            aes(x = PC1, y = PC2, 
                group = site_name), 
            arrow = arrow(ends = 'first', angle = 20, length = unit(0.1, "inches")), col = rgb(0,0,0, alpha = 0.3)) + # arrows
  geom_point(dat = subset(to_plot[no_fut_minis,], site_type == 'garden'),
             aes(x = PC1, y = PC2, color = site_name, shape = period, size = period)) +
  scale_size_manual(values = c("1961_1990" = 3, "2041_2070_13GCMs" = 1, "2021" = 4, "2023" = 1, "2020" = 1, "2022" = 1)) +
  scale_color_manual(name = 'Garden', values = col_pal) + # plot gardens
  new_scale_color() +
  geom_point(dat = subset(to_plot[no_fut_minis,], site_type == 'provenance'), 
             aes(x = PC1, y = PC2, fill = Pt), pch = 21, size = 4) + 
  scale_fill_gradient2(name = '% P. trichocarpa ancestry',
                       low = 'dodgerblue2', 
                       mid = 'grey20', 
                       high = 'darkolivegreen2', 
                       midpoint = 0.5)
```

```
# only 2021 for minis
no_fut_minis <- which(to_plot$period != '2041_2070_13GCMs' & (to_plot$in_minis == TRUE | to_plot$site_type == 'garden'))
  
#no_fut_minis <- which(to_plot$period == '2021' & (to_plot$in_minis == TRUE | to_plot$site_type == 'garden'))
  
# PC1 and PC2

ggplot(dat = to_plot, aes(x = PC1, y = PC2)) +
  xlab(paste('PC1 (', round(info$cont$importance['Proportion Explained','PC1'],3)*100, '% variance explained)', sep = '')) +
  ylab(paste('PC2 (', round(info$cont$importance['Proportion Explained','PC2'],3)*100, '% variance explained)', sep = '')) +
  geom_vline(xintercept = 0, col = 'grey60') + 
  geom_hline(yintercept = 0, col = 'grey60') + 
  geom_line(dat = subset(to_plot, site_type == 'provenance' & in_minis == TRUE), 
            aes(x = PC1, y = PC2, 
                group = site_name), 
            arrow = arrow(ends = 'first', angle = 20, length = unit(0.1, "inches")), col = rgb(0,0,0, alpha = 0.3)) + # arrows
  geom_point(dat = subset(to_plot[no_fut_minis,], site_type == 'garden' & period == '2021'),
             aes(x = PC1, y = PC2, fill = site_name), pch = 24, size = 4) +
  scale_fill_manual(name = 'Garden', values = col_pal) + # plot gardens
  new_scale_fill() +
  geom_point(dat = subset(to_plot[no_fut_minis,], site_type == 'provenance'), 
             aes(x = PC1, y = PC2, fill = Pt), pch = 21, size = 4) + 
  scale_fill_gradient2(name = '% P. trichocarpa ancestry',
                       low = 'dodgerblue2', 
                       mid = 'grey20', 
                       high = 'darkolivegreen2', 
                       midpoint = 0.5) + # provenance points (historic)
  geom_text(data = clims, 
            label = rownames(clims), 
            aes(x = PC1/4, y = PC2/4), 
            size = 6) # climate loadings
```

```
#ggsave('results/climate/climatePCA_1-2_gardens2021_provenanceMinisPastFutureArrows.png', height = 8, width = 12)


# PC1 and PC3

ggplot(dat = to_plot, aes(x = PC1, y = PC3)) +
  xlab(paste('PC1 (', round(info$cont$importance['Proportion Explained','PC1'],3)*100, '% variance explained)', sep = '')) +
  ylab(paste('PC3 (', round(info$cont$importance['Proportion Explained','PC3'],3)*100, '% variance explained)', sep = '')) +
  geom_vline(xintercept = 0, col = 'grey60') + 
  geom_hline(yintercept = 0, col = 'grey60') + 
  geom_line(dat = subset(to_plot, site_type == 'provenance' & in_minis == TRUE), 
            aes(x = PC1, y = PC3, 
                group = site_name), 
            arrow = arrow(ends = 'first', angle = 20, length = unit(0.1, "inches")), col = rgb(0,0,0, alpha = 0.3)) + # arrows
  geom_point(dat = subset(to_plot[no_fut_minis,], site_type == 'garden' & period == '2021'),
             aes(x = PC1, y = PC3, fill = site_name), pch = 24, size = 4) +
  scale_fill_manual(name = 'Garden', values = col_pal) + # plot gardens
  new_scale_fill() +
  geom_point(dat = subset(to_plot[no_fut_minis,], site_type == 'provenance'), 
             aes(x = PC1, y = PC3, fill = Pt), pch = 21, size = 4) + 
  scale_fill_gradient2(name = '% P. trichocarpa ancestry',
                       low = 'dodgerblue2', 
                       mid = 'grey20', 
                       high = 'darkolivegreen2', 
                       midpoint = 0.5) + # provenance points (historic)
  geom_text(data = clims, 
            label = rownames(clims), 
            aes(x = PC1/4, y = PC3/4), 
            size = 6) # climate loadings
```

```
#ggsave('results/climate/climatePCA_1-3_gardens2021_provenanceMinisPastFutureArrows.png', height = 8, width = 12)


# PC2 and PC3

ggplot(dat = to_plot, aes(x = PC2, y = PC3)) +
  xlab(paste('PC2 (', round(info$cont$importance['Proportion Explained','PC2'],3)*100, '% variance explained)', sep = '')) +
  ylab(paste('PC3 (', round(info$cont$importance['Proportion Explained','PC3'],3)*100, '% variance explained)', sep = '')) +
  geom_vline(xintercept = 0, col = 'grey60') + 
  geom_hline(yintercept = 0, col = 'grey60') + 
  geom_line(dat = subset(to_plot, site_type == 'provenance' & in_minis == TRUE), 
            aes(x = PC2, y = PC3, 
                group = site_name), 
            arrow = arrow(ends = 'first', angle = 20, length = unit(0.1, "inches")), col = rgb(0,0,0, alpha = 0.3)) + # arrows
  geom_point(dat = subset(to_plot[no_fut_minis,], site_type == 'garden' & period == '2021'),
             aes(x = PC2, y = PC3, fill = site_name), pch = 24, size = 4) +
  scale_fill_manual(name = 'Garden', values = col_pal) + # plot gardens
  new_scale_fill() +
  geom_point(dat = subset(to_plot[no_fut_minis,], site_type == 'provenance'), 
             aes(x = PC2, y = PC3, fill = Pt), pch = 21, size = 4) + 
  scale_fill_gradient2(name = '% P. trichocarpa ancestry',
                       low = 'dodgerblue2', 
                       mid = 'grey20', 
                       high = 'darkolivegreen2', 
                       midpoint = 0.5) + # provenance points (historic)
  geom_text(data = clims, 
            label = rownames(clims), 
            aes(x = PC2/4, y = PC3/4), 
            size = 6) # climate loadings
```

```
#ggsave('results/climate/climatePCA_2-3_gardens2021_provenanceMinisPastFutureArrows.png', height = 8, width = 12)


##############################

# 2021 and 2022 for minis

ggplot(dat = to_plot, aes(x = PC1, y = PC2)) +
  xlab(paste('PC1 (', round(info$cont$importance['Proportion Explained','PC1'],3)*100, '% variance explained)', sep = '')) +
  ylab(paste('PC2 (', round(info$cont$importance['Proportion Explained','PC2'],3)*100, '% variance explained)', sep = '')) +
  geom_vline(xintercept = 0, col = 'grey60') + 
  geom_hline(yintercept = 0, col = 'grey60') + 
  geom_line(dat = subset(to_plot, site_type == 'provenance' & in_minis == TRUE), 
            aes(x = PC1, y = PC2, 
                group = site_name), 
            arrow = arrow(ends = 'first', angle = 20, length = unit(0.1, "inches")), col = rgb(0,0,0, alpha = 0.3)) + # arrows
  geom_point(dat = subset(to_plot[no_fut_minis,], site_type == 'garden' & period %in% c('2021', '2022'), ),
             aes(x = PC1, y = PC2, fill = site_name, shape = period),  size = 4) +
  scale_shape_manual(name = 'Year', values = c(24,22)) +
  scale_fill_manual(name = 'Garden', values = col_pal) + # plot gardens
    guides(shape = guide_legend(order = 1), 
         fill = guide_legend(order = 2, override.aes = list(shape = 22), ncol = 2)) + # order legends and get garden legend to show correct colors 
  new_scale_fill() +
  geom_point(dat = subset(to_plot[no_fut_minis,], site_type == 'provenance'), 
             aes(x = PC1, y = PC2, fill = Pt), pch = 21, size = 4) + 
  scale_fill_gradient2(name = "% *P. trichocarpa*<br>ancestry",
                       low = 'dodgerblue2', 
                       mid = 'grey20', 
                       high = 'darkolivegreen2', 
                       midpoint = 0.5) + # provenance points (historic)
  theme(legend.title = element_markdown()) +
  geom_text(data = clims, 
            label = rownames(clims), 
            aes(x = PC1/4, y = PC2/4), 
            size = 6) # climate loadings
```

```
#ggsave('results/climate/climatePCA_1-2_gardens2021-2022_provenanceMinisPastFutureArrows.png', height = 8, width = 12)


# PC1 and PC3

ggplot(dat = to_plot, aes(x = PC1, y = PC3)) +
  xlab(paste('PC1 (', round(info$cont$importance['Proportion Explained','PC1'],3)*100, '% variance explained)', sep = '')) +
  ylab(paste('PC3 (', round(info$cont$importance['Proportion Explained','PC3'],3)*100, '% variance explained)', sep = '')) +
  geom_vline(xintercept = 0, col = 'grey60') + 
  geom_hline(yintercept = 0, col = 'grey60') + 
  geom_line(dat = subset(to_plot, site_type == 'provenance' & in_minis == TRUE), 
            aes(x = PC1, y = PC3, 
                group = site_name), 
            arrow = arrow(ends = 'first', angle = 20, length = unit(0.1, "inches")), col = rgb(0,0,0, alpha = 0.3)) + # arrows
  geom_point(dat = subset(to_plot[no_fut_minis,], site_type == 'garden' & period %in% c('2021', '2022'), ),
             aes(x = PC1, y = PC3, fill = site_name, shape = period),  size = 4) +
  scale_shape_manual(name = 'Year', values = c(24,22)) +
  scale_fill_manual(name = 'Garden', values = col_pal) + # plot gardens
    guides(shape = guide_legend(order = 1), 
         fill = guide_legend(order = 2, override.aes = list(shape = 22), ncol = 2)) + # order legends and get garden legend to show correct colors 
  new_scale_fill() +
  geom_point(dat = subset(to_plot[no_fut_minis,], site_type == 'provenance'), 
             aes(x = PC1, y = PC3, fill = Pt), pch = 21, size = 4) + 
  scale_fill_gradient2(name = "% *P. trichocarpa*<br>ancestry",
                       low = 'dodgerblue2', 
                       mid = 'grey20', 
                       high = 'darkolivegreen2', 
                       midpoint = 0.5) + # provenance points (historic)
  theme(legend.title = element_markdown()) +
  geom_text(data = clims, 
            label = rownames(clims), 
            aes(x = PC1/4, y = PC3/4), 
            size = 6) # climate loadings
```

```
#ggsave('results/climate/climatePCA_1-3_gardens2021-2022_provenanceMinisPastFutureArrows.png', height = 8, width = 12)

################################
# all 544 genotypes, not just those in minis

# PC1 and PC2

ggplot(dat = to_plot, aes(x = PC1, y = PC2)) +
  xlab(paste('PC1 (', round(info$cont$importance['Proportion Explained','PC1'],3)*100, '% variance explained)', sep = '')) +
  ylab(paste('PC2 (', round(info$cont$importance['Proportion Explained','PC2'],3)*100, '% variance explained)', sep = '')) +
  geom_vline(xintercept = 0, col = 'grey60') + 
  geom_hline(yintercept = 0, col = 'grey60') + 
  geom_line(dat = subset(to_plot, site_type == 'provenance'), 
            aes(x = PC1, y = PC2, 
                group = site_name), 
            arrow = arrow(ends = 'first', angle = 20, length = unit(0.1, "inches")), col = rgb(0,0,0, alpha = 0.3)) + # arrows
  geom_point(dat = subset(to_plot, site_type == 'garden' & period == '2021'),
             aes(x = PC1, y = PC2, fill = site_name), pch = 24, size = 4) +
  scale_fill_manual(name = 'Garden', values = col_pal) + # plot gardens
  new_scale_fill() +
  geom_point(dat = subset(to_plot[no_fut,], site_type == 'provenance'), 
             aes(x = PC1, y = PC2, fill = Pt), pch = 21, size = 4) + 
  scale_fill_gradient2(name = '% P. trichocarpa ancestry',
                       low = 'dodgerblue2', 
                       mid = 'grey20', 
                       high = 'darkolivegreen2', 
                       midpoint = 0.5) + # provenance points (historic)
  geom_text(data = clims, 
            label = rownames(clims), 
            aes(x = PC1/4, y = PC2/4), 
            size = 6) # climate loadings
```

```
#ggsave('results/climate/climatePCA_1-2_gardens2021_provenanceAllPastFutureArrows.png', height = 8, width = 12)


# PC1 and PC3


ggplot(dat = to_plot, aes(x = PC1, y = PC3)) +
  xlab(paste('PC1 (', round(info$cont$importance['Proportion Explained','PC1'],3)*100, '% variance explained)', sep = '')) +
  ylab(paste('PC3 (', round(info$cont$importance['Proportion Explained','PC3'],3)*100, '% variance explained)', sep = '')) +
  geom_vline(xintercept = 0, col = 'grey60') + 
  geom_hline(yintercept = 0, col = 'grey60') + 
  geom_line(dat = subset(to_plot, site_type == 'provenance'), 
            aes(x = PC1, y = PC3, 
                group = site_name), 
            arrow = arrow(ends = 'first', angle = 20, length = unit(0.1, "inches")), col = rgb(0,0,0, alpha = 0.3)) + # arrows
  geom_point(dat = subset(to_plot, site_type == 'garden' & period == '2021'),
             aes(x = PC1, y = PC3, fill = site_name), pch = 24, size = 4) +
  scale_fill_manual(name = 'Garden', values = col_pal) + # plot gardens
  new_scale_fill() +
  geom_point(dat = subset(to_plot[no_fut,], site_type == 'provenance'), 
             aes(x = PC1, y = PC3, fill = Pt), pch = 21, size = 4) + 
  scale_fill_gradient2(name = '% P. trichocarpa ancestry',
                       low = 'dodgerblue2', 
                       mid = 'grey20', 
                       high = 'darkolivegreen2', 
                       midpoint = 0.5) + # provenance points (historic)
  geom_text(data = clims, 
            label = rownames(clims), 
            aes(x = PC1/4, y = PC3/4), 
            size = 6) # climate loadings
```

```
#ggsave('results/climate/climatePCA_1-3_gardens2021_provenanceAllPastFutureArrows.png', height = 8, width = 12)
```

### 5 PCA - no lat/lon

Remove latitude and lognitude, which are correlated with some of the
climatic variables.

```
# reorder so gardens are plotted last
clim <- clim[order(clim$site_type, decreasing = T),]

# add site-period column
clim$site_period <- paste(clim$site_name, clim$period, sep = '_')


# climate variables to use

vars <- c("CMD", "MAP", "MAT", "MCMT", 'MWMT', "PAS", "RH", "TD")

# run PCA
rda <- rda(clim[,vars], scale = T)
summary(rda)
```

```
## 
## Call:
## rda(X = clim[, vars], scale = T) 
## 
## Partitioning of correlations:
##               Inertia Proportion
## Total               8          1
## Unconstrained       8          1
## 
## Eigenvalues, and their contribution to the correlations 
## 
## Importance of components:
##                          PC1    PC2    PC3    PC4     PC5     PC6      PC7
## Eigenvalue            3.5457 2.6496 0.8360 0.5472 0.31946 0.08639 0.015647
## Proportion Explained  0.4432 0.3312 0.1045 0.0684 0.03993 0.01080 0.001956
## Cumulative Proportion 0.4432 0.7744 0.8789 0.9473 0.98724 0.99804 0.999996
##                             PC8
## Eigenvalue            3.525e-05
## Proportion Explained  4.406e-06
## Cumulative Proportion 1.000e+00
```

```
biplot(rda)
```

```
plot(rda)
```

```
info <- summary(rda)
barplot(info$cont$importance[2,])
```

```
# look at loadings
rda$CA$v
```

```
##              PC1           PC2        PC3        PC4         PC5         PC6
## CMD  -0.35841759  0.3016181428  0.4214767 -0.2155735 -0.59118618  0.45489245
## MAP  -0.08609709 -0.5521871145 -0.1352323 -0.3891942  0.31399992  0.64695490
## MAT  -0.52019799  0.0006535279 -0.1354497 -0.1206182  0.11145300 -0.18910180
## MCMT -0.50225933 -0.1636508840  0.1444310  0.1029873  0.13787135 -0.19673280
## MWMT -0.41782970  0.2135787135 -0.4423320 -0.4018011  0.01615456 -0.23491157
## PAS   0.21027076 -0.4467734571  0.3225293 -0.5729301 -0.29196425 -0.48757515
## RH   -0.09989816 -0.4614079308 -0.4878117  0.3595157 -0.63999431  0.01783541
## TD    0.33537837  0.3452740652 -0.4766756 -0.3974371 -0.15929802  0.08263090
##                PC7           PC8
## CMD   0.0045303488 -0.0001258185
## MAP   0.0277489184 -0.0004502574
## MAT  -0.8051796141  0.0016713696
## MCMT  0.3484399445 -0.7143980327
## MWMT  0.4629380812  0.3910814615
## PAS  -0.0305445317  0.0001521449
## RH   -0.0004049065  0.0005154744
## TD   -0.1193317569 -0.5802477406
```

```
barplot(rda$CA$v[,1])
```

```
barplot(rda$CA$v[,2])
```

```
barplot(rda$CA$v[,3])
```

```
# PC1 mostly temperatures
# PC2 mostly precipitation/continentality
# PC3 mostly CMD/MWMT/RH/continentality
```

```
# setup colors and shapes for nice plot

# color based on transect or garden
# provenances are colored by genotype ancestry, garden sites each have their own color

clim$gards <-  NA
clim$gards[clim$site_type == 'garden'] <- clim$site_name[clim$site_type == 'garden']

clim$gards <- factor(clim$gards, levels = c("EVERGREEN", "ID", "LOCK", "MORTON", "MSU", "NDSU","NWMO", "OLLU", "OSU", "PENN", "SU", "SWMN", "UCM", "VA", "VT", "WI", "WSU", "WYO"))


# colors for gardens - picking distinguishable colors from transects
kelly(22)
```

```
##  [1] "#F2F3F4" "#222222" "#F3C300" "#875692" "#F38400" "#A1CAF1" "#BE0032"
##  [8] "#C2B280" "#848482" "#008856" "#E68FAC" "#0067A5" "#F99379" "#604E97"
## [15] "#F6A600" "#B3446C" "#DCD300" "#882D17" "#8DB600" "#654522" "#E25822"
## [22] "#2B3D26"
```

```
# remove some colors from output
col_pal <- c("#008856",  "#8DB600", "#BE0032", "#875692", "#A1CAF1", "#2B3D26", "#C2B280", "#F3C300", "#848482", "#0067A5", "#E68FAC", "#F99379", "#604E97", "#DCD300", "#2b3e85",  "#B3446C", "#882D17", "#E25822")


names(col_pal) <- levels(clim$gards)

dput(col_pal)
```

```
## c(EVERGREEN = "#008856", ID = "#8DB600", LOCK = "#BE0032", MORTON = "#875692", 
## MSU = "#A1CAF1", NDSU = "#2B3D26", NWMO = "#C2B280", OLLU = "#F3C300", 
## OSU = "#848482", PENN = "#0067A5", SU = "#E68FAC", SWMN = "#F99379", 
## UCM = "#604E97", VA = "#DCD300", VT = "#2b3e85", WI = "#B3446C", 
## WSU = "#882D17", WYO = "#E25822")
```

```
# c(EVERGREEN = "#008856", ID = "#8DB600", LOCK = "#BE0032", MORTON = "#875692", 
# MSU = "#A1CAF1", NDSU = "#2B3D26", NWMO = "#C2B280", OLLU = "#F3C300", 
# OSU = "#848482", PENN = "#0067A5", SU = "#E68FAC", SWMN = "#F99379", 
# UCM = "#604E97", VA = "#DCD300", VT = "#2b3e85", WI = "#B3446C", 
# WSU = "#882D17", WYO = "#E25822")

# vector with colors
# start with genotype colors
clim$cols <- clim$color_Pt

# add garden colors

for(n in 1:nrow(clim)){
  
  if(clim$site_type[n] == 'garden'){
    clim$cols[n] <- col_pal[clim$gards[n]]
  }
}

# shape by year
shapes <- clim$period
shapes[shapes == '2020'] <- 22
shapes[shapes == '2021'] <- 23
shapes[shapes == '2022'] <- 24
shapes[shapes == '2023'] <- 25
shapes[shapes == '1961_1990'] <- 21
shapes[shapes == '2041_2070_13GCMs'] <- 1
shapes <- as.numeric(shapes)
```

```
#####################
# ggplot

# merge PCA data for each individual with information dataframe
to_plot <- cbind.data.frame(clim, rda$CA$u)

# change names to denote maxi gardens
to_plot[to_plot$site_name == 'VT', 'site_name'] <- 'VT (maxi)'
to_plot[to_plot$site_name == 'VA', 'site_name'] <- 'VA (both)'
to_plot[to_plot$site_name == 'NDSU', 'site_name'] <- 'NDSU (both)'


names(col_pal)[names(col_pal) == 'VT'] <- 'VT (maxi)'
names(col_pal)[names(col_pal) == 'VA'] <- 'VA (both)'
names(col_pal)[names(col_pal) == 'NDSU'] <- 'NDSU (both)'


clims <- rda$CA$v

no_fut <- which(to_plot$period != '2041_2070_13GCMs')

# only minis
no_fut_minis <- which(to_plot$period != '2041_2070_13GCMs' & (to_plot$in_minis == TRUE | to_plot$site_type == 'garden'))

ggplot(dat = to_plot, aes(x = PC1, y = PC2)) +
  geom_line(dat = subset(to_plot, site_type == 'provenance' & in_minis == TRUE), aes(x = PC1, y = PC2, group = site_name), arrow = arrow(ends = 'first', angle = 20, length = unit(0.1, "inches")), col = rgb(0,0,0, alpha = 0.3)) + # arrows
  geom_point(dat = to_plot[no_fut_minis,], aes(x = PC1, y = PC2), bg = clim$cols[no_fut_minis], pch = shapes[no_fut_minis], cex = 2.5, show.legend = T) + # sites
  geom_text(data = clims, label = rownames(clims), aes(x = PC1/3, y = PC2/3), size = 8) + # climate loadings
  scale_color_manual(name = 'sites', breaks = names(col_pal), values = col_pal)
```

```
## Warning: No shared levels found between `names(values)` of the manual scale and the
## data's colour values.
```

```
# 2021 and 2022 for minis

ggplot(dat = to_plot, aes(x = PC1, y = PC2)) +
  xlab(paste('PC1 (', round(info$cont$importance['Proportion Explained','PC1'],3)*100, '% variance explained)', sep = '')) +
  ylab(paste('PC2 (', round(info$cont$importance['Proportion Explained','PC2'],3)*100, '% variance explained)', sep = '')) +
  geom_vline(xintercept = 0, col = 'grey60') + 
  geom_hline(yintercept = 0, col = 'grey60') + 
  geom_line(dat = subset(to_plot, site_type == 'provenance' & in_minis == TRUE), 
            aes(x = PC1, y = PC2, 
                group = site_name), 
            arrow = arrow(ends = 'first', angle = 20, length = unit(0.1, "inches")), col = rgb(0,0,0, alpha = 0.3)) + # arrows
  geom_point(dat = subset(to_plot[no_fut_minis,], site_type == 'garden' & period %in% c('2021', '2022'), ),
             aes(x = PC1, y = PC2, fill = site_name, shape = period),  size = 4) +
  scale_shape_manual(name = 'Year', values = c(24,22)) +
  scale_fill_manual(name = 'Garden', values = col_pal) + # plot gardens
    guides(shape = guide_legend(order = 1), 
         fill = guide_legend(order = 2, override.aes = list(shape = 22), ncol = 2)) + # order legends and get garden legend to show correct colors 
  new_scale_fill() +
  geom_point(dat = subset(to_plot[no_fut_minis,], site_type == 'provenance'), 
             aes(x = PC1, y = PC2, fill = Pt), pch = 21, size = 4) + 
  scale_fill_gradient2(name = "% *P. trichocarpa*<br>ancestry",
                       low = 'dodgerblue2', 
                       mid = 'grey20', 
                       high = 'darkolivegreen2', 
                       midpoint = 0.5) + # provenance points (historic)
  theme(legend.title = element_markdown(),
        axis.text=element_text(size=20),
        axis.title=element_text(size=24)) +
  geom_text(data = clims, 
            label = rownames(clims), 
            aes(x = PC1/4, y = PC2/4), 
            size = 8) # climate loadings
```

```
#ggsave('results/climate/climatePCA_1-2_gardens2021-2022_provenanceMinisPastFutureArrows_noLatLon.png', height = 8, width = 12)
#ggsave('results/climate/climatePCA_1-2_gardens2021-2022_provenanceMinisPastFutureArrows_noLatLon.pdf', height = 8, width = 12)

# PC1 and PC3

ggplot(dat = to_plot, aes(x = PC1, y = PC3)) +
  xlab(paste('PC1 (', round(info$cont$importance['Proportion Explained','PC1'],3)*100, '% variance explained)', sep = '')) +
  ylab(paste('PC3 (', round(info$cont$importance['Proportion Explained','PC3'],3)*100, '% variance explained)', sep = '')) +
  geom_vline(xintercept = 0, col = 'grey60') + 
  geom_hline(yintercept = 0, col = 'grey60') + 
  geom_line(dat = subset(to_plot, site_type == 'provenance' & in_minis == TRUE), 
            aes(x = PC1, y = PC3, 
                group = site_name), 
            arrow = arrow(ends = 'first', angle = 20, length = unit(0.1, "inches")), col = rgb(0,0,0, alpha = 0.3)) + # arrows
  geom_point(dat = subset(to_plot[no_fut_minis,], site_type == 'garden' & period %in% c('2021', '2022'), ),
             aes(x = PC1, y = PC3, fill = site_name, shape = period),  size = 4) +
  scale_shape_manual(name = 'Year', values = c(24,22)) +
  scale_fill_manual(name = 'Garden', values = col_pal) + # plot gardens
    guides(shape = guide_legend(order = 1), 
         fill = guide_legend(order = 2, override.aes = list(shape = 22), ncol = 2)) + # order legends and get garden legend to show correct colors 
  new_scale_fill() +
  geom_point(dat = subset(to_plot[no_fut_minis,], site_type == 'provenance'), 
             aes(x = PC1, y = PC3, fill = Pt), pch = 21, size = 4) + 
  scale_fill_gradient2(name = "% *P. trichocarpa*<br>ancestry",
                       low = 'dodgerblue2', 
                       mid = 'grey20', 
                       high = 'darkolivegreen2', 
                       midpoint = 0.5) + # provenance points (historic)
  theme(legend.title = element_markdown()) +
  geom_text(data = clims, 
            label = rownames(clims), 
            aes(x = PC1/4, y = PC3/4), 
            size = 6) # climate loadings
```

```
#ggsave('results/climate/climatePCA_1-3_gardens2021-2022_provenanceMinisPastFutureArrows_noLatLon.png', height = 8, width = 12)
#ggsave('results/climate/climatePCA_1-3_gardens2021-2022_provenanceMinisPastFutureArrows_noLatLon.pdf', height = 8, width = 12)
```
