## Supplementary material for "Variation in responses to temperature in admixed *Populus* genotypes predicts geographic shifts in regions where hybrids are favored": Rmarkdown output files: map_best_genotype_by_climate.html

Map optimal genotypes by climate


### Map optimal genotypes by climate

###### Alayna Mead

#### 2025-09-24

- 1 Setup
  - 1.1 Load packages and data
  - 1.2 Setup genotype info
  - 1.3 Merge genotype responses
  - 1.4 Load climate rasters
- 2 Mapping (color by species
  ancestry)
  - 2.1 Functions
  - 2.2 Plot
- 3 Map change in ancestry
  - 3.1 Functions
  - 3.2 Plot

Using reaction norms modeled in the
‘transfer\_function\_multiyear\_linear\_mixed\_effects\_model.Rmd’ script,
identify the genotype having highest growth (taking into account
mortality probability) across the sampled hybrid zone under historic and
future values of MCMT. Maps show where the ‘optimal’ genotype/species
ancestry could shift spatially under future temperatures, resulting in
spatial shifts in the location of the hybrid zone.

### 1 Setup

#### 1.1 Load packages and data

```
library(sf) # mapping
```

```
## Linking to GEOS 3.13.1, GDAL 3.11.3, PROJ 9.6.0; sf_use_s2() is TRUE
```

```
library(terra) # mapping
```

```
## terra 1.8.54
```

```
library(RColorBrewer) # colors
library(circlize) # colorRamp2
```

```
## ========================================
## circlize version 0.4.16
## CRAN page: https://cran.r-project.org/package=circlize
## Github page: https://github.com/jokergoo/circlize
## Documentation: https://jokergoo.github.io/circlize_book/book/
## 
## If you use it in published research, please cite:
## Gu, Z. circlize implements and enhances circular visualization
##   in R. Bioinformatics 2014.
## 
## This message can be suppressed by:
##   suppressPackageStartupMessages(library(circlize))
## ========================================
```

```
# predictions of growth and mortality across climate for each genotype
load('results/model_prediction/predictedGrowth_acrossClimates_byGenotype_garden_MCMT_2021-2022.Rdata')
# rename
preds <- predictions
rm(predictions)

# make genotype info its own df
geno.info <- preds$genotypes
#cleanup
geno.info$genotype <- rownames(geno.info)
rownames(geno.info) <- paste('genotype_', rownames(geno.info), sep = '')


# garden data
load('data/clean/garden_climates_average1961-1990_yearly2020-2023.Rdata')
gards <- clim

# provenance data
# extract from garden climate and phenotype origin data - named 'dat'
load('data/clean/mini_garden_phenotypic_and_climate_data_2021-2023.Rdata')
# get first instance of genotype, extract genotype info
provs <- dat[! duplicated(dat$Genotype), c("Genotype", "Pb", "Pt", 
"transect", "provenance_latitude", "provenance_longitude", "provenance_elevation_m")]
# remove NA genotype
provs <- provs[! is.na(provs$Genotype),]
# remove genotypes without genetic data
provs <- provs[! is.na(provs$Pt),]

# change rownames to genotypes
rownames(provs) <- provs$Genotype

sessionInfo()
```

```
## R version 4.5.1 (2025-06-13)
## Platform: x86_64-pc-linux-gnu
## Running under: Arch Linux
## 
## Matrix products: default
## BLAS:   /usr/lib/libblas.so.3.12.0 
## LAPACK: /usr/lib/liblapack.so.3.12.0  LAPACK version 3.12.0
## 
## locale:
##  [1] LC_CTYPE=en_US.UTF-8       LC_NUMERIC=C              
##  [3] LC_TIME=en_US.UTF-8        LC_COLLATE=en_US.UTF-8    
##  [5] LC_MONETARY=en_US.UTF-8    LC_MESSAGES=en_US.UTF-8   
##  [7] LC_PAPER=en_US.UTF-8       LC_NAME=C                 
##  [9] LC_ADDRESS=C               LC_TELEPHONE=C            
## [11] LC_MEASUREMENT=en_US.UTF-8 LC_IDENTIFICATION=C       
## 
## time zone: US/Eastern
## tzcode source: system (glibc)
## 
## attached base packages:
## [1] stats     graphics  grDevices datasets  utils     methods   base     
## 
## other attached packages:
## [1] circlize_0.4.16    RColorBrewer_1.1-3 terra_1.8-54       sf_1.0-21         
## 
## loaded via a namespace (and not attached):
##  [1] cli_3.6.5           knitr_1.50          rlang_1.1.6        
##  [4] xfun_0.52           DBI_1.2.3           KernSmooth_2.23-26 
##  [7] renv_0.17.3         jsonlite_2.0.0      colorspace_2.1-1   
## [10] htmltools_0.5.8.1   e1071_1.7-16        GlobalOptions_0.1.2
## [13] sass_0.4.10         rmarkdown_2.29      grid_4.5.1         
## [16] evaluate_1.0.3      jquerylib_0.1.4     classInt_0.4-11    
## [19] fastmap_1.2.0       yaml_2.3.10         lifecycle_1.0.4    
## [22] compiler_4.5.1      codetools_0.2-20    Rcpp_1.0.14        
## [25] rstudioapi_0.17.1   digest_0.6.37       R6_2.6.1           
## [28] class_7.3-23        shape_1.4.6.1       magrittr_2.0.3     
## [31] bslib_0.9.0         tools_4.5.1         proxy_0.4-27       
## [34] units_0.8-7         cachem_1.1.0
```

```
knitr::opts_chunk$set(fig.width = 12, 
                      fig.height = 10)
```

#### 1.2 Setup genotype info

```
# get additional genotype info

# get first instances of each genotype
genos <- dat[match(geno.info$genotype, dat$Genotype),]

# check for match
cbind(as.character(genos$Genotype), geno.info$genotype)
```

```
##       [,1]  [,2] 
##  [1,] "206" "206"
##  [2,] "210" "210"
##  [3,] "218" "218"
##  [4,] "233" "233"
##  [5,] "255" "255"
##  [6,] "258" "258"
##  [7,] "307" "307"
##  [8,] "311" "311"
##  [9,] "317" "317"
## [10,] "333" "333"
## [11,] "334" "334"
## [12,] "342" "342"
## [13,] "353" "353"
## [14,] "364" "364"
## [15,] "374" "374"
## [16,] "380" "380"
## [17,] "381" "381"
## [18,] "405" "405"
## [19,] "411" "411"
## [20,] "416" "416"
## [21,] "419" "419"
## [22,] "423" "423"
## [23,] "427" "427"
## [24,] "432" "432"
## [25,] "437" "437"
## [26,] "443" "443"
## [27,] "453" "453"
## [28,] "463" "463"
## [29,] "469" "469"
## [30,] "522" "522"
## [31,] "533" "533"
## [32,] "543" "543"
## [33,] "545" "545"
## [34,] "564" "564"
## [35,] "567" "567"
## [36,] "572" "572"
## [37,] "588" "588"
## [38,] "601" "601"
## [39,] "808" "808"
## [40,] "821" "821"
## [41,] "827" "827"
## [42,] "865" "865"
## [43,] "972" "972"
## [44,] "973" "973"
```

```
# add to dataframe
geno.info$transect <- genos$transect
geno.info$interspecific_heterozygosity <- genos$interspecific_heterozygosity
geno.info$heterozygosity <- genos$heterozygosity
geno.info$hybrid_index <- genos$hybrid_index
geno.info$plastid_ID <- genos$plastid_ID
geno.info$Pt <- genos$Pt
geno.info$color_Pt <- genos$color_Pt
geno.info$color_k3 <- genos$color_k3

str(geno.info)
```

```
## 'data.frame':    44 obs. of  14 variables:
##  $ pc1                         : num  0.00423 -0.03207 -0.02393 -0.03496 -0.00995 ...
##  $ pc2                         : num  0.02407 -0.00722 0.01811 0.039 0.03374 ...
##  $ pc3                         : num  -0.0243 -0.0365 -0.0505 -0.0418 -0.0256 ...
##  $ home_clim                   : num  -9.6 -9.9 -8.9 -9.3 -12.2 -12.4 -6.9 -5.8 -5.8 -4.8 ...
##  $ Pt                          : num  0.571 0.908 0.864 0.966 0.715 ...
##  $ optimal_clim                : num  -4.04 -4.04 -3.66 -3.66 -4.79 ...
##  $ genotype                    : chr  "206" "210" "218" "233" ...
##  $ transect                    : Factor w/ 5 levels "Alaska","Cassiar",..: 3 3 3 3 3 3 5 5 5 5 ...
##  $ interspecific_heterozygosity: num  0.4066 NA 0.0956 NA 0.1766 ...
##  $ heterozygosity              : num  0.0996 0.0992 0.0925 0.0989 0.0942 ...
##  $ hybrid_index                : num  0.319 NA 0.0376 NA 0.1769 ...
##  $ plastid_ID                  : chr  "balsamifera" "trichocarpa" "balsamifera" "balsamifera" ...
##  $ color_Pt                    : chr  "#464A3BFF" "#A2C95FFF" "#95B75BFF" "#B2E065FF" ...
##  $ color_k3                    : chr  "#1B7F65" "#846912" "#4D961C" "#3DC200" ...
```

#### 1.3 Merge genotype responses

Merge each genotype’s response to MCMT into one dataframe for
comparison across genotypes

```
# get the overall predictions by genotype and join into one dataframe to identify optimal genotype for each climate

# get first genotype
pred.overall <- preds$overall[[1]]

# rename column
colnames(pred.overall)[2] <- names(preds$overall)[1]


# loop through other genotypes and add them
for(n in 2:length(preds$overall)){
  
  df <- preds$overall[[n]]
  
  pred.overall[,names(preds$overall)[n]] <- df[,2]
  
}

head(pred.overall)
```

```
##   garden_clim genotype_206 pred_se genotype_210  genotype_218  genotype_233
## 1   -23.90000 0.0001453033      NA  0.001661313 -0.0005488445 -0.0007477098
## 2   -23.52525 0.0008145990      NA  0.002649368  0.0002324151 -0.0005727152
## 3   -23.15051 0.0019332056      NA  0.004098252  0.0016045524 -0.0001682450
## 4   -22.77576 0.0037126400      NA  0.006188216  0.0038470405  0.0005955691
## 5   -22.40101 0.0064433609      NA  0.009159220  0.0073407967  0.0019025695
## 6   -22.02626 0.0105181509      NA  0.013327362  0.0125968296  0.0040081112
##   genotype_255 genotype_258  genotype_307  genotype_311 genotype_317
## 1  0.003805555   0.01860611 -0.0007562458 -0.0004039832 6.948694e-05
## 2  0.006222452   0.02524778 -0.0009025296 -0.0003998759 2.372378e-04
## 3  0.009765093   0.03396243 -0.0010361616 -0.0003212149 5.263518e-04
## 4  0.014860769   0.04530761 -0.0011261331 -0.0001126377 1.001396e-03
## 5  0.022070746   0.05996561 -0.0011221848  0.0003094876 1.755144e-03
## 6  0.032124017   0.07876507 -0.0009464785  0.0010684587 2.918889e-03
##    genotype_333  genotype_334  genotype_342 genotype_353 genotype_364
## 1 -1.095539e-04 -1.139752e-04 -7.107731e-05 0.0002346455  0.000644507
## 2 -8.759561e-05 -1.141680e-04 -6.451135e-05 0.0004022242  0.001024722
## 3 -2.756082e-05 -9.365559e-05 -3.840461e-05 0.0006624340  0.001599075
## 4  9.623987e-05 -3.648954e-05  2.170487e-05 0.0010584855  0.002454974
## 5  3.223790e-04  8.170086e-05  1.380191e-04 0.0016509541  0.003714517
## 6  7.075636e-04  2.972501e-04  3.438409e-04 0.0025237490  0.005546452
##   genotype_374 genotype_380 genotype_381 genotype_405 genotype_411 genotype_416
## 1  0.001303226  0.001900577  0.002171135    0.1026272   0.06154594   0.01065818
## 2  0.001971294  0.002792657  0.003239044    0.1254766   0.07962004   0.01462319
## 3  0.002941899  0.004062166  0.004773389    0.1529474   0.10256484   0.01993558
## 4  0.004336623  0.005852323  0.006955304    0.1858737   0.13157179   0.02700721
## 5  0.006320055  0.008354450  0.010027854    0.2252171   0.16809249   0.03636068
## 6  0.009112912  0.011821945  0.014314319    0.2720793   0.21388572   0.04865472
##   genotype_419 genotype_423 genotype_427 genotype_432 genotype_437
## 1   0.01979687  0.004140289  0.002022510  0.001742792 2.083826e-05
## 2   0.02695186  0.005848717  0.002905605  0.002599711 4.632462e-05
## 3   0.03646607  0.008203736  0.004140725  0.003835822 9.176714e-05
## 4   0.04903868  0.011426821  0.005854513  0.005601381 1.698369e-04
## 5   0.06555132  0.015806728  0.008213951  0.008099184 3.001007e-04
## 6   0.08710875  0.021717066  0.011437362  0.011600313 5.122747e-04
##    genotype_443  genotype_453  genotype_463  genotype_469  genotype_522
## 1 -3.405892e-05 -3.604828e-05 -0.0001412089 -0.0003234593 -6.376468e-05
## 2 -3.399126e-05 -4.152790e-05 -0.0001846170 -0.0004112153  1.887725e-04
## 3 -2.346953e-05 -4.152389e-05 -0.0002335555 -0.0005033438  6.356856e-04
## 4  7.975878e-06 -2.885478e-05 -0.0002828596 -0.0005846372  1.376603e-03
## 5  7.810963e-05  9.168883e-06 -0.0003216962 -0.0006264336  2.551654e-03
## 6  2.161200e-04  9.406120e-05 -0.0003299451 -0.0005788848  4.354868e-03
##   genotype_533 genotype_543 genotype_545 genotype_564 genotype_567 genotype_572
## 1 9.796994e-05   0.00997307  0.005722742   0.01015404   0.01669202   0.01806589
## 2 6.508648e-04   0.01433008  0.008893600   0.01465558   0.02289059   0.02396830
## 3 1.559214e-03   0.02030636  0.013404834   0.02084088   0.03108871   0.03156783
## 4 2.981932e-03   0.02841833  0.019722373   0.02924724   0.04184065   0.04128529
## 5 5.134050e-03   0.03932193  0.028446212   0.04055611   0.05582855   0.05362795
## 6 8.302429e-03   0.05384373  0.040341602   0.05562446   0.07388533   0.06920291
##   genotype_588 genotype_601  genotype_808  genotype_821 genotype_827
## 1   0.05864509  0.002224909 -4.528101e-05 -4.234797e-06 0.0005726378
## 2   0.07700620  0.003738267 -5.948961e-05  1.154623e-05 0.0009001163
## 3   0.10045541  0.005996458 -7.456155e-05  4.366055e-05 0.0013919919
## 4   0.13021542  0.009298395 -8.721068e-05  1.033954e-04 0.0021213891
## 5   0.16775328  0.014042733 -9.070855e-05  2.083540e-04 0.0031900826
## 6   0.21481549  0.020754936 -7.250923e-05  3.854411e-04 0.0047381810
##   genotype_865 genotype_972 genotype_973
## 1   0.08367808  0.001978613  0.006505981
## 2   0.10179996  0.002889569  0.009246759
## 3   0.12342780  0.004179856  0.012990662
## 4   0.14914923  0.005991342  0.018055764
## 5   0.17963117  0.008512826  0.024845478
## 6   0.21562559  0.011993422  0.033866880
```

```
# which genotype has highest predicted success for each value of MCMT (row)?

# just get genotypes
tmp <- pred.overall[,-1]


pred.overall$best_genotype <- sapply(1:nrow(tmp), function(x) names(which.max(tmp[x,])))

# get genotype ancestry and colors
pred.overall$best_genotype_Pt <- geno.info[pred.overall$best_genotype, 'Pt']
pred.overall$best_genotype_col_Pt <- geno.info[pred.overall$best_genotype, 'color_Pt']
pred.overall$best_genotype_col_k3 <- geno.info[pred.overall$best_genotype, 'color_k3']

pred.overall$best_genotype <- gsub('genotype_', '', pred.overall$best_genotype)

# quick plot to check
plot(pred.overall$garden_clim, col = pred.overall$best_genotype_col_Pt, pch = 16)
```

```
plot(pred.overall$garden_clim, col = pred.overall$best_genotype_col_k3, pch = 16)
```

```
# list 'best' genotype
cbind(pred.overall$garden_clim, pred.overall$best_genotype)
```

```
##        [,1]                 [,2] 
##   [1,] "-23.9"              "405"
##   [2,] "-23.5252525252525"  "405"
##   [3,] "-23.150505050505"   "405"
##   [4,] "-22.7757575757576"  "405"
##   [5,] "-22.4010101010101"  "405"
##   [6,] "-22.0262626262626"  "405"
##   [7,] "-21.6515151515151"  "405"
##   [8,] "-21.2767676767677"  "405"
##   [9,] "-20.9020202020202"  "405"
##  [10,] "-20.5272727272727"  "405"
##  [11,] "-20.1525252525253"  "588"
##  [12,] "-19.7777777777778"  "588"
##  [13,] "-19.4030303030303"  "588"
##  [14,] "-19.0282828282828"  "411"
##  [15,] "-18.6535353535354"  "411"
##  [16,] "-18.2787878787879"  "411"
##  [17,] "-17.9040404040404"  "411"
##  [18,] "-17.5292929292929"  "411"
##  [19,] "-17.1545454545455"  "411"
##  [20,] "-16.779797979798"   "411"
##  [21,] "-16.4050505050505"  "411"
##  [22,] "-16.030303030303"   "411"
##  [23,] "-15.6555555555556"  "411"
##  [24,] "-15.2808080808081"  "588"
##  [25,] "-14.9060606060606"  "588"
##  [26,] "-14.5313131313131"  "588"
##  [27,] "-14.1565656565657"  "588"
##  [28,] "-13.7818181818182"  "588"
##  [29,] "-13.4070707070707"  "588"
##  [30,] "-13.0323232323232"  "545"
##  [31,] "-12.6575757575758"  "545"
##  [32,] "-12.2828282828283"  "545"
##  [33,] "-11.9080808080808"  "545"
##  [34,] "-11.5333333333333"  "545"
##  [35,] "-11.1585858585859"  "545"
##  [36,] "-10.7838383838384"  "545"
##  [37,] "-10.4090909090909"  "545"
##  [38,] "-10.0343434343434"  "218"
##  [39,] "-9.65959595959596"  "218"
##  [40,] "-9.28484848484849"  "218"
##  [41,] "-8.91010101010101"  "218"
##  [42,] "-8.53535353535354"  "218"
##  [43,] "-8.16060606060606"  "218"
##  [44,] "-7.78585858585859"  "311"
##  [45,] "-7.41111111111111"  "311"
##  [46,] "-7.03636363636364"  "311"
##  [47,] "-6.66161616161616"  "311"
##  [48,] "-6.28686868686869"  "311"
##  [49,] "-5.91212121212121"  "311"
##  [50,] "-5.53737373737374"  "311"
##  [51,] "-5.16262626262627"  "307"
##  [52,] "-4.78787878787879"  "307"
##  [53,] "-4.41313131313132"  "307"
##  [54,] "-4.03838383838384"  "307"
##  [55,] "-3.66363636363637"  "307"
##  [56,] "-3.28888888888889"  "307"
##  [57,] "-2.91414141414142"  "307"
##  [58,] "-2.53939393939394"  "307"
##  [59,] "-2.16464646464647"  "307"
##  [60,] "-1.78989898989899"  "307"
##  [61,] "-1.41515151515152"  "307"
##  [62,] "-1.04040404040405"  "307"
##  [63,] "-0.665656565656569" "307"
##  [64,] "-0.290909090909096" "307"
##  [65,] "0.0838383838383798" "307"
##  [66,] "0.458585858585856"  "307"
##  [67,] "0.833333333333329"  "307"
##  [68,] "1.2080808080808"    "307"
##  [69,] "1.58282828282828"   "311"
##  [70,] "1.95757575757575"   "311"
##  [71,] "2.33232323232323"   "311"
##  [72,] "2.7070707070707"    "311"
##  [73,] "3.08181818181818"   "311"
##  [74,] "3.45656565656565"   "311"
##  [75,] "3.83131313131313"   "311"
##  [76,] "4.2060606060606"    "311"
##  [77,] "4.58080808080808"   "311"
##  [78,] "4.95555555555555"   "311"
##  [79,] "5.33030303030302"   "311"
##  [80,] "5.7050505050505"    "311"
##  [81,] "6.07979797979797"   "311"
##  [82,] "6.45454545454545"   "311"
##  [83,] "6.82929292929292"   "311"
##  [84,] "7.2040404040404"    "311"
##  [85,] "7.57878787878787"   "311"
##  [86,] "7.95353535353535"   "311"
##  [87,] "8.32828282828282"   "311"
##  [88,] "8.7030303030303"    "311"
##  [89,] "9.07777777777777"   "311"
##  [90,] "9.45252525252525"   "311"
##  [91,] "9.82727272727272"   "311"
##  [92,] "10.2020202020202"   "311"
##  [93,] "10.5767676767677"   "311"
##  [94,] "10.9515151515151"   "311"
##  [95,] "11.3262626262626"   "311"
##  [96,] "11.7010101010101"   "405"
##  [97,] "12.0757575757576"   "405"
##  [98,] "12.450505050505"    "405"
##  [99,] "12.8252525252525"   "405"
## [100,] "13.2"               "405"
```

```
# save
# save(pred.overall, file = 'results/model_prediction/genotypePredictedHeights_byGardenMCMT.Rdata')
# write.csv(pred.overall, file = 'results/model_prediction/genotypePredictedHeights_byGardenMCMT.csv', row.names = F)
```

#### 1.4 Load climate rasters

Load future and historic MCMT raster files

ClimateNA raster files used here are available from DataBasin:

https://adaptwest.databasin.org/pages/adaptwest-climatena/

Species range shapefiles are from Little 1971 and are available from
DataBasin:

P. balsamifera: https://databasin.org/datasets/91380e091ca048359a66fc65962ed210/

P. trichocarpa: https://databasin.org/datasets/84e47784fe2a463c8b292007fb43f2cd/

```
# which climate variable is being used? 
clim_var <- 'MCMT'
# calculate 2-year average, used in model
dat$garden_MCMT_2020_2021_avg <- (dat$garden_MCMT_2020 + dat$garden_MCMT_2021)/2
gard_clim_colname <- 'garden_MCMT_2020_2021_avg'

# get historic and future climate

hist <- rast(paste('data/climate/climateNA/Normal_1961_1990/Normal_1961_1990_bioclim/Normal_1961_1990_', clim_var, '.tif', sep = ''))
crs(hist, proj = T)
```

```
## [1] "+proj=laea +lat_0=45 +lon_0=-100 +x_0=0 +y_0=0 +datum=WGS84 +units=m +no_defs"
```

```
hist91 <- rast(paste('data/climate/climateNA/Normal_1991_2020/Normal_1991_2020_', clim_var, '.tif', sep = ''))
crs(hist, proj = T)
```

```
## [1] "+proj=laea +lat_0=45 +lon_0=-100 +x_0=0 +y_0=0 +datum=WGS84 +units=m +no_defs"
```

```
fut8 <- rast(paste('data/climate/climateNA/future/ensemble_8GCMs_ssp245_2041_2070_bioclim/ensemble_8GCMs_ssp245_2041_2070_', clim_var, '.tif', sep = ''))

fut13 <- rast(paste('data/climate/climateNA/future/ensemble_13GCMs_ssp245_2041_2070_bioclim/ensemble_13GCMs_ssp245_2041_2070_', clim_var, '.tif', sep = ''))

################ convert coords

# convert coords to crs of climateNA
crs.cna <- crs(hist, proj = T)

# provenance coordinates
coords.prov <- st_as_sf(provs[,c("provenance_longitude", "provenance_latitude")], coords = c(1,2), crs = st_crs(4326))
coords.prov.cna <- st_transform(coords.prov$geometry, crs = crs.cna)

# garden coordinates
coords.gards <- st_as_sf(gards[,c("Longitude", "Latitude")], coords = c(1,2), crs = st_crs(4326))
coords.gards.cna <- st_transform(coords.gards$geometry, crs = crs.cna)

# load shapefile of state/province borders
borders <- read_sf('data/shapefiles/NorthAmerica_PoliticalBoundaries_Shapefile/NA_PoliticalDivisions/data/bound_p/boundaries_p_2021_v3.shp')
borders <- st_transform(borders, crs = crs.cna)
# simplify to 1 km
borders <- st_simplify(borders, dTolerance = 1000)

# species rangemaps
# shapefile
balsam <- st_read('data/shapefiles/Pbal_shapefile/data/commondata/data0/popubals.shp')
```

```
## Reading layer `popubals' from data source 
##   `/home/alayna/Documents/research/projects/2023_populus_common_gardens/data/shapefiles/Pbal_shapefile/data/commondata/data0/popubals.shp' 
##   using driver `ESRI Shapefile'
## Simple feature collection with 415 features and 5 fields
## Geometry type: POLYGON
## Dimension:     XY
## Bounding box:  xmin: -18247640 ymin: 4662920 xmax: -5857014 ymax: 10826350
## Projected CRS: WGS 84 / Pseudo-Mercator
```

```
tricho <- st_read('data/shapefiles/Ptri_shapefile/data/commondata/data0/poputric.shp')
```

```
## Reading layer `poputric' from data source 
##   `/home/alayna/Documents/research/projects/2023_populus_common_gardens/data/shapefiles/Ptri_shapefile/data/commondata/data0/poputric.shp' 
##   using driver `ESRI Shapefile'
## Simple feature collection with 450 features and 5 fields
## Geometry type: POLYGON
## Dimension:     XY
## Bounding box:  xmin: -17154180 ymin: 3609620 xmax: -11488690 ymax: 8896231
## Projected CRS: WGS 84 / Pseudo-Mercator
```

```
# convert CRS
balsam <- st_transform(balsam, crs.cna)
tricho <- st_transform(tricho, crs.cna)

# test plot
plot(hist)

plot(balsam$geometry, add = T, border = 'navy')
plot(tricho$geometry, add = T, border = 'grey50')
plot(borders$geometry, add = T)
```

### 2 Mapping (color by species ancestry)

#### 2.1 Functions

```
# function for plotting
map_best_geno <- function(raster, raster_name, save = F, ...){
  
  if(save == T){
    pdf(file = paste('results/model_prediction/best_genotype_2yearModel_mapped_by_', clim_var, '_', raster_name, '.pdf', sep = ''),
   height = 8, width = 8)
  }
  
  
  plot(raster,
       col = colf.pt(seq(0,1,0.01)),
       breaks = seq(0,1,0.01),
       pax = list(side=NA),
       colNA = 'white',
       legend = F,
       ...)
  
  plot(borders$geometry, add = T, lwd = 0.5)
  plot(balsam$geometry, add = T, border = 'navy')
  plot(tricho$geometry, add = T, border = 'darkgreen')
  plot(coords.prov.cna, add = T, lwd = 2, bg = geno.info$color_Pt, pch = 21, cex = 1.5)
  plot(coords.gards.cna, add = T, pch = 17, cex = 2)

  
    legend(-3e6, -1.8e6, 
         pch = c(22,22,21,24),
         pt.cex = 2,
         xjust = 0.5,
         col =  c('darkgreen', 'navy', 'black','black'), 
         pt.lwd = 2,
         pt.bg = c('white', 'white', 'white', 'white'),
         legend = c(substitute(paste(italic('P. trichocarpa'), ' range')), 
                    substitute(paste(italic('P. balsamifera'), ' range')),
                    'Collection Site',
                    'Common Garden Site'))
  
  
  
  # add inset legend for species ancestry gradient
  # https://stackoverflow.com/questions/13355176/gradient-legend-in-base

  # get breaks for legend labels from color palette
  pal_range <- range(attr(colf.pt, 'breaks'))
  pal_min <- pal_range[1]
  pal_max <- pal_range[2]

  legend_image <- as.raster(matrix(colf.pt(seq(pal_max, pal_min, length = 100)), ncol=1))

  # look at NDC coords
  grconvertX(seq(-4e6, 3e6, 1000000), from = 'user', to = 'ndc')
  grconvertY(seq(-3e6, 4e6, 1000000), from = 'user', to = 'ndc')

  figSet <- c(0.1, 0.4, 0.15, 0.5)
  op <- par(  ## set and store par
    fig=figSet,    ## set figure region,
    mar=c(1, 1, 1, 9.5),                                  ## set margins
    new=TRUE)                                ## set new for overplot w/ next plot

  plot(0,0, type='n', axes=F, xlab='', ylab='')  ## ini plot2
  rasterImage(legend_image, 0, 0, 1, 1)   ## the gradient
  lbsq <- seq.int(0, 1, l=5) ## seq. for labels
  axis(4, at=lbsq, pos=1, labels=F, col=0, col.ticks=1, tck=-.1)  ## axis ticks
  mtext(c(0,25,50,75,100), 4, 0.3, at=lbsq, las=2, cex=.8)  ## tick labels

  mtext(expression(atop('Best-performing\ngenotype', italic('(% P. trichocarpa)'))), side=3, line=0.2, cex=1, adj=.1) ## legend title

  par(op)  ## reset par
  
  
  if(save == T){
    dev.off()
  }
}

######################################
# inset zoomed to collection sites


# function
map_best_geno_crop <- function(raster, raster_name, save = F, legends = T, ...){
  
  if(save == T){
    pdf(file = paste('results/model_prediction/best_genotype_2yearModel_mapped_by_', clim_var, '_inset_', raster_name, '.pdf', sep = ''),
        height = 8, width = 6)
  }
  
  
  plot(raster,
       col = colf.pt(seq(0,1,0.01)),
       breaks = seq(0,1,0.01),
       pax = list(side=NA),
       colNA = 'white',
       legend = F,
       ...)
  
  plot(borders$geometry, add = T, lwd = 0.5)
  plot(balsam$geometry, add = T, border = 'navy')
  plot(tricho$geometry, add = T, border = 'darkgreen')
  plot(coords.prov.cna, add = T, lwd = 2, bg = geno.info$color_Pt, pch = 21, cex = 2)
  
  
  if(legends == T){
    
    legend(-2.53e6, 0.9e6,
           pch = c(22,22,21),
           pt.cex = 2,
           xjust = 0.5,
           pt.bg =  c('darkgreen', 'navy', 'white'),
           legend = c(substitute(paste(italic('P. trichocarpa'), ' range')),
                      substitute(paste(italic('P. balsamifera'), ' range')),
                      'Collection Site'))
    
    
    # add inset legend for species ancestry gradient
    # https://stackoverflow.com/questions/13355176/gradient-legend-in-base
    
    # get breaks for legend labels from color palette
    pal_range <- range(attr(colf.pt, 'breaks'))
    pal_min <- pal_range[1]
    pal_max <- pal_range[2]
    
    legend_image <- as.raster(matrix(colf.pt(seq(pal_max, pal_min, length = 100)), ncol=1))
    
    # look at NDC coords
    grconvertX(seq(-4e6, 3e6, 1000000), from = 'user', to = 'ndc')
    grconvertY(seq(-3e6, 4e6, 1000000), from = 'user', to = 'ndc')
    
    figSet <- c(0.06, 0.5, 0.15, 0.5)
    op <- par(  # set and store par
      fig=figSet, # set figure region,
      mar=c(1, 1, 1, 9.5), # set margins
      new=TRUE) # set new for overplot w/ next plot
    
    plot(0,0, type='n', axes=F, xlab='', ylab='')  ## ini plot2
    rasterImage(legend_image, 0, 0, 1, 1)  ## the gradient
    lbsq <- seq.int(0, 1, l=5) ## seq. for labels
    axis(4, at=lbsq, pos=1, labels=F, col=0, col.ticks=1, tck=-.1)  ## axis ticks
    mtext(c(0, 25, 50, 75, 100), 4, 0.3, at=lbsq, las=2, cex=.8)  ## tick labels
    
    mtext(expression(atop('Best-performing\ngenotype', italic('(% P. trichocarpa)'))), side=3, line=0.2, cex=1, adj=.1)          ## legend title
    
    par(op)  ## reset par
    
    
  }
  
  if(save == T){
    dev.off()
  }
}


# testing
# extent <- ext(-3e6, -5e5, 45e4, 3e6)
# hist.crop <- crop(hist.mask.anc, extent)
# map_best_geno_crop(hist.crop, main = '1961-1990', save = save, raster_name = 'historic')
# map_best_geno_crop(hist.crop, main = '1961-1990', save = save, raster_name = 'historic_nolegend', legends = F)
```

#### 2.2 Plot

```
# color scale for ancestry
colf.pt <- colorRamp2(c(0,0.5,1), colors = c('dodgerblue2', 'grey20', 'darkolivegreen2'))


# make values outside prediction range (common gardens) NA
# get the range of actual garden climates that were tested
pred_clim <- c(dat$garden_MCMT_2021, dat$garden_MCMT_2022)

msk <- ifel(hist > max(pred_clim) | hist < min(pred_clim), NA, 1)
hist.mask <- mask(hist, msk)
plot(hist.mask, colNA = 'grey')
```

```
plot(hist.mask, colNA = 'grey')

hist91.mask <- mask(hist91, msk)
plot(hist91.mask, colNA = 'grey')
```

```
# future climate

# make values outside predict range NA
msk <- ifel(fut13 > max(pred_clim) | fut13 < min(pred_clim), NA, 1)
fut13.mask <- mask(fut13, msk)
```

```
## |---------|---------|---------|---------|=========================================
```

```
plot(fut13.mask, colNA = 'grey')
```

```
msk <- ifel(fut8 > max(pred_clim) | fut8 < min(pred_clim), NA, 1)
fut8.mask <- mask(fut8, msk)
plot(fut8.mask, colNA = 'grey')
```

```
# use classify to convert from MCMT values to the ancestry of the 'optimal' genotype

# make a matrix to pass to classify()
# columns 1-2 are the range of MCMT values, column 3 is the ancestry value to convert to
mat <- cbind.data.frame(from = pred.overall$garden_clim, to = NA, geno = pred.overall$best_genotype_Pt)
# add interval to get range of temps
mat$to <- mat$from + (mat$from[2] - mat$from[1])

# reclassify on all 4 rasters
hist.mask.anc <- classify(hist.mask, mat, include.lowest = T, right = T, others = NA)
hist91.mask.anc <- classify(hist91.mask, mat, include.lowest = T, right = T, others = NA)
fut8.mask.anc <- classify(fut8.mask, mat, include.lowest = T, right = T, others = NA)
fut13.mask.anc <- classify(fut13.mask, mat, include.lowest = T, right = T, others = NA)
```

```
## |---------|---------|---------|---------|=========================================
```

```
# quick plots to check
plot(hist.mask.anc)
```

```
plot(hist91.mask.anc)
```

```
plot(fut8.mask.anc)
```

```
plot(fut13.mask.anc)
```

```
# Maps!

# are we saving plots?
save = FALSE

# plot full range
map_best_geno(hist.mask.anc, raster_name = 'historic', main = '1961-1990', save = save)
```

```
map_best_geno(hist91.mask.anc, raster_name = '1991-2020', main = '1991-2020', save = save)
```

```
map_best_geno(fut13.mask.anc, raster_name = '2041-2070_13GCMs', main = '2041-2070 (13 GCM ensemble)', save = save)
```

```
map_best_geno(fut8.mask.anc, raster_name = '2041-2070_8GCMs', main = '2041-2070 (8 GCM ensemble)', save = save)
```

```
# crop rasters, then plot
extent <- ext(-3e6, -5e5, 45e4, 3e6)
hist.crop <- crop(hist.mask.anc, extent)
hist91.crop <- crop(hist91.mask.anc, extent)
fut8.crop <- crop(fut8.mask.anc, extent)
fut13.crop <- crop(fut13.mask.anc, extent)


map_best_geno_crop(hist.crop, main = '1961-1990', save = save, raster_name = 'historic')
```

```
map_best_geno_crop(hist91.crop, raster_name = '1991-2020', main = '1991-2020', save = save)
```

```
map_best_geno_crop(fut8.crop, main = '2041-2070 (8 GCM ensemble)', save = save, raster_name = '2041-2070_8GCMs')
```

```
map_best_geno_crop(fut13.crop, main = '2041-2070 (13 GCM ensemble)', save = save, raster_name = '2041-2070_13GCMs')
```

```
# version without legends
map_best_geno_crop(hist.crop, main = '1961-1990', save = save, raster_name = 'historic_nolegend', legends = F)
```

```
map_best_geno_crop(hist91.crop, raster_name = '1991-2020_nolegend', main = '1991-2020', legends = F, save = save)
```

```
map_best_geno_crop(fut8.crop, main = '2041-2070 (8 GCM ensemble)', save = save, raster_name = '2041-2070_8GCMs_nolegend', legends = F)
```

```
map_best_geno_crop(fut13.crop, main = '2041-2070 (13 GCM ensemble)', save = save, raster_name = '2041-2070_13GCMs_nolegend', legends = F)
```

```
# crop to western interior region

extent <- ext(-15e5, 1e5, -10e5, 1e6)
hist.crop <- crop(hist.mask.anc, extent)
hist91.crop <- crop(hist91.mask.anc, extent)
fut8.crop <- crop(fut8.mask.anc, extent)
fut13.crop <- crop(fut13.mask.anc, extent)

map_best_geno_crop(hist.crop, main = '1961-1990', save = save, raster_name = 'west_historic', legends  = F)
```

```
map_best_geno_crop(hist91.crop, raster_name = 'west_1991-2020', main = '1991-2020', save = save, legends  = F)
```

```
map_best_geno_crop(fut8.crop, main = '2041-2070 (8 GCM ensemble)', save = save, raster_name = 'west_2041-2070_8GCMs', legends  = F)
```

```
map_best_geno_crop(fut13.crop, main = '2041-2070 (13 GCM ensemble)', save = save, raster_name = 'west_2041-2070_13GCMs', legends  = F)
```

### 3 Map change in ancestry

Now calculate the difference in optimal ancestry between future and
historic maps to more clearly show where we expect changes (mostly
increases in tricho ancestry).

#### 3.1 Functions

```
# function for plotting
map_diff_geno <- function(raster, raster_name, save = F, ...){
  
  if(save == T){
    pdf(file = paste('results/model_prediction/best_genotype_difference_2yearModel_mapped_by_', clim_var, '_', raster_name, '.pdf', sep = ''),
   height = 8, width = 8)
  }
  
  
  plot(raster,
       col = colf.diff(seq(-1,1,0.01)), 
       breaks = seq(-1,1,0.01),
       pax = list(side=NA),
       colNA = 'white',
       legend = F,
       ...)
  
  plot(borders$geometry, add = T, lwd = 0.5)
  plot(balsam$geometry, add = T, border = 'navy')
  plot(tricho$geometry, add = T, border = 'darkgreen')
  plot(coords.prov.cna, add = T, lwd = 2, bg = geno.info$color_Pt, pch = 21, cex = 1.5)
  plot(coords.gards.cna, add = T, pch = 17, cex = 2)

  
    legend(-3e6, -1.8e6, 
         pch = c(22,22,21,24),
         pt.cex = 2,
         xjust = 0.5,
         col =  c('darkgreen', 'navy', 'black','black'), 
         pt.lwd = 2,
         pt.bg = c('white', 'white', 'white', 'white'),
         legend = c(substitute(paste(italic('P. trichocarpa'), ' range')), 
                    substitute(paste(italic('P. balsamifera'), ' range')),
                    'Collection Site',
                    'Common Garden Site'))
  
  
  
  # add inset legend for species ancestry gradient
  # https://stackoverflow.com/questions/13355176/gradient-legend-in-base

  # get breaks for legend labels from color palette
  pal_range <- range(attr(colf.diff, 'breaks'))
  pal_min <- pal_range[1]
  pal_max <- pal_range[2]

  legend_image <- as.raster(matrix(colf.diff(seq(pal_max, pal_min, length = 100)), ncol=1))

  # look at NDC coords
  grconvertX(seq(-4e6, 3e6, 1000000), from = 'user', to = 'ndc')
  grconvertY(seq(-3e6, 4e6, 1000000), from = 'user', to = 'ndc')

  figSet <- c(0.1, 0.4, 0.15, 0.5)
  op <- par(  ## set and store par
    fig=figSet,    ## set figure region,
    mar=c(1, 1, 1, 9.5),  ## set margins
    new=TRUE)  ## set new for overplot w/ next plot

  plot(0,0, type='n', axes=F, xlab='', ylab='')  ## ini plot2
  rasterImage(legend_image, 0, 0, 1, 1) ## the gradient
  lbsq <- seq.int(0, 1, l=5) ## seq. for labels
  axis(4, at=lbsq, pos=1, labels=F, col=0, col.ticks=1, tck=-.1)  ## axis ticks
  mtext(c('-100', '-50', '0', '+50', '+100'), 4, 0.3, at=lbsq, las=2, cex=.8) ## tick labels

  mtext(expression(atop('Change in ancestry of\nbest-performing\ngenotype', italic('(% P. trichocarpa)'), )), side=3, line=0.2, cex=1, adj=0.1)  ## legend title

  par(op)  ## reset par
  
  
  if(save == T){
    dev.off()
  }
}

######################################
# inset zoomed to collection sites


# function
map_diff_geno_crop <- function(raster, raster_name, save = F, legends = T, ...){
  
  if(save == T){
    # png(file = paste('results/model_prediction/best_genotype_difference_2yearModel_mapped_by_', clim_var, '_inset_', raster_name, '.png', sep = ''),
    #     height = 8, width = 6, res = 300, units = 'in')
    pdf(file = paste('results/model_prediction/best_genotype_difference_2yearModel_mapped_by_', clim_var, '_inset_', raster_name, '.pdf', sep = ''),
        height = 8, width = 6)
  }
  
  
  plot(raster,
       col = colf.diff(seq(-1,1,0.01)), 
       breaks = seq(-1,1,0.01),
       pax = list(side=NA),
       colNA = 'white',
       legend = F,
       ...)
  
  plot(borders$geometry, add = T, lwd = 0.5)
  plot(balsam$geometry, add = T, border = 'navy')
  plot(tricho$geometry, add = T, border = 'darkgreen')
  plot(coords.prov.cna, add = T, lwd = 2, bg = geno.info$color_Pt, pch = 21, cex = 2)
  
  
  if(legends == T){
    
    legend(-2.53e6, 0.9e6,
           pch = c(22,22,21,24),
           pt.cex = 2,
           xjust = 0.5,
           pt.bg =  c('darkgreen', 'navy', 'white'),
           legend = c(substitute(paste(italic('P. trichocarpa'), ' range')),
                      substitute(paste(italic('P. balsamifera'), ' range')),
                      'Collection Site'))
    
    
    # add inset legend for species ancestry gradient
    # https://stackoverflow.com/questions/13355176/gradient-legend-in-base
    
    # get breaks for legend labels from color palette
    pal_range <- range(attr(colf.diff, 'breaks'))
    pal_min <- pal_range[1]
    pal_max <- pal_range[2]
    
    legend_image <- as.raster(matrix(colf.diff(seq(pal_max, pal_min, length = 100)), ncol=1))
    
    # look at NDC coords
    grconvertX(seq(-4e6, 3e6, 1000000), from = 'user', to = 'ndc')
    grconvertY(seq(-3e6, 4e6, 1000000), from = 'user', to = 'ndc')
    
    figSet <- c(0.06, 0.5, 0.15, 0.5)
    op <- par(  # set and store par
      fig=figSet, # set figure region,
      mar=c(1, 1, 1, 9.5), # set margins
      new=TRUE) # set new for overplot w/ next plot
    
    plot(0,0, type='n', axes=F, xlab='', ylab='')  ## ini plot2
    rasterImage(legend_image, 0, 0, 1, 1)  ## the gradient
    lbsq <- seq.int(0, 1, l=5) ## seq. for labels
    axis(4, at=lbsq, pos=1, labels=F, col=0, col.ticks=1, tck=-.1)  ## axis ticks
    mtext(c('-100', '-50', '0', '+50', '+100'), 4, 0.3, at=lbsq, las=2, cex=.8)  ## tick labels
    
    mtext(expression(atop('Change in ancestry of\nbest-performing\ngenotype', italic('(% P. trichocarpa)'), )), side=3, line=0.2, cex=1, adj=0.1)  ## legend title
    
    par(op)  ## reset par
  }
  if(save == T){
    dev.off()
  }
}


# map_diff_geno_crop(hist.diff.crop)
# 
# map_diff_geno_crop(hist.diff.crop, main = 'Change in optimal ancestry', save = save, raster_name = 'ancestry_difference', legends  = T)
```

#### 3.2 Plot

```
# map difference in species ancestry of optimal genotype

# new raster with difference
hist.diff <- fut13.mask.anc - hist.mask.anc

# color palette: green = more tricho, blue = more balsam
colf.diff <- colorRamp2(c(-1,0,1), colors = c('blue4', 'grey80', 'green4'))


# maps
save = FALSE

# full map
map_diff_geno(hist.diff, main = 'Change in optimal ancestry', save = save, raster_name = 'ancestry_difference')
```

```
# crop to hybrid zone region

extent <- ext(-3e6, -5e5, 45e4, 3e6)
hist.diff.crop <- crop(hist.diff, extent)

map_diff_geno_crop(hist.diff.crop, main = 'Change in optimal ancestry', save = save, raster_name = 'ancestry_difference_nolegends', legends  = F)
map_diff_geno_crop(hist.diff.crop, main = 'Change in optimal ancestry', save = save, raster_name = 'ancestry_difference', legends  = T)
```

```
# look at regions with increasing balsam ancestry
extent2 <- ext(-2e6, -12e5, 15e5, 22e5)
hist.diff.crop <- crop(hist.diff, extent2)
map_diff_geno_crop(hist.diff.crop, legend = F, main = 'Change in optimal ancestry',  save = save, raster_name = 'ancestry_difference_zoom_to_balsam_regions')
```
