## Supplementary material for "Variation in responses to temperature in admixed *Populus* genotypes predicts geographic shifts in regions where hybrids are favored": Rmarkdown output files: predict_growth_future_climates.html

Predict growth changes under future climate


### Predict growth changes under future climate

###### Alayna Mead

#### 2025-05-12

- 1 Setup
  - 1.1 Load libraries and data
  - 1.2 Back-transform input data
  - 1.3 Extract genotype info
- 2 Get future climates for each
  genotype
  - 2.1 Load climate data
  - 2.2 Extract future climates for each
    genotype’s home site
- 3 Predict height at future
  climates
  - 3.1 Predict functions
  - 3.2 Predict response to historic and
    future climate for each genotype
  - 3.3 Calculate change in fitness under
    future climates
- 4 Map predictions for each collected
  genotype/site
- 5 Map rangewide predictions for each
  genotype
  - 5.1 Map past and future fitness
    predictions and change for each genotype

Using the genotype-specific modeled reaction norms produced by
‘transfer\_function\_multiyear\_linear\_mixed\_effects\_model.Rmd’ script,
predict how each genotype’s growth and survival could change under
future temperatures (represented by MCMT).

### 1 Setup

#### 1.1 Load libraries and data

```
library(ggplot2)
library(glmmTMB) # model
library(sf) # mapping
```

```
## Linking to GEOS 3.13.1, GDAL 3.10.2, PROJ 9.6.0; sf_use_s2() is TRUE
```

```
library(terra) # climate rasters
```

```
## terra 1.8.42
```

```
library(RColorBrewer) # colors
library(circlize) # colorRamp2
```

```
## ========================================
## circlize version 0.4.16
## CRAN page: https://cran.r-project.org/package=circlize
## Github page: https://github.com/jokergoo/circlize
## Documentation: https://jokergoo.github.io/circlize_book/book/
## 
## If you use it in published research, please cite:
## Gu, Z. circlize implements and enhances circular visualization
##   in R. Bioinformatics 2014.
## 
## This message can be suppressed by:
##   suppressPackageStartupMessages(library(circlize))
## ========================================
```

```
# load garden data - named 'dat'
# remove the 2024 data
load('data/clean/mini_garden_phenotypic_and_climate_data_2021-2024.Rdata')
dat <- dat[, !endsWith(colnames(dat), '2024')]

# load model info
# mod: model
# df.scaled: scaled mini garden data that went into the model
# scaling_factor: used to scale variables before fitting model
load('results/model_prediction/glmTMB_multiyear_model_outputs_garden_MCMT_2021-2022_vs_GrowthIncrement_2021-2022.Rda')

# load predicted heights for each genotype - named 'predictions'
load('results/model_prediction/predictedGrowth_acrossClimates_byGenotype_garden_MCMT_2021-2022.Rdata')
preds <- predictions
rm(predictions)

sessionInfo()
```

```
## R version 4.5.0 (2025-04-11)
## Platform: x86_64-pc-linux-gnu
## Running under: Arch Linux
## 
## Matrix products: default
## BLAS:   /usr/lib/libblas.so.3.12.0 
## LAPACK: /usr/lib/liblapack.so.3.12.0  LAPACK version 3.12.0
## 
## locale:
##  [1] LC_CTYPE=en_US.UTF-8       LC_NUMERIC=C              
##  [3] LC_TIME=en_US.UTF-8        LC_COLLATE=en_US.UTF-8    
##  [5] LC_MONETARY=en_US.UTF-8    LC_MESSAGES=en_US.UTF-8   
##  [7] LC_PAPER=en_US.UTF-8       LC_NAME=C                 
##  [9] LC_ADDRESS=C               LC_TELEPHONE=C            
## [11] LC_MEASUREMENT=en_US.UTF-8 LC_IDENTIFICATION=C       
## 
## time zone: US/Eastern
## tzcode source: system (glibc)
## 
## attached base packages:
## [1] stats     graphics  grDevices datasets  utils     methods   base     
## 
## other attached packages:
## [1] circlize_0.4.16    RColorBrewer_1.1-3 terra_1.8-42       sf_1.0-20         
## [5] glmmTMB_1.1.11     ggplot2_3.5.2     
## 
## loaded via a namespace (and not attached):
##  [1] sass_0.4.10         generics_0.1.3      renv_0.17.3        
##  [4] class_7.3-23        shape_1.4.6.1       KernSmooth_2.23-26 
##  [7] lattice_0.22-6      lme4_1.1-37         digest_0.6.37      
## [10] magrittr_2.0.3      evaluate_1.0.3      grid_4.5.0         
## [13] fastmap_1.2.0       jsonlite_2.0.0      Matrix_1.7-3       
## [16] e1071_1.7-16        DBI_1.2.3           GlobalOptions_0.1.2
## [19] mgcv_1.9-1          scales_1.3.0        codetools_0.2-20   
## [22] numDeriv_2016.8-1.1 jquerylib_0.1.4     reformulas_0.4.0   
## [25] Rdpack_2.6.4        cli_3.6.4           rlang_1.1.6        
## [28] units_0.8-7         rbibutils_2.3       munsell_0.5.1      
## [31] splines_4.5.0       withr_3.0.2         cachem_1.1.0       
## [34] yaml_2.3.10         tools_4.5.0         nloptr_2.2.1       
## [37] minqa_1.2.8         dplyr_1.1.4         colorspace_2.1-1   
## [40] boot_1.3-31         vctrs_0.6.5         R6_2.6.1           
## [43] proxy_0.4-27        lifecycle_1.0.4     classInt_0.4-11    
## [46] MASS_7.3-65         pkgconfig_2.0.3     pillar_1.10.2      
## [49] bslib_0.9.0         gtable_0.3.6        glue_1.8.0         
## [52] Rcpp_1.0.14         xfun_0.52           tibble_3.2.1       
## [55] tidyselect_1.2.1    knitr_1.50          htmltools_0.5.8.1  
## [58] nlme_3.1-168        rmarkdown_2.29      TMB_1.9.17         
## [61] compiler_4.5.0
```

```
knitr::opts_chunk$set(fig.width = 8, fig.height = 10)
```

#### 1.2 Back-transform input data

Data used for input to model was scaled - back-transform it to the
actual values.

```
df <- df.scaled

for(n in 1:length(scaling_factor)){
  
  column <- names(scaling_factor)[n]
  
  df[,column] <- df.scaled[,column]/scaling_factor[[n]]
  
}

head(df)
```

```
##                      pheno genotype garden  block           indiv year
## NDSU.405.1.1.24_2022     0      405   NDSU NDSU.1 NDSU.405.1.1.24 2022
## NDSU.411.2.4.3_2022      0      411   NDSU NDSU.2  NDSU.411.2.4.3 2022
## NDSU.419.1.1.20_2022     0      419   NDSU NDSU.1 NDSU.419.1.1.20 2022
## NDSU.419.2.3.8_2022      0      419   NDSU NDSU.2  NDSU.419.2.3.8 2022
## NDSU.416.1.2.3_2022      0      416   NDSU NDSU.1  NDSU.416.1.2.3 2022
## NDSU.590.1.2.11_2022     0      590   NDSU NDSU.1 NDSU.590.1.2.11 2022
##                      garden_clim garden_clim_2 home_clim home_clim_2       Pt
## NDSU.405.1.1.24_2022       -16.5        272.25     -23.9      571.21 0.334682
## NDSU.411.2.4.3_2022        -16.5        272.25     -20.0      400.00 0.293048
## NDSU.419.1.1.20_2022       -16.5        272.25     -16.7      278.89 0.324367
## NDSU.419.2.3.8_2022        -16.5        272.25     -16.7      278.89 0.324367
## NDSU.416.1.2.3_2022        -16.5        272.25     -16.4      268.96 0.315421
## NDSU.590.1.2.11_2022       -16.5        272.25     -14.6      213.16       NA
##                             pc1        pc2        pc3         pc4        pc5
## NDSU.405.1.1.24_2022 0.02748668 0.02411587 0.03816905 0.000560667 0.03818182
## NDSU.411.2.4.3_2022  0.03132797 0.02236233 0.03757786 0.012868191 0.04275311
## NDSU.419.1.1.20_2022 0.02841758 0.02582109 0.02883672 0.008416795 0.02996169
## NDSU.419.2.3.8_2022  0.02841758 0.02582109 0.02883672 0.008416795 0.02996169
## NDSU.416.1.2.3_2022  0.03009858 0.02374161 0.03372735 0.012894037 0.04524013
## NDSU.590.1.2.11_2022         NA         NA         NA          NA         NA
```

#### 1.3 Extract genotype info

Get genetic and climate data for each genotype

```
# get info for each genotype to use for extracting climate data and predicting height under future climates

# columns to get - these will be the same for each replicate of a genotype
dput(colnames(dat))
```

```
## c("Unique_ID", "MiniCG_Site", "Genotype", "Plant_ID", "Pb", "Pt", 
## "transect", "provenance_latitude", "provenance_longitude", "provenance_elevation_m", 
## "block", "row", "column", "Survival_09_2021", "PreFlush_Height_cm_2021", 
## "PostSet_Height_cm_2021", "GrowthIncrement_2021", "PetioleColor_2021", 
## "RustDisease_2021", "DOY_Stage2_2021", "DOY_Stage3_2021", "DOY_Stage6_2021", 
## "Stage2_cGDD_2021", "Stage3_cGDD_2021", "MAT_TransferDist", "MAP_TransferDist", 
## "MiniCG_Site_2022", "Unique_ID_2022", "Genotype_2022", "PLANT_ID_2022", 
## "Survival_09_2022", "PreFlush_Height_cm_2022", "PostSet_Height_cm_2022", 
## "GrowthIncrement_2022", "RustDisease_2022", "DOY_Stage2_2022", 
## "DOY_Stage3_2022", "DOY_Stage6_2022", "Stage2_cGDD_2022", "Stage3_cGDD_2022", 
## "Unique_ID_2023", "MiniCG_Site_2023", "Genotype_2023", "block_2023", 
## "row_2023", "column_2023", "DOY_Stage2.DD.MM.YY_2023", "DOY_Stage3.DD.MM.YY_2023", 
## "DOY_Stage6.DD.MM.YY_2023", "DOY_Stage7.DD.MM.YY_2023", "DOY_Stage8.DD.MM.YY_2023", 
## "Survival_09_2023", "PreFlush_Height_cm_2023", "PostSet_Height_cm_2023", 
## "GrowthIncrement_2023", "RustDisease.1.or.0_2023", "DOY_Stage2_2023", 
## "DOY_Stage3_2023", "DOY_Stage6_2023", "DOY_Stage7_2023", "DOY_Stage8_2023", 
## "Notes_2023", "DOY_leaf_measurements.DD.MM.YY_2023", "DOY_leaf_measurements_2023", 
## "leaf_measurement_growth_stage_2023", "leaf_thickness_1_mm_2023", 
## "leaf_thickness_2_mm_2023", "leaf_thickness_3_mm_2023", "leaf_thickness_4_mm_2023", 
## "leaf_thickness_5_mm_2023", "leaf_thickness_avg_mm_2023", "leaf_thickness_sd_mm_2023", 
## "DOY_LICOR_measurement.DD.MM.YY_2023", "DOY_LICOR_measurement_2023", 
## "weather_LICOR_measurement_2023", "LICOR_codes_2023", "LICOR_notes_2023", 
## "Survival_09_2021_2023", "notes_2023", "leaf_mass_g_2023", "leaf_area_cm2_2023", 
## "LMA_g_m2_2023", "DOY_Stage2.DD.MM.YY_2021", "DOY_Stage3.DD.MM.YY_2021", 
## "DOY_Stage6.DD.MM.YY_2021", "DOY_Stage2.DD.MM.YY_2022", "DOY_Stage3.DD.MM.YY_2022", 
## "DOY_Stage6.DD.MM.YY_2022", "total_growth_increment_2021_2022", 
## "total_growth_increment_2021_2023", "rgr_2021", "rgr_2022", "rgr_2023", 
## "DOY_last_budset_2023", "stage7_presence_2023", "growing_season_days_2021", 
## "growing_season_days_2022", "growing_season_days_2023", "garden_Arboreta.University.Partner", 
## "garden_City", "garden_State", "garden_Latitude", "garden_Longitude", 
## "garden_Elevation_m", "garden_MAT_1961_1990", "garden_MWMT_1961_1990", 
## "garden_MCMT_1961_1990", "garden_TD_1961_1990", "garden_MAP_1961_1990", 
## "garden_MSP_1961_1990", "garden_AHM_1961_1990", "garden_SHM_1961_1990", 
## "garden_DD_0_1961_1990", "garden_DD5_1961_1990", "garden_DD_18_1961_1990", 
## "garden_DD18_1961_1990", "garden_NFFD_1961_1990", "garden_bFFP_1961_1990", 
## "garden_eFFP_1961_1990", "garden_FFP_1961_1990", "garden_PAS_1961_1990", 
## "garden_EMT_1961_1990", "garden_EXT_1961_1990", "garden_Eref_1961_1990", 
## "garden_CMD_1961_1990", "garden_MAR_1961_1990", "garden_RH_1961_1990", 
## "garden_CMI_1961_1990", "garden_DD1040_1961_1990", "garden_MAT_2020", 
## "garden_MWMT_2020", "garden_MCMT_2020", "garden_TD_2020", "garden_MAP_2020", 
## "garden_MSP_2020", "garden_AHM_2020", "garden_SHM_2020", "garden_DD_0_2020", 
## "garden_DD5_2020", "garden_DD_18_2020", "garden_DD18_2020", "garden_NFFD_2020", 
## "garden_bFFP_2020", "garden_eFFP_2020", "garden_FFP_2020", "garden_PAS_2020", 
## "garden_EMT_2020", "garden_EXT_2020", "garden_Eref_2020", "garden_CMD_2020", 
## "garden_MAR_2020", "garden_RH_2020", "garden_CMI_2020", "garden_DD1040_2020", 
## "garden_MAT_2021", "garden_MWMT_2021", "garden_MCMT_2021", "garden_TD_2021", 
## "garden_MAP_2021", "garden_MSP_2021", "garden_AHM_2021", "garden_SHM_2021", 
## "garden_DD_0_2021", "garden_DD5_2021", "garden_DD_18_2021", "garden_DD18_2021", 
## "garden_NFFD_2021", "garden_bFFP_2021", "garden_eFFP_2021", "garden_FFP_2021", 
## "garden_PAS_2021", "garden_EMT_2021", "garden_EXT_2021", "garden_Eref_2021", 
## "garden_CMD_2021", "garden_MAR_2021", "garden_RH_2021", "garden_CMI_2021", 
## "garden_DD1040_2021", "garden_MAT_2022", "garden_MWMT_2022", 
## "garden_MCMT_2022", "garden_TD_2022", "garden_MAP_2022", "garden_MSP_2022", 
## "garden_AHM_2022", "garden_SHM_2022", "garden_DD_0_2022", "garden_DD5_2022", 
## "garden_DD_18_2022", "garden_DD18_2022", "garden_NFFD_2022", 
## "garden_bFFP_2022", "garden_eFFP_2022", "garden_FFP_2022", "garden_PAS_2022", 
## "garden_EMT_2022", "garden_EXT_2022", "garden_Eref_2022", "garden_CMD_2022", 
## "garden_MAR_2022", "garden_RH_2022", "garden_CMI_2022", "garden_DD1040_2022", 
## "garden_MAT_2023", "garden_MWMT_2023", "garden_MCMT_2023", "garden_TD_2023", 
## "garden_MAP_2023", "garden_MSP_2023", "garden_AHM_2023", "garden_SHM_2023", 
## "garden_DD_0_2023", "garden_DD5_2023", "garden_DD_18_2023", "garden_DD18_2023", 
## "garden_NFFD_2023", "garden_bFFP_2023", "garden_eFFP_2023", "garden_FFP_2023", 
## "garden_PAS_2023", "garden_EMT_2023", "garden_EXT_2023", "garden_Eref_2023", 
## "garden_CMD_2023", "garden_MAR_2023", "garden_RH_2023", "garden_CMI_2023", 
## "garden_DD1040_2023", "licor_Unique_ID", "licor_obs", "licor_gsw", 
## "licor_gbw", "licor_gtw", "licor_E_apparent", "licor_VPcham", 
## "licor_VPref", "licor_VPleaf", "licor_VPDleaf", "licor_H2O_r", 
## "licor_H2O_s", "licor_H2O_leaf", "licor_Fs", "licor_Fm.", "licor_PhiPS2", 
## "licor_PS2.1", "licor_abs", "licor_ETR", "licor_rh_s", "licor_rh_r", 
## "licor_Tref", "licor_Tleaf", "licor_P_atm", "licor_Qamb", "licor_doy", 
## "licor_pitch", "licor_roll", "licor_heading", "licor_angle_inc_leaf", 
## "licor_direct_pct", "licor_slope_leaf", "licor_az_leaf", "licor_dec_solar", 
## "licor_az_solar", "licor_zenith_solar", "licor_latitude", "licor_longitude", 
## "licor_altitude", "licor_gps_sats", "licor_MiniCG_Site_first_measure", 
## "licor_Time_first_measure", "licor_Date_first_measure", "licor_lciSerNum_first_measure", 
## "licor_remark_first_measure", "licor_gps_time_first_measure", 
## "licor_gps_date_first_measure", "licor_Notes_first_measure", 
## "licor_Bud.stage_first_measure", "licor_Shade._first_measure", 
## "licor_Weather.notes_first_measure", "lower_stomata_presence", 
## "lower_fiveOrMoreStomata", "lower_stomata_count", "lower_stomata_pore_length_mean_um", 
## "lower_stomata_pore_length_sd_um", "upper_stomata_presence", 
## "upper_fiveOrMoreStomata", "upper_stomata_count", "upper_stomata_pore_length_mean_um", 
## "upper_stomata_pore_length_sd_um", "upper_stomata_density_mm2", 
## "lower_stomata_density_mm2", "stomata_ratio", "provenance_MAT", 
## "provenance_MWMT", "provenance_MCMT", "provenance_TD", "provenance_MAP", 
## "provenance_MSP", "provenance_AHM", "provenance_SHM", "provenance_DD_0", 
## "provenance_DD5", "provenance_DD_18", "provenance_DD18", "provenance_NFFD", 
## "provenance_bFFP", "provenance_eFFP", "provenance_FFP", "provenance_PAS", 
## "provenance_EMT", "provenance_EXT", "provenance_MAR", "provenance_Eref", 
## "provenance_CMD", "provenance_RH", "ID", "k2_balsam", "k2_tricho", 
## "k3_balsam", "k3_coastal_tricho", "k3_interior_tricho", "genetic_PC1", 
## "genetic_PC2", "genetic_PC3", "genetic_PC4", "genetic_PC5", "genetic_PC6", 
## "interspecific_heterozygosity", "hybrid_index_Pb", "heterozygosity", 
## "plastid_ID", "garden_climate_1961_1990_PC1", "garden_climate_1961_1990_PC2", 
## "garden_climate_1961_1990_PC3", "garden_climate_1961_1990_PC4", 
## "garden_climate_1961_1990_PC5", "garden_climate_1961_1990_PC6", 
## "garden_climate_1961_1990_PC7", "garden_climate_1961_1990_PC8", 
## "garden_climate_1961_1990_PC9", "garden_climate_2020_PC1", "garden_climate_2020_PC2", 
## "garden_climate_2020_PC3", "garden_climate_2020_PC4", "garden_climate_2020_PC5", 
## "garden_climate_2020_PC6", "garden_climate_2020_PC7", "garden_climate_2020_PC8", 
## "garden_climate_2020_PC9", "garden_climate_2021_PC1", "garden_climate_2021_PC2", 
## "garden_climate_2021_PC3", "garden_climate_2021_PC4", "garden_climate_2021_PC5", 
## "garden_climate_2021_PC6", "garden_climate_2021_PC7", "garden_climate_2021_PC8", 
## "garden_climate_2021_PC9", "garden_climate_2022_PC1", "garden_climate_2022_PC2", 
## "garden_climate_2022_PC3", "garden_climate_2022_PC4", "garden_climate_2022_PC5", 
## "garden_climate_2022_PC6", "garden_climate_2022_PC7", "garden_climate_2022_PC8", 
## "garden_climate_2022_PC9", "garden_climate_2023_PC1", "garden_climate_2023_PC2", 
## "garden_climate_2023_PC3", "garden_climate_2023_PC4", "garden_climate_2023_PC5", 
## "garden_climate_2023_PC6", "garden_climate_2023_PC7", "garden_climate_2023_PC8", 
## "garden_climate_2023_PC9", "provenance_climate_PC1", "provenance_climate_PC2", 
## "provenance_climate_PC3", "provenance_climate_PC4", "provenance_climate_PC5", 
## "provenance_climate_PC6", "provenance_climate_PC7", "provenance_climate_PC8", 
## "provenance_climate_PC9", "color_Pt", "color_k3")
```

```
columns <- c("Genotype", "Plant_ID", "Pb", "Pt", "transect", "provenance_latitude", "provenance_longitude", "provenance_elevation_m", "provenance_TD", "provenance_MCMT", "genetic_PC1", "genetic_PC2", "genetic_PC3", "genetic_PC4", "genetic_PC5", "plastid_ID", "color_Pt", "color_k3")

# get genotypes
genos <- unique(dat$Genotype)
genos <- genos[! is.na(genos)]

# get the rows that have the first match of each genotype
rows <- match(genos, dat$Genotype)

# get subset of dataframe with these rows and the info we want for each genotype
geno.info <- dat[rows, columns]
rownames(geno.info) <- geno.info$Genotype

# remove the ones with missing genetic info
geno.info <- geno.info[! is.na(geno.info$genetic_PC1),]
dim(geno.info)
```

```
## [1] 44 18
```

```
str(geno.info)
```

```
## 'data.frame':    44 obs. of  18 variables:
##  $ Genotype              : Factor w/ 47 levels "206","210","218",..: 1 2 3 4 5 6 7 8 9 10 ...
##  $ Plant_ID              : chr  "206" "210" "218" "233" ...
##  $ Pb                    : num  0.4289 0.0916 0.1356 0.0342 0.2851 ...
##  $ Pt                    : num  0.571 0.908 0.864 0.966 0.715 ...
##  $ transect              : Factor w/ 5 levels "Alaska","Cassiar",..: 3 3 3 3 3 3 5 5 5 5 ...
##  $ provenance_latitude   : num  51.9 52.1 52 53.3 55.5 ...
##  $ provenance_longitude  : num  -124 -123 -122 -122 -123 ...
##  $ provenance_elevation_m: num  1064 744 381 677 816 ...
##  $ provenance_TD         : num  22.6 25.5 26.4 24.7 25.7 27.6 19.9 23.4 24.6 23.3 ...
##  $ provenance_MCMT       : num  -9.6 -9.9 -8.9 -9.3 -12.2 -12.4 -6.9 -5.8 -5.8 -4.8 ...
##  $ genetic_PC1           : num  0.00423 -0.03207 -0.02393 -0.03496 -0.00995 ...
##  $ genetic_PC2           : num  0.02407 -0.00722 0.01811 0.039 0.03374 ...
##  $ genetic_PC3           : num  -0.0243 -0.0365 -0.0505 -0.0418 -0.0256 ...
##  $ genetic_PC4           : num  0.0378 0.0122 0.046 0.0279 0.0399 ...
##  $ genetic_PC5           : num  0.00732 0.02208 0.01169 -0.02046 -0.00303 ...
##  $ plastid_ID            : chr  "balsamifera" "trichocarpa" "balsamifera" "balsamifera" ...
##  $ color_Pt              : chr  "#464B3CFF" "#A0C65FFF" "#94B55AFF" "#AFDB64FF" ...
##  $ color_k3              : chr  "#1B8163" "#836913" "#4E941D" "#39C600" ...
```

### 2 Get future climates for each genotype

#### 2.1 Load climate data

Load future and historic MCMT raster files

ClimateNA raster files used here are available from DataBasin:

https://adaptwest.databasin.org/pages/adaptwest-climatena/

Species range shapefiles are from Little 1971 and are available from
DataBasin:

P. balsamifera: https://databasin.org/datasets/91380e091ca048359a66fc65962ed210/

P. trichocarpa: https://databasin.org/datasets/84e47784fe2a463c8b292007fb43f2cd/

```
# which climate variable is being used? 
# can change this to use a different variable
clim_var <- 'MCMT'

# get historic and future climate

hist <- rast(paste('data/climate/climateNA/Normal_1961_1990/Normal_1961_1990_bioclim/Normal_1961_1990_', clim_var, '.tif', sep = ''))
crs(hist, proj = T)
```

```
## [1] "+proj=laea +lat_0=45 +lon_0=-100 +x_0=0 +y_0=0 +datum=WGS84 +units=m +no_defs"
```

```
hist91 <- rast(paste('data/climate/climateNA/Normal_1991_2020/Normal_1991_2020_', clim_var, '.tif', sep = ''))
crs(hist, proj = T)
```

```
## [1] "+proj=laea +lat_0=45 +lon_0=-100 +x_0=0 +y_0=0 +datum=WGS84 +units=m +no_defs"
```

```
# climateNA provides future ensemble models built from 8 GCMs and 13 GCMs - test both

fut8 <- rast(paste('data/climate/climateNA/future/ensemble_8GCMs_ssp245_2041_2070_bioclim/ensemble_8GCMs_ssp245_2041_2070_', clim_var, '.tif', sep = ''))

fut13 <- rast(paste('data/climate/climateNA/future/ensemble_13GCMs_ssp245_2041_2070_bioclim/ensemble_13GCMs_ssp245_2041_2070_', clim_var, '.tif', sep = ''))

################ convert coords

# convert coordinates of collection sites to the CRS used by climateNA
crs.cna <- crs(hist, proj = T)

# provenance coordinates
coords.prov <- st_as_sf(geno.info[,c("provenance_longitude", "provenance_latitude")], coords = c(1,2), crs = st_crs(4326))
coords.prov.cna <- st_transform(coords.prov$geometry, crs = crs.cna)

# load shapefile of state/province borders
borders <- read_sf('data/shapefiles/NorthAmerica_PoliticalBoundaries_Shapefile/NA_PoliticalDivisions/data/bound_p/boundaries_p_2021_v3.shp')
borders <- st_transform(borders, crs = crs.cna)
# simplify to 1 km
borders <- st_simplify(borders, dTolerance = 1000)

# species rangemaps
# shapefile
balsam <- st_read('data/shapefiles/Pbal_shapefile/data/commondata/data0/popubals.shp')
```

```
## Reading layer `popubals' from data source 
##   `/home/alayna/Documents/research/projects/2023_populus_common_gardens/data/shapefiles/Pbal_shapefile/data/commondata/data0/popubals.shp' 
##   using driver `ESRI Shapefile'
## Simple feature collection with 415 features and 5 fields
## Geometry type: POLYGON
## Dimension:     XY
## Bounding box:  xmin: -18247640 ymin: 4662920 xmax: -5857014 ymax: 10826350
## Projected CRS: WGS 84 / Pseudo-Mercator
```

```
tricho <- st_read('data/shapefiles/Ptri_shapefile/data/commondata/data0/poputric.shp')
```

```
## Reading layer `poputric' from data source 
##   `/home/alayna/Documents/research/projects/2023_populus_common_gardens/data/shapefiles/Ptri_shapefile/data/commondata/data0/poputric.shp' 
##   using driver `ESRI Shapefile'
## Simple feature collection with 450 features and 5 fields
## Geometry type: POLYGON
## Dimension:     XY
## Bounding box:  xmin: -17154180 ymin: 3609620 xmax: -11488690 ymax: 8896231
## Projected CRS: WGS 84 / Pseudo-Mercator
```

```
# convert CRS
balsam <- st_transform(balsam, crs.cna)
tricho <- st_transform(tricho, crs.cna)

# test plot
plot(hist)

plot(balsam$geometry, add = T, border = 'navy')
plot(tricho$geometry, add = T, border = 'grey50')
plot(borders$geometry, add = T)
```

#### 2.2 Extract future climates for each genotype’s home site

```
# extract from raster and add to geno.info

# 8 GCM ensemble
tmp <- extract(fut8, y = st_as_sf(coords.prov))
```

```
## Warning: [extract] transforming vector data to the CRS of the raster
```

```
geno.info$future8_MCMT <- tmp[,2]

# 13 GCM ensemble
tmp <- extract(fut13, y = st_as_sf(coords.prov))
```

```
## Warning: [extract] transforming vector data to the CRS of the raster
```

```
geno.info$future13_MCMT <- tmp[,2]

# predictions from the two ensembles are very similar
plot(geno.info$future8_MCMT, geno.info$future13_MCMT)
abline(0, 1, col = 'blue')
```

### 3 Predict height at future climates

#### 3.1 Predict functions

```
# originally from transfer function script

# predict each genotype's response to a vector of climate (MCMT) values
# garden_clims is a vector

predict_genotype <- function(model, 
                             type = 'response', # can also be conditional or zprob
                             effects,
                             home_clim,
                             garden_clims){
  
  # vector to save predictions
  pred_height <- vector()
  
  # loop through garden climates
  for(g in 1:length(garden_clims)){
    
    home <- home_clim
    garden <- garden_clims[g]
    
    # make newdata to give to predict()
    new_data <- as.data.frame(cbind(home, garden, home^2, garden^2))
    colnames(new_data) <- c('home_clim', 'garden_clim', 'home_clim_2', 'garden_clim_2')
    # add other random or fixed effects
      for(n in 1:length(effects)){
        new_data[names(effects[n])] <- effects[n]
      }
    
    # first have to scale new_data to match the scaled data used in model
    for(v in 1:length(new_data)){
      var <- names(new_data)[v]
      
      # only scale if var exists in scaling_factor - random effects like genotype are not in scaling factor
      # IF VARIABLES ARE NAMED DIFFERENTLY HERE IN effects AND scaling_factor IT WILL SKIP THEM
      if(var %in% names(scaling_factor)){
        new_data[var] <- new_data[var]*scaling_factor[[var]]
      } else {
        warning(var, ' does not have a scaling factor, skipping')
      }
      
      
    }
    
    pred <- predict(mod, type = type, newdata = new_data, re.form = NA, allow.new.levels=TRUE)
    # back transform from log if we're predicting height
    # zprob is probability of a zero, so don't transform
    if(type %in% c('response', 'conditional')){
      pred <- exp(pred)-1
    }
    pred_height[g] <- pred
    
  }
  
  # save
  pred_height_df <- data.frame(garden_clim = garden_clims, predicted_height = pred_height) 
  
return(pred_height_df)
}
```

#### 3.2 Predict response to historic and future climate for each genotype

```
# model predictions for historic data
# we already have historic provenance data for the model, so don't need to extract it from the raster

# what climate variable are we using?
clim_var <- 'MCMT'
prov_clim_colname <- paste('provenance', clim_var, sep = '_')
gard_clim_colname <- paste('garden', clim_var, '1961_1990', sep = '_')
fut8_colname <- paste('future8', clim_var, sep = '_')
fut13_colname <- paste('future13', clim_var, sep = '_')


# add predictions to geno.info dataframe
# historic
geno.info$hist_response <- NA
geno.info$hist_height <- NA
geno.info$hist_mortality <- NA

# future 8
geno.info$fut8_response <- NA
geno.info$fut8_height <- NA
geno.info$fut8_mortality <- NA

# future 13
geno.info$fut13_response <- NA
geno.info$fut13_height <- NA
geno.info$fut13_mortality <- NA


for(n in 1:nrow(geno.info)){
  
  geno <- rownames(geno.info)[n]
  
  # setup effects
  g.pc1 <- geno.info$genetic_PC1[n]
  g.pc2 <- geno.info$genetic_PC2[n]
  g.pc3 <- geno.info$genetic_PC3[n]
  g.home_clim <- geno.info[,prov_clim_colname][n]
  
  # garden clims are the historic and future climates
  # in order: home climate (historic), future climate 8 GCMs, future climate 13 GCMs
  g.clims <- c(g.home_clim, geno.info[,fut8_colname][n], geno.info[,fut13_colname][n])
  
  # predict overall response
  resp <- predict_genotype(mod,
                   type = 'response',
                   effects = list(pc1 = g.pc1, 
                                  pc2 = g.pc2,
                                  pc3 = g.pc3,
                                  genotype = geno), 
                   home_clim = g.home_clim, 
                   garden_clims = g.clims)
  
  # save to df
  geno.info$hist_response[n] <- resp$predicted_height[1]
  geno.info$fut8_response[n] <- resp$predicted_height[2]
  geno.info$fut13_response[n] <- resp$predicted_height[3]

  
  
  # predict height (conditional model)
  cond <- predict_genotype(mod,
                           type = 'conditional',
                           effects = list(pc1 = g.pc1, 
                                          pc2 = g.pc2,
                                          pc3 = g.pc3,
                                          genotype = geno), 
                           home_clim = g.home_clim, 
                           garden_clims = g.clims)
  
  # save to df
  geno.info$hist_height[n] <- cond$predicted_height[1]
  geno.info$fut8_height[n] <- cond$predicted_height[2]
  geno.info$fut13_height[n] <- cond$predicted_height[3]

  # predict probability of mortality (zero-inflated model)
  zi <- predict_genotype(mod,
                         type = 'zprob',
                         effects = list(pc1 = g.pc1, 
                                        pc2 = g.pc2,
                                        pc3 = g.pc3,
                                        genotype = geno), 
                         home_clim = g.home_clim, 
                         garden_clims = g.clims)
  # save to df
  geno.info$hist_mortality[n] <- zi$predicted_height[1]
  geno.info$fut8_mortality[n] <- zi$predicted_height[2]
  geno.info$fut13_mortality[n] <- zi$predicted_height[3]

  
 #cat(paste('done with', n, '\n'))
  
}

# predict_genotype() gives warning:
# '## Warning in predict_genotype(mod, type = "response", effects = list(pc1 = g.pc1,
## : genotype does not have a scaling factor, skipping'

# this is fine, genotype is a string not numeric and shouldn't be scaled
```

#### 3.3 Calculate change in fitness under future climates

```
# calculate changes in fitness under future climates

##########
# future (8 GCMs)

# net change in response
geno.info$fut8_response_change <- geno.info$fut8_response - geno.info$hist_response
# percent change
geno.info$fut8_response_change_percent <- (geno.info$fut8_response - geno.info$hist_response)/geno.info$hist_response

# net change in height
geno.info$fut8_height_change <- geno.info$fut8_height - geno.info$hist_height
# percent change
geno.info$fut8_height_change_percent <- (geno.info$fut8_height - geno.info$hist_height)/geno.info$hist_height

# net change in mortality probability
geno.info$fut8_mortality_change <- geno.info$fut8_mortality - geno.info$hist_mortality
# percent change
geno.info$fut8_mortality_change_percent <- (geno.info$fut8_mortality - geno.info$hist_mortality)/geno.info$hist_mortality

##########
# future (13 GCMs)

# net change in response
geno.info$fut13_response_change <- geno.info$fut13_response - geno.info$hist_response
# percent change
geno.info$fut13_response_change_percent <- (geno.info$fut13_response - geno.info$hist_response)/geno.info$hist_response

# net change in height
geno.info$fut13_height_change <- geno.info$fut13_height - geno.info$hist_height
# percent change
geno.info$fut13_height_change_percent <- (geno.info$fut13_height - geno.info$hist_height)/geno.info$hist_height

# net change in mortality probability
geno.info$fut13_mortality_change <- geno.info$fut13_mortality - geno.info$hist_mortality
# percent change
geno.info$fut13_mortality_change_percent <- (geno.info$fut13_mortality - geno.info$hist_mortality)/geno.info$hist_mortality

# preliminary plots
par(mfrow = c(2,3))
hist(geno.info$fut8_response_change_percent)
hist(geno.info$fut8_height_change_percent)
hist(geno.info$fut8_mortality_change_percent)

hist(geno.info$fut13_response_change_percent)
hist(geno.info$fut13_height_change_percent)
hist(geno.info$fut13_mortality_change_percent)
```

### 4 Map predictions for each collected genotype/site

```
# baseline plot of climate difference

diff13 <- fut13 - hist
diff8 <- fut8 - hist

par(mfrow = c(1,2))
plot(diff13)
plot(diff8)
```

```
# color palette for MCMT base layer

colf <- colorRampPalette(brewer.pal(9, 'YlOrRd'))
cols <- colf(100)
cols2 <- adjustcolor(cols, alpha = 0.5)

# function to map the response 

# future = the raster object for change in climate to use as background map
# response = the column name from geno.info of the response to future climate to plot
# main = plot title
# rev = reverse response color palette? Can use for mortality so a negative/positive response has the same color
map_change <- function(future, response, main = '', rev = FALSE, legend = FALSE, raster = FALSE, ...){
  
  # which model response are we plotting?
  resp_name <- response
  
  colf <- colorRampPalette(brewer.pal(9, 'YlOrRd'), alpha = 0.5)
  cols <- colf(100)
  cols <- adjustcolor(cols, alpha = 0.6)
  
  #par(mfrow = c(1,1))
  plot(future, 
       col = ifelse(raster == TRUE, cols, 'white'),
       main = main,
       xaxt = 'n', yaxt = 'n', 
       pax = list(side=NA),
       xlim = c(-3000000,-500000), ylim = c(400000,3200000), 
       legend = legend,
       plg = list(title = 'Future -\nHistoric\n MCMT (°C)'),
       ...)
  
  # add borders and species ranges
  plot(borders$geometry, add = T, lwd = 0.5)
  plot(balsam$geometry, add = T, border = 'navy')
  plot(tricho$geometry, add = T, border = 'darkgreen')
  

  # color palette for change in response
  min_change <- min(geno.info[,resp_name])
  max_change <- max(geno.info[,resp_name])
  
  ceil2 <- max(abs(min_change), abs(max_change))
  
  if(rev == FALSE){
    colf <- colorRamp2( breaks = c(-ceil2, 0, ceil2), colors = c('darkorchid4', 'white', 'palegreen4'))
  } else if(rev == TRUE){
    colf <- colorRamp2( breaks = c(-ceil2, 0, ceil2), colors = c('palegreen4', 'white', 'darkorchid4'))
  }
  
  
  cols2 <- colf(geno.info[,resp_name])
  
  # plot points for each genotype
  points(coords.prov.cna, bg = cols2,  pch = 21, cex = 2, lwd = 1.5)
  
  # add colors of upper and lower bounds to legend
  if(rev == FALSE){
     pt.cols <- c('darkorchid4', 'white', 'palegreen4')
  } else if(rev == TRUE) {
    pt.cols = c('palegreen4', 'white', 'darkorchid4')
  }
 
  
  # legend(-2.8e6,1.5e6, title = '% Change',
  #        legend = c(round(-ceil2,2), 0, round(ceil2,2)),
  #        pch = 21,
  #        pt.bg = pt.cols,
  #        pt.cex = 2)
  
  vals <- round(c(min_change, 0, max_change),2)
  # if range doesn't overlap 0, remove it
  if(min_change * max_change > 0){
    vals <- vals[vals!=0]
  }
  
  legend(-2.8e6,1.5e6, title = '% Change',
         legend = vals,
         pch = 21,
         pt.bg = colf(vals),
         pt.cex = 2)
  
  
}


par(mfrow = c(2,3))
map_change(diff13, 'fut13_response_change_percent')
map_change(diff13, 'fut13_height_change_percent')
map_change(diff13, 'fut13_mortality_change_percent', rev = T)

map_change(diff8, 'fut8_response_change_percent')
map_change(diff8, 'fut8_height_change_percent')
map_change(diff8, 'fut8_mortality_change_percent', rev = T)
```

```
# raw values (cm or % mortality)
# legend title not correct here
par(mfrow = c(2,3))
map_change(diff13, 'fut13_response_change')
map_change(diff13, 'fut13_height_change')
map_change(diff13, 'fut13_mortality_change', rev = T)

map_change(diff8, 'fut8_response_change')
map_change(diff8, 'fut8_height_change')
map_change(diff8, 'fut8_mortality_change', rev = T)
```

```
#png(file = paste('results/model_prediction/fitness_changes_map_2yearModel', clim_var, '_13GCMs.png', sep = ''), height = 8, width = 8, res = 300, units = 'in')


# lighter colors for temperature raster
colf <- colorRampPalette(brewer.pal(9, 'YlOrRd'))
cols <- colf(100)
cols2 <- adjustcolor(cols, alpha = 0.5)

# plot genotypes and the 13-gcm ensemble
par(mfrow = c(2,2))
# set margins
marg <- c(1,1,4,3)

plot(diff13, col = cols2, 
     main = paste(clim_var, ' 2041-2070, 13 GCMs', sep = ''),
     xaxt = 'n', yaxt = 'n', 
     pax = list(side=NA),
     mar = marg,
     xlim = c(-3000000,-500000), ylim = c(400000,3200000), 
     plg = list(title = 'Future -\nHistoric\n MCMT (°C)'))
# add borders and species ranges
plot(borders$geometry, add = T, lwd = 0.5)
plot(balsam$geometry, add = T, border = 'navy')
plot(tricho$geometry, add = T, border = 'darkgreen')

points(coords.prov.cna, bg = geno.info$color_Pt, pch = 21, cex = 2)

map_change(diff13, 'fut13_response_change_percent', main = 'Overall response', mar = marg)
map_change(diff13, 'fut13_height_change_percent', main = 'Growth', mar = marg)
map_change(diff13, 'fut13_mortality_change_percent', rev = T, main = 'Probability of Mortality', mar = marg)
```

```
#dev.off()
```

### 5 Map rangewide predictions for each genotype

#### 5.1 Map past and future fitness predictions and change for each genotype

```
# custom predict function - predict height from discrete set of climate values that have already been calculated

predict_disc <- function(value, pred_clim, pred_value){
  if(! is.na(value)){
    # which value in pred_clim (already predicted climates) is closest to the value we want to predict?
    # returns the index
    ind <- which.min(abs(pred_clim - value))
    
    # return the predicted height for that index
    return(pred_value[ind])
    
  } else if(is.na(value)){
    return(NA)
  }
}

# which climate variable to use?
clim_var <- 'MCMT'
hist_rast <- hist
fut_rast <- fut13

# set up input raster - historic climate
hist.crop <- crop(hist_rast, ext(-4e6, 0, 0, 3e6))
plot(hist.crop)
```

```
# make values outside predict range NA
# need pred_clim - this is the same for all genotypes, so can just use first one
#pred_clim <- preds$overall$genotype_210$garden_clim
#pred_clim <- dat[,gard_clim_colname]
pred_clim <- c(dat$garden_MCMT_2021, dat$garden_MCMT_2022)

msk <- ifel(hist.crop > max(pred_clim) | hist.crop < min(pred_clim), NA, 1)
hist.crop <- mask(hist.crop, msk)
plot(hist.crop, colNA = 'grey')
```

```
# future climate
fut13.crop <- crop(fut_rast, ext(-4e6, 0, 0, 3e6))
plot(fut13.crop)
```

```
# make values outside predict range NA
# need pred_clim - this is the same for all genotypes, so can just use first one
# pred_clim <- preds$overall$genotype_210$garden_clim
msk <- ifel(fut13.crop > max(pred_clim) | fut13.crop < min(pred_clim), NA, 1)
fut13.crop <- mask(fut13.crop, msk)
plot(fut13.crop, colNA = 'grey')
```

```
# loop through genotypes, predict height across range

par(mfrow = c(1,3), mar = c(5,4,4,5))
for(n in 1:length(preds$overall)){
  
  # predicted climate and predicted height for this genotype
  pred_clim <- preds$overall[[n]]$garden_clim
  pred_value <- preds$overall[[n]]$predicted_height
  
  #### historic
  # set heights for each raster cell in historic climate (returns a list)
  heights_cells_hist <- lapply(values(hist.crop), FUN = function(x) predict_disc(x, pred_clim, pred_value))
  # assign heights to raster
  height_hist <- setValues(hist.crop, values = unlist(heights_cells_hist))
  
  # plot
   colf <- colorRampPalette(rev(brewer.pal(9, 'YlGn')))
   cols <- rev(colf(100))
  
  plot(height_hist, col = cols, main = paste(names(preds$overall)[n], '\nFitness in Historic Climate', sep = ''), colNA = 'grey90')
  
  plot(borders$geometry, add = T)
  plot(balsam$geometry, add = T, border = 'navy')
  plot(tricho$geometry, add = T, border = 'grey50')
  
  points(coords.prov.cna[n], col = 'mediumpurple', lwd = 2, cex = 1.5)
  #points(coords.prov.cna)
  
    #### future
  # set heights for each raster cell in historic climate (returns a list)
  heights_cells_fut <- lapply(values(fut13.crop), FUN = function(x) predict_disc(x, pred_clim, pred_value))
  # assign heights to raster
  height_fut <- setValues(fut13.crop, values = unlist(heights_cells_fut))
  
  # plot

  plot(height_fut, col = cols, main = 'Fitness in Future Climate', colNA = 'grey90')
  
  plot(borders$geometry, add = T)
  plot(balsam$geometry, add = T, border = 'navy')
  plot(tricho$geometry, add = T, border = 'grey50')
  
  points(coords.prov.cna[n], col = 'mediumpurple', lwd = 2, cex = 1.5)
  #points(coords.prov.cna)
  
  ### plot difference
  height_diff <- height_fut - height_hist
  
  # min and max values for diverging color palette
  min_var <- minmax(height_diff)[1,]
  max_var <- minmax(height_diff)[2,]
  ceil <- max(abs(min_var), abs(max_var))
  
  colf_diff <- colorRampPalette(brewer.pal(11, 'RdBu'))
  cols_diff <- colf_diff(100)
  
  plot(height_diff, col = cols_diff, range = c(-ceil, ceil), main = 'Fitness change', colNA = 'grey90')
  
  plot(borders$geometry, add = T)
  plot(balsam$geometry, add = T, border = 'navy')
  plot(tricho$geometry, add = T, border = 'grey50')
  
  points(coords.prov.cna[n], col = 'black', lwd = 2, cex = 1.5)
  
  print(paste('done with', n))
  
}
```

```
## [1] "done with 1"
```

```
## [1] "done with 2"
```

```
## [1] "done with 3"
```

```
## [1] "done with 4"
```

```
## [1] "done with 5"
```

```
## [1] "done with 6"
```

```
## [1] "done with 7"
```

```
## [1] "done with 8"
```

```
## [1] "done with 9"
```

```
## [1] "done with 10"
```

```
## [1] "done with 11"
```

```
## [1] "done with 12"
```

```
## [1] "done with 13"
```

```
## [1] "done with 14"
```

```
## [1] "done with 15"
```

```
## [1] "done with 16"
```

```
## [1] "done with 17"
```

```
## [1] "done with 18"
```

```
## [1] "done with 19"
```

```
## [1] "done with 20"
```

```
## [1] "done with 21"
```

```
## [1] "done with 22"
```

```
## [1] "done with 23"
```

```
## [1] "done with 24"
```

```
## [1] "done with 25"
```

```
## [1] "done with 26"
```

```
## [1] "done with 27"
```

```
## [1] "done with 28"
```

```
## [1] "done with 29"
```

```
## [1] "done with 30"
```

```
## [1] "done with 31"
```

```
## [1] "done with 32"
```

```
## [1] "done with 33"
```

```
## [1] "done with 34"
```

```
## [1] "done with 35"
```

```
## [1] "done with 36"
```

```
## [1] "done with 37"
```

```
## [1] "done with 38"
```

```
## [1] "done with 39"
```

```
## [1] "done with 40"
```

```
## [1] "done with 41"
```

```
## [1] "done with 42"
```

```
## [1] "done with 43"
```

```
## [1] "done with 44"
```
