## Supplementary material for "Variation in responses to temperature in admixed *Populus* genotypes predicts geographic shifts in regions where hybrids are favored": Rmarkdown output files: predict_maxi_garden_phenotypes.html

Predict height in maxis


### Predict height in maxis

###### Alayna Mead

#### 2025-09-23

- 1 Setup
  - 1.1 packages and data
- 2 clean up data
  - 2.1 change ND name
  - 2.2 set dead to 0
  - 2.3 remove negative heights
- 3 Set up model
  - 3.1 choose model variables
  - 3.2 scale variables
- 4 Predict growth and mortality from
  model
  - 4.1 Functions
  - 4.2 Predict for maxi garden
    data

Test whether model built from mini gardens dataset can accurately
predict growth in the maxi gardens (3 sites, one of which is novel, and
544 genotypes, 500 of which are novel)

This code is mostly pulled from
‘code/transfer\_function\_multiyear\_linear\_mixed\_effects\_model.Rmd’
script

### 1 Setup

#### 1.1 packages and data

```
library(ggplot2)
library(sjPlot) # nice plots of models
```

```
## Learn more about sjPlot with 'browseVignettes("sjPlot")'.
```

```
library(glmmTMB) #zero-inflated  model and also used with sjPlot
library(bbmle) # for AICtab
```

```
## Loading required package: stats4
```

```
library(patchwork) # for wrap_plots

# maxi garden data - named 'maxis'
load('data/clean/maxi_garden_phenotypic_climate_genetic_data_2021-2023.Rdata')
str(maxis)
```

```
## 'data.frame':    4928 obs. of  276 variables:
##  $ ID                                 : chr  "ND.201.1.5.1" "ND.201.2.19.8" "ND.201.3.35.24" "ND.202.1.6.43" ...
##  $ garden                             : chr  "ND" "ND" "ND" "ND" ...
##  $ TAG                                : chr  "201" "201" "201" "202" ...
##  $ PLANT_ID                           : chr  "201" "201" "201" "202" ...
##  $ CORRECTED_PLANT_ID                 : chr  NA NA NA NA ...
##  $ TRANSECT                           : chr  "Chilcotin" "Chilcotin" "Chilcotin" "Chilcotin" ...
##  $ BLOCK                              : chr  "1" "2" "3" "1" ...
##  $ ROW                                : int  5 19 35 6 9 25 5 10 24 5 ...
##  $ COLUMN                             : chr  "1" "8" "24" "43" ...
##  $ LIVE_Y_N_2021                      : num  0 0 0 0 0 0 0 0 0 1 ...
##  $ Budflush2021                       : int  NA 131 124 NA 124 124 NA NA NA 106 ...
##  $ first_budset2021                   : int  NA NA NA NA NA NA NA NA NA 194 ...
##  $ Budset2021                         : int  NA NA NA NA NA NA NA NA NA NA ...
##  $ initial_height2021                 : num  32.31 29.87 12.19 7.01 6.1 ...
##  $ scar_height2021                    : num  NA 9.53 0 NA 8.89 ...
##  $ ground_height2021                  : num  NA 38.1 36.8 NA 16.5 ...
##  $ lammasDuration2021                 : int  NA NA NA NA NA NA NA NA NA NA ...
##  $ ground_Diff2021                    : num  NA 8.23 24.64 NA 10.41 ...
##  $ Budflush2021_cGDD                  : num  NA 348 299 NA 299 ...
##  $ LIVE_Y_N_2022                      : num  NA NA NA NA NA NA NA NA NA NA ...
##  $ scar_height2022                    : num  NA NA NA NA NA NA NA NA NA NA ...
##  $ ground_height2022                  : num  NA NA NA NA NA NA NA NA NA NA ...
##  $ initial_height2022                 : num  NA NA NA NA NA NA NA NA NA NA ...
##  $ lammasDuration2022                 : int  NA NA NA NA NA NA NA NA NA NA ...
##  $ ground_Diff2022                    : num  NA NA NA NA NA NA NA NA NA NA ...
##  $ Budflush2022_cGDD                  : num  NA NA NA NA NA NA NA NA NA NA ...
##  $ Budflush2022                       : int  NA NA NA NA NA NA NA NA NA NA ...
##  $ first_budset2022                   : int  NA NA NA NA NA NA NA NA NA NA ...
##  $ Budset2022                         : int  NA NA NA NA NA NA NA NA NA NA ...
##  $ Height_flagged                     : chr  NA NA NA NA ...
##  $ Petiole_color_2021                 : chr  NA NA NA NA ...
##  $ Stage2_2022                        : int  NA NA NA NA NA NA NA NA NA NA ...
##  $ Stage3_2022                        : int  NA NA NA NA NA NA NA NA NA NA ...
##  $ Stage6_2022                        : int  NA NA NA NA NA NA NA NA NA NA ...
##  $ Stage7_2022                        : int  NA NA NA NA NA NA NA NA NA NA ...
##  $ FinalBudset2022                    : int  NA NA NA NA NA NA NA NA NA NA ...
##  $ Scar_height_to_term_bud_2022       : num  NA NA NA NA NA NA NA NA NA NA ...
##  $ Height_flagged_2022                : chr  NA NA NA NA ...
##  $ LIVE_Y_N_2023                      : num  NA NA NA NA NA NA NA NA NA NA ...
##  $ Budflush2023                       : int  NA NA NA NA NA NA NA NA NA NA ...
##  $ first_budset2023                   : int  NA NA NA NA NA NA NA NA NA NA ...
##  $ Budset2023                         : int  NA NA NA NA NA NA NA NA NA NA ...
##  $ initial_height2023                 : num  NA NA NA NA NA NA NA NA NA NA ...
##  $ scar_height2023                    : num  NA NA NA NA NA NA NA NA NA NA ...
##  $ ground_height2023                  : num  NA NA NA NA NA NA NA NA NA NA ...
##  $ Height_notes_2023                  : chr  NA NA NA NA ...
##  $ lammasDuration2023                 : int  NA NA NA NA NA NA NA NA NA NA ...
##  $ ground_Diff2023                    : num  NA NA NA NA NA NA NA NA NA NA ...
##  $ Scar_height_to_ground_height2023   : num  NA NA NA NA NA NA NA NA NA NA ...
##  $ Budflush2023_cGDD                  : num  NA NA NA NA NA NA NA NA NA NA ...
##  $ stem_notes                         : chr  NA NA NA NA ...
##  $ branch_height20211                 : num  NA NA NA NA NA ...
##  $ branch_increm2021                  : num  NA NA NA NA NA ...
##  $ garden_Latitude                    : num  46.9 46.9 46.9 46.9 46.9 ...
##  $ garden_Longitude                   : num  -96.8 -96.8 -96.8 -96.8 -96.8 ...
##  $ garden_Elevation_m                 : int  272 272 272 272 272 272 272 272 272 272 ...
##  $ garden_MAT_1961_1990               : num  4.9 4.9 4.9 4.9 4.9 4.9 4.9 4.9 4.9 4.9 ...
##  $ garden_MWMT_1961_1990              : num  21.8 21.8 21.8 21.8 21.8 21.8 21.8 21.8 21.8 21.8 ...
##  $ garden_MCMT_1961_1990              : num  -14.3 -14.3 -14.3 -14.3 -14.3 -14.3 -14.3 -14.3 -14.3 -14.3 ...
##  $ garden_TD_1961_1990                : num  36.1 36.1 36.1 36.1 36.1 36.1 36.1 36.1 36.1 36.1 ...
##  $ garden_MAP_1961_1990               : int  544 544 544 544 544 544 544 544 544 544 ...
##  $ garden_MSP_1961_1990               : int  358 358 358 358 358 358 358 358 358 358 ...
##  $ garden_AHM_1961_1990               : num  27.4 27.4 27.4 27.4 27.4 27.4 27.4 27.4 27.4 27.4 ...
##  $ garden_SHM_1961_1990               : num  60.7 60.7 60.7 60.7 60.7 60.7 60.7 60.7 60.7 60.7 ...
##  $ garden_DD_0_1961_1990              : int  1379 1379 1379 1379 1379 1379 1379 1379 1379 1379 ...
##  $ garden_DD5_1961_1990               : int  2144 2144 2144 2144 2144 2144 2144 2144 2144 2144 ...
##  $ garden_DD_18_1961_1990             : int  5032 5032 5032 5032 5032 5032 5032 5032 5032 5032 ...
##  $ garden_DD18_1961_1990              : int  297 297 297 297 297 297 297 297 297 297 ...
##  $ garden_NFFD_1961_1990              : int  162 162 162 162 162 162 162 162 162 162 ...
##  $ garden_bFFP_1961_1990              : int  134 134 134 134 134 134 134 134 134 134 ...
##  $ garden_eFFP_1961_1990              : int  269 269 269 269 269 269 269 269 269 269 ...
##  $ garden_FFP_1961_1990               : int  135 135 135 135 135 135 135 135 135 135 ...
##  $ garden_PAS_1961_1990               : int  85 85 85 85 85 85 85 85 85 85 ...
##  $ garden_EMT_1961_1990               : num  -40.9 -40.9 -40.9 -40.9 -40.9 -40.9 -40.9 -40.9 -40.9 -40.9 ...
##  $ garden_EXT_1961_1990               : num  40.8 40.8 40.8 40.8 40.8 40.8 40.8 40.8 40.8 40.8 ...
##  $ garden_MAR_1961_1990               : num  12.7 12.7 12.7 12.7 12.7 12.7 12.7 12.7 12.7 12.7 ...
##  $ garden_Eref_1961_1990              : int  722 722 722 722 722 722 722 722 722 722 ...
##  $ garden_CMD_1961_1990               : int  273 273 273 273 273 273 273 273 273 273 ...
##  $ garden_RH_1961_1990                : int  58 58 58 58 58 58 58 58 58 58 ...
##  $ garden_CMI_1961_1990               : num  -7.77 -7.77 -7.77 -7.77 -7.77 -7.77 -7.77 -7.77 -7.77 -7.77 ...
##  $ garden_DD1040_1961_1990            : int  1200 1200 1200 1200 1200 1200 1200 1200 1200 1200 ...
##  $ garden_ID2_2020                    : chr  "ND1" "ND1" "ND1" "ND1" ...
##  $ garden_MAT_2020                    : num  5.6 5.6 5.6 5.6 5.6 5.6 5.6 5.6 5.6 5.6 ...
##  $ garden_MWMT_2020                   : num  22.7 22.7 22.7 22.7 22.7 22.7 22.7 22.7 22.7 22.7 ...
##  $ garden_MCMT_2020                   : num  -11.5 -11.5 -11.5 -11.5 -11.5 -11.5 -11.5 -11.5 -11.5 -11.5 ...
##  $ garden_TD_2020                     : num  34.1 34.1 34.1 34.1 34.1 34.1 34.1 34.1 34.1 34.1 ...
##  $ garden_MAP_2020                    : int  702 702 702 702 702 702 702 702 702 702 ...
##  $ garden_MSP_2020                    : int  536 536 536 536 536 536 536 536 536 536 ...
##  $ garden_AHM_2020                    : num  22.2 22.2 22.2 22.2 22.2 22.2 22.2 22.2 22.2 22.2 ...
##  $ garden_SHM_2020                    : num  42.3 42.3 42.3 42.3 42.3 42.3 42.3 42.3 42.3 42.3 ...
##  $ garden_DD_0_2020                   : int  1080 1080 1080 1080 1080 1080 1080 1080 1080 1080 ...
##  $ garden_DD5_2020                    : int  2133 2133 2133 2133 2133 2133 2133 2133 2133 2133 ...
##  $ garden_DD_18_2020                  : int  4866 4866 4866 4866 4866 4866 4866 4866 4866 4866 ...
##  $ garden_DD18_2020                   : int  373 373 373 373 373 373 373 373 373 373 ...
##  $ garden_NFFD_2020                   : int  157 157 157 157 157 157 157 157 157 157 ...
##  $ garden_bFFP_2020                   : int  129 129 129 129 129 129 129 129 129 129 ...
##  $ garden_eFFP_2020                   : int  264 264 264 264 264 264 264 264 264 264 ...
##  $ garden_FFP_2020                    : int  135 135 135 135 135 135 135 135 135 135 ...
##  $ garden_PAS_2020                    : int  137 137 137 137 137 137 137 137 137 137 ...
##   [list output truncated]
```

```
# load model info
# mod: model
# df.scaled: scaled mini garden data that went into the model
# scaling_factor: used to scale variables before fitting model
load('results/model_prediction/glmTMB_multiyear_model_outputs_garden_MCMT_2021-2022_vs_GrowthIncrement_2021-2022.Rda')

# print session info
sessionInfo()
```

```
## R version 4.5.1 (2025-06-13)
## Platform: x86_64-pc-linux-gnu
## Running under: Arch Linux
## 
## Matrix products: default
## BLAS:   /usr/lib/libblas.so.3.12.0 
## LAPACK: /usr/lib/liblapack.so.3.12.0  LAPACK version 3.12.0
## 
## locale:
##  [1] LC_CTYPE=en_US.UTF-8       LC_NUMERIC=C              
##  [3] LC_TIME=en_US.UTF-8        LC_COLLATE=en_US.UTF-8    
##  [5] LC_MONETARY=en_US.UTF-8    LC_MESSAGES=en_US.UTF-8   
##  [7] LC_PAPER=en_US.UTF-8       LC_NAME=C                 
##  [9] LC_ADDRESS=C               LC_TELEPHONE=C            
## [11] LC_MEASUREMENT=en_US.UTF-8 LC_IDENTIFICATION=C       
## 
## time zone: US/Eastern
## tzcode source: system (glibc)
## 
## attached base packages:
## [1] stats4    stats     graphics  grDevices datasets  utils     methods  
## [8] base     
## 
## other attached packages:
## [1] patchwork_1.3.0 bbmle_1.0.25.1  glmmTMB_1.1.11  sjPlot_2.8.17  
## [5] ggplot2_3.5.2  
## 
## loaded via a namespace (and not attached):
##  [1] sass_0.4.10         generics_0.1.4      tidyr_1.3.1        
##  [4] renv_0.17.3         lattice_0.22-7      lme4_1.1-37        
##  [7] digest_0.6.37       magrittr_2.0.3      estimability_1.5.1 
## [10] evaluate_1.0.3      grid_4.5.1          mvtnorm_1.3-3      
## [13] fastmap_1.2.0       jsonlite_2.0.0      Matrix_1.7-3       
## [16] ggeffects_2.2.1     mgcv_1.9-3          purrr_1.0.4        
## [19] scales_1.3.0        numDeriv_2016.8-1.1 jquerylib_0.1.4    
## [22] Rdpack_2.6.4        reformulas_0.4.0    cli_3.6.5          
## [25] sjmisc_2.8.10       rlang_1.1.6         rbibutils_2.3      
## [28] performance_0.15.1  munsell_0.5.1       splines_4.5.1      
## [31] withr_3.0.2         cachem_1.1.0        yaml_2.3.10        
## [34] tools_4.5.1         datawizard_1.2.0    sjstats_0.19.0     
## [37] coda_0.19-4.1       nloptr_2.2.1        bdsmatrix_1.3-7    
## [40] minqa_1.2.8         dplyr_1.1.4         colorspace_2.1-1   
## [43] sjlabelled_1.2.0    boot_1.3-31         vctrs_0.6.5        
## [46] R6_2.6.1            emmeans_1.11.1      lifecycle_1.0.4    
## [49] MASS_7.3-65         insight_1.4.2       pkgconfig_2.0.3    
## [52] pillar_1.10.2       bslib_0.9.0         gtable_0.3.6       
## [55] glue_1.8.0          Rcpp_1.0.14         xfun_0.52          
## [58] tibble_3.2.1        tidyselect_1.2.1    rstudioapi_0.17.1  
## [61] knitr_1.50          farver_2.1.2        xtable_1.8-4       
## [64] htmltools_0.5.8.1   nlme_3.1-168        rmarkdown_2.29     
## [67] TMB_1.9.17          compiler_4.5.1
```

```
knitr::opts_chunk$set(fig.width = 10, fig.height = 10)
```

```
# which genotypes are in maxis but not minis?
genoMinis <- unique(df.scaled$genotype)
genoMaxis <- unique(maxis$TAG)


length(genoMinis)
```

```
## [1] 47
```

```
length(genoMaxis)
```

```
## [1] 546
```

```
sum(! genoMaxis %in% genoMinis) # 500
```

```
## [1] 500
```

### 2 clean up data

#### 2.1 change ND name

```
# rename ND to NDSU to match mini garden dataset
maxis$garden <- gsub('ND', 'NDSU', maxis$garden)
```

Cleanup using same methods as in model fitting script
‘code/transfer\_function\_linear\_mixed\_effects\_model.Rmd’

#### 2.2 set dead to 0

```
# dead trees should have a growth of zero if they were dead the whole year
# some trees died during the season - keep these growth values since it's real growth and should be related to fitness

# change these growth values from NA to 0
par(mfrow = c(1,2))
hist(maxis$GrowthIncrement_2021)
maxis[which(maxis$LIVE_Y_N_2021 == 0 & is.na(maxis$GrowthIncrement_2021)), 'GrowthIncrement_2021'] <- 0
hist(maxis$GrowthIncrement_2021)
```

```
hist(maxis$GrowthIncrement_2022)
maxis[which(maxis$LIVE_Y_N_2022 == 0 & is.na(maxis$GrowthIncrement_2022)), 'GrowthIncrement_2022'] <- 0
hist(maxis$GrowthIncrement_2022)
```

```
# save a list of which trees are alive each year
# used later to exclude dead trees when evaluating conditional model
alive21 <- rownames(maxis)[maxis$LIVE_Y_N_2021 == 1]
alive22 <- rownames(maxis)[maxis$LIVE_Y_N_2022 == 1]
```

#### 2.3 remove negative heights

```
# remove negative heights - these are either measurement error or being eaten

# 2021
hist(maxis$GrowthIncrement_2021)
```

```
sum(maxis$GrowthIncrement_2021 < 0, na.rm = T) # 65
```

```
## [1] 234
```

```
maxis[which(maxis$GrowthIncrement_2021 <0), 'GrowthIncrement_2021'] <- NA

# 2022
hist(maxis$GrowthIncrement_2022)
```

```
sum(maxis$GrowthIncrement_2022 < 0, na.rm = T) # 47
```

```
## [1] 47
```

```
maxis[which(maxis$GrowthIncrement_2022 <0), 'GrowthIncrement_2022'] <- NA

# 2023
hist(maxis$GrowthIncrement_2023)
```

```
sum(maxis$GrowthIncrement_2023 < 0, na.rm = T) # 50
```

```
## [1] 50
```

```
maxis[which(maxis$GrowthIncrement_2023 <0), 'GrowthIncrement_2023'] <- NA


# check
hist(maxis$GrowthIncrement_2021)
```

```
hist(maxis$GrowthIncrement_2022)
```

```
hist(maxis$GrowthIncrement_2023)
```

### 3 Set up model

#### 3.1 choose model variables

Use MCMT as climate variable

```
# MCMT vs height

# climate
garden_clim_colname <- 'garden_MCMT_2021'
home_clim_colname <- 'provenance_MCMT_1961_1990'
clim_label <- 'MCMT'

# phenotype
pheno_colname <- 'GrowthIncrement_2021'
pheno_label <- 'Growth Increment 2021'

###############################################

# setup variables and put into a dataframe

# set climate and phenotype variables
garden_clim <- maxis[,garden_clim_colname]
garden_clim_2 <- maxis[,garden_clim_colname]^2
home_clim <- maxis[,home_clim_colname]
home_clim_2 <- maxis[,home_clim_colname]^2
pheno <- maxis[,pheno_colname]

# random effects

# need to edit block so the VA and ND gardens don't have same block names?
# add 'maxi' to block name to distinguish from the NDSU and VA mini gardens blocks
# otherwise the same random effect will be applied even though they are actually different blocks
block <- paste('maxi', as.character(interaction(maxis$garden, maxis$BLOCK, drop = T)), sep = '.')

Pt <- maxis$k2_tricho
genotype <- as.character(maxis$TAG)
garden <- as.character(maxis$garden)
pc1 <- maxis$genetic_PC1
pc2 <- maxis$genetic_PC2
pc3 <- maxis$genetic_PC3
pc4 <- maxis$genetic_PC4
pc5 <- maxis$genetic_PC5

# put in df
df <- data.frame(pheno, garden_clim, garden_clim_2, home_clim, home_clim_2, Pt, genotype, garden, block, pc1, pc2, pc3, pc4, pc5)
rownames(df) <- rownames(maxis)

# add year and individual
df$year <- 2021
df$indiv <- rownames(maxis)

str(df)
```

```
## 'data.frame':    4928 obs. of  16 variables:
##  $ pheno        : num  0 8.23 24.64 0 10.41 ...
##  $ garden_clim  : num  -14.6 -14.6 -14.6 -14.6 -14.6 -14.6 -14.6 -14.6 -14.6 -14.6 ...
##  $ garden_clim_2: num  213 213 213 213 213 ...
##  $ home_clim    : num  -8.8 -8.8 -8.8 -8.8 -8.8 ...
##  $ home_clim_2  : num  77.4 77.4 77.4 77.4 77.4 ...
##  $ Pt           : num  0.941 0.941 0.941 0.593 0.593 ...
##  $ genotype     : chr  "201" "201" "201" "202" ...
##  $ garden       : chr  "NDSU" "NDSU" "NDSU" "NDSU" ...
##  $ block        : chr  "maxi.NDSU.1" "maxi.NDSU.2" "maxi.NDSU.3" "maxi.NDSU.1" ...
##  $ pc1          : num  -0.03339 -0.03339 -0.03339 0.00239 0.00239 ...
##  $ pc2          : num  0.0377 0.0377 0.0377 0.024 0.024 ...
##  $ pc3          : num  -0.0422 -0.0422 -0.0422 -0.0359 -0.0359 ...
##  $ pc4          : num  0.0339 0.0339 0.0339 0.0438 0.0438 ...
##  $ pc5          : num  -0.01619 -0.01619 -0.01619 0.00572 0.00572 ...
##  $ year         : num  2021 2021 2021 2021 2021 ...
##  $ indiv        : chr  "ND.201.1.5.1" "ND.201.2.19.8" "ND.201.3.35.24" "ND.202.1.6.43" ...
```

#### 3.2 scale variables

```
# subset to the variables that were used in original model
# get variables that were put into model
# variables have to be named the same in this script as in 'choose_model_variables' chunk
vars <- colnames(mod$frame)
# rename response variable
vars[1] <- 'pheno'

df <- df[,vars]
str(df)
```

```
## 'data.frame':    4928 obs. of  13 variables:
##  $ pheno        : num  0 8.23 24.64 0 10.41 ...
##  $ garden_clim  : num  -14.6 -14.6 -14.6 -14.6 -14.6 -14.6 -14.6 -14.6 -14.6 -14.6 ...
##  $ home_clim    : num  -8.8 -8.8 -8.8 -8.8 -8.8 ...
##  $ garden_clim_2: num  213 213 213 213 213 ...
##  $ home_clim_2  : num  77.4 77.4 77.4 77.4 77.4 ...
##  $ pc1          : num  -0.03339 -0.03339 -0.03339 0.00239 0.00239 ...
##  $ pc2          : num  0.0377 0.0377 0.0377 0.024 0.024 ...
##  $ pc3          : num  -0.0422 -0.0422 -0.0422 -0.0359 -0.0359 ...
##  $ genotype     : chr  "201" "201" "201" "202" ...
##  $ garden       : chr  "NDSU" "NDSU" "NDSU" "NDSU" ...
##  $ block        : chr  "maxi.NDSU.1" "maxi.NDSU.2" "maxi.NDSU.3" "maxi.NDSU.1" ...
##  $ year         : num  2021 2021 2021 2021 2021 ...
##  $ indiv        : chr  "ND.201.1.5.1" "ND.201.2.19.8" "ND.201.3.35.24" "ND.202.1.6.43" ...
```

```
# scale using scaling factors
# go through each column and check if it exists in scaling_factor
# skips non-numeric columns

df.scaled <- df

for(n in 1:ncol(df)){
  
  if(colnames(df)[n] %in% names(scaling_factor)){
    # to scale, multiply by scaling factor
    df.scaled[,n] <- df.scaled[,n] * as.numeric(scaling_factor[colnames(df)[n]])
  }
}

head(df.scaled)
```

```
##                pheno garden_clim  home_clim garden_clim_2 home_clim_2
## ND.201.1.5.1    0.00   -1.921996 -0.7437497      2.484054   0.4322519
## ND.201.2.19.8   8.23   -1.921996 -0.7437497      2.484054   0.4322519
## ND.201.3.35.24 24.64   -1.921996 -0.7437497      2.484054   0.4322519
## ND.202.1.6.43   0.00   -1.921996 -0.7437497      2.484054   0.4322519
## ND.202.2.9.10  10.41   -1.921996 -0.7437497      2.484054   0.4322519
## ND.202.3.25.22    NA   -1.921996 -0.7437497      2.484054   0.4322519
##                        pc1       pc2        pc3 genotype garden       block
## ND.201.1.5.1   -0.94192857 0.9897121 -1.0812383      201   NDSU maxi.NDSU.1
## ND.201.2.19.8  -0.94192857 0.9897121 -1.0812383      201   NDSU maxi.NDSU.2
## ND.201.3.35.24 -0.94192857 0.9897121 -1.0812383      201   NDSU maxi.NDSU.3
## ND.202.1.6.43   0.06731134 0.6310293 -0.9192134      202   NDSU maxi.NDSU.1
## ND.202.2.9.10   0.06731134 0.6310293 -0.9192134      202   NDSU maxi.NDSU.2
## ND.202.3.25.22  0.06731134 0.6310293 -0.9192134      202   NDSU maxi.NDSU.3
##                year          indiv
## ND.201.1.5.1   2021   ND.201.1.5.1
## ND.201.2.19.8  2021  ND.201.2.19.8
## ND.201.3.35.24 2021 ND.201.3.35.24
## ND.202.1.6.43  2021  ND.202.1.6.43
## ND.202.2.9.10  2021  ND.202.2.9.10
## ND.202.3.25.22 2021 ND.202.3.25.22
```

### 4 Predict growth and mortality from model

#### 4.1 Functions

```
# modified from 'code/transfer_function_linear_mixed_effects_model.Rmd'

# function to calculate correlation between predicted and observed values for a model (essentially the R value)

# mod is the model
# it pulls the dataset from the model object, which allows us to remove rows with NAs for just the variables used in this model
# re.form argument goes to predict(); to include random effects set to NULL, to set random effects to zero set to NA

model_R <- function(mod, newdata, type = 'response', re.form = NULL, se.fit = FALSE){
  
  # setup
  # include random effects in predictions or set random effects to zero?
  # this is the re.form argument in predict()
  #re.form <- ifelse(include_REs == TRUE, eval(NULL), NA)
  
  df <- newdata
  
  # make new dataframe with no NA values
  noNA <- df[complete.cases(df),]

  
  # predict phenotype (generally height)
  noNA$pred <- predict(mod, newdata = noNA, type = type, re.form = re.form, se.fit = se.fit)
  
  # un-log-transform
  noNA$pred <- exp(noNA$pred) - 1
  
  # if type = conditional, remove dead trees (which are modeled separately in zero-inflated model)
  if(type == 'conditional'){
    noNA <- noNA[alive21,]
  }
  
  # calculate correlation
  cor <- cor.test(noNA$pheno, noNA$pred)
  
  return(list(actual = noNA$pheno, predicted = noNA$pred, cor = cor, data_noNAs = noNA))
  
}

# test
# cor <- model_R(mod, re.form = NULL)

# plot which takes output of model_R

plot_predicted_vs_actual <- function(input, title = NULL, col = rgb(0,0,0,0.1), col_1to1 = 'red', col_fit = 'blue', legend = TRUE){
  
  pval <- round(input$cor$p.value, 4)
  cor_est <- as.numeric(round(input$cor$estimate, 3))
  main <- paste(title, '\n', 'R = ', cor_est, ' | p = ', pval, sep = '')
  
  # if pval rounds to zero, display as <0.001
  if(pval == 0){
    pval <- '<0.001'
    main <- paste(title, '\n', 'R = ', cor_est, ' | p < 0.001', sep = '')
  } else {
    main <- paste(title, '\n', 'R = ', cor_est, ' | p = ', pval, sep = '')
  }
  
  maxVal <- max(c(input$actual, input$predicted), na.rm = T)
  
  plot(input$actual, input$predicted, 
       pch = 16, 
       col = col, 
       xlab = "Actual growth increment (cm)", 
       ylab = "Predicted growth increment (cm)",
       title(main, adj = 0),
       xlim = c(0, maxVal),
       ylim = c(0, maxVal))
  abline(0, 1, col = col_1to1, lty = 2) # 1to1 line
  abline(lm(input$predicted ~ input$actual), col = col_fit) # fit line
  
  # add text
  if(legend == TRUE){
    legend('bottomright', fill = c(col_1to1, col_fit), legend = c('1:1 line', 'Best fit'))
  }
}

#######
# for plotting actual mortality vs mortality probability - need logistic model
# combined the two functions used for overall/conditional models into one to pass the p-value directly to the plot
model_plot_R_mortality <- function(mod, newdata, re.form = NULL, se.fit = FALSE, # model parameters
                              title = NULL, # plot parameters
                              col = rgb(0,0,0,0.1), 
                              col_fit = 'blue'){
  
  # setup
  # include random effects in predictions or set random effects to zero?
  # this is the re.form argument in predict()
  #re.form <- ifelse(include_REs == TRUE, eval(NULL), NA)
  
  df <- newdata

  # first column is phenotype
  colnames(df)[1] <- 'pheno'
  
  # make new dataframe with no NA values
  noNA <- df[complete.cases(df),]
  
  # predict phenotype - here, probability of mortality
  noNA$pred <- predict(mod, newdata = noNA, type = 'zprob', re.form = re.form, se.fit = se.fit)
  
  # convert growth increment to binary 0/1
  # if value is not 0, convert to 1
  noNA$alive <- ifelse(noNA$pheno == 0, 0, 1)
  
  # logistic model
  mort <- glm(alive ~ pred, family = 'binomial', data = noNA)
  mort.info <- summary(mort)
  
  # combine outputs into list  
  out <- list(actual = noNA$alive, predicted = noNA$pred, cor = cor, data_noNAs = noNA)
  
  # plot
  pval <-  round(mort.info$coefficients[2,'Pr(>|z|)'], 3)
  # if pval rounds to zero, display as <0.001
  if(pval == 0){
    pval <- '<0.001'
    main <- paste(title, '\np < 0.001', sep = '')
  } else {
    main <- paste(title, '\np = ', pval, sep = '')
  }
  
  # save plot to output
  out$plot <- ggplot(data = noNA, aes(x = pred, y = alive,)) +
  geom_jitter(height = 0.05, width = 0, color = col) +
  xlab('Probability of Mortality') +
  ylab('Actual Survival')+
    ggtitle(main) +
geom_smooth(method = glm, formula = y ~ x, method.args = list(family = binomial), col = col_fit)
  
  return(out)
  
}
```

#### 4.2 Predict for maxi garden data

```
# predict growth increment for each tree in maxi gardens, based on genetic PCs, home climate, and garden climate, and compare predictions to actual values

# see ?predict.glmmTMB

# This will give a warning about new random effects, because some genotypes and gardens were not included in training dataset (mini gardens)

# Warning messages:
# 1: In checkTerms(data.tmb1$terms, data.tmb0$terms) :
#   Predicting new random effect levels for terms: 1 | genotype, 1 | block:garden, 1 | garden, 1 | indiv
# Disable this warning with 'allow.new.levels=TRUE'

# 2: In checkTerms(data.tmb1$termszi, data.tmb0$termszi) :
#   Predicting new random effect levels for terms: 1 | genotype, 1 | block:garden, 1 | garden, 1 | indiv
# Disable this warning with 'allow.new.levels=TRUE

par(mfrow = c(2,2))
# overall model
plot_predicted_vs_actual(model_R(mod, newdata =  df.scaled, re.form = NULL), title = 'overall, with random effects')
plot_predicted_vs_actual(model_R(mod, newdata = df.scaled, re.form = NA), title = 'overall, no random effects')
# conditional model
plot_predicted_vs_actual(model_R(mod, type = 'conditional', newdata =  df.scaled, re.form = NULL), title = 'conditional, with random effects')
plot_predicted_vs_actual(model_R(mod, type = 'conditional', newdata = df.scaled, re.form = NA), title = 'conditional, no random effects')
```

```
# for each garden

# overall response
# png(file = paste('results/model_prediction/maxi_growth_predictions_vs_actual_from_model_2yearModel_', garden_clim_colname, '1to1.png', sep = ''),
# height = 8,
# width = 12,
# res = 300,
# units = 'in')

par(mfrow = c(3,4))
gards <- unique(df.scaled$garden)

for(n in 1:length(gards)){
  
  gard <- gards[n]
  # overall model
  plot_predicted_vs_actual(model_R(mod, df.scaled[df.scaled$garden == gard,], re.form = NULL),
                           title = paste(gard, 'overall, with random effects'),
                           legend = ifelse(n == 1, T, F))
  plot_predicted_vs_actual(model_R(mod, df.scaled[df.scaled$garden == gard,], re.form = NA), 
                           title = paste(gard, 'overall, no random effects'), legend = F)
  # conditional model
    plot_predicted_vs_actual(model_R(mod, df.scaled[df.scaled$garden == gard,], re.form = NULL, type = 'conditional'),
                           title = paste(gard, 'conditional, with random effects'), legend = F)
  plot_predicted_vs_actual(model_R(mod, df.scaled[df.scaled$garden == gard,], re.form = NA, type = 'conditional'), 
                           title = paste(gard, 'conditional, no random effects'), legend = F)
  
}
```

```
#dev.off()


# just conditional with random effects, to match the mini gardens validation plots

png(file = paste('results/model_prediction/validation_MaxiPredictions_2yearModel', garden_clim_colname, '_vs_', pheno_colname, '.png', sep = ''), width = 9, height = 4, res = 300, units = 'in')

cols <- c("#2B3D26", "#DCD300", "#2b3e85")

par(mfrow = c(1,3))

for(n in 1:length(gards)){
  
  gard <- gards[n]
  col <- adjustcolor(cols[n], alpha = 0.6)
  
  plot_predicted_vs_actual(model_R(mod, df.scaled[df.scaled$garden == gard,], re.form = NA, type = 'conditional'), 
                           title = paste(gard, '(Maxi)'),
                           col = col,
                           col_fit = col,
                           col_1to1 = 'grey',
                           legend = F)
  
}

dev.off()
```

```
## png 
##   2
```

```
# mortality probability

mort_plots <- list()
for(n in 1:length(gards)){
  
  
  gard <- gards[n]
  
  newdata <- df.scaled[df.scaled$garden == gard,]

  
  col <- adjustcolor(cols[n], alpha = 0.6)
   
  # zero-inflated model
  out <-  model_plot_R_mortality(mod, 
                                     newdata = newdata, 
                                     re.form = NULL, 
                                     title = gard,
                                     col = col,
                                     col_fit = cols[n])
  p <- out$plot
  #plot(p)
  
  mort_plots[[n]] <- p
  
}


wrap_plots(mort_plots, ncol = 3)
```

```
ggsave(filename = paste('results/model_prediction/validation_MaxiPredictions_zero-inflated_2yearModel', garden_clim_colname, '_vs_', pheno_colname, '.png', sep = ''), 
 width = 12, height = 5)

   
    

# test prediction ability for each genotype

genos <- unique(df.scaled$genotype)


geno.r.resp <- rep(NA, length(genos))
geno.r.cond <- rep(NA, length(genos))


for(n in 1:length(genos)){
  
  geno <- genos[n]
  
  sub <- df.scaled[df.scaled$genotype == geno,]
  
  # checks before trying to predict
  # 1. need to have at least three observations
  # 2. if genotype is missing information, skip it
  # ok if phenotype has some NAs
  if(length(sub$pheno[! is.na(sub$pheno)]) > 2 &
     sum(is.na(sub[,colnames(sub != 'pheno')])) == 0){
    
    resp <- model_R(mod, sub, type = 'response', re.form = NULL)
    
    # only run conditional if there are enough live trees
    if(nrow(sub[rownames(sub) %in% alive21,]) > 2){
       cond <- model_R(mod, sub, type = 'conditional', re.form = NULL)
    }
   
    
    geno.r.resp[n] <- resp$cor$estimate
    geno.r.cond[n] <- cond$cor$estimate
    
  }
  
  
  # if(n %% 20 == 0){
  #   cat(paste('done with', n, '\n'))
  # }
}

names(geno.r.resp) <- genos
names(geno.r.cond) <- genos

par(mfrow = c(1,2))
hist(geno.r.resp)
hist(geno.r.cond)
```

```
# which genotypes are more predictable?

geno_info <- as.data.frame(matrix(nrow= length(genos), ncol = 6))
colnames(geno_info) <- c('k2_tricho', 'genetic_PC1', 'genetic_PC2', 'genetic_PC3', 'TRANSECT', 'in_minis' )

rownames(geno_info) <- genos

geno_colnames <- c('k2_tricho', 'genetic_PC1', 'genetic_PC2', 'genetic_PC3', 'TRANSECT')

for(n in 1:nrow(geno_info)){
  
  geno <- rownames(geno_info)[n]
  
  # get first row with info for this genotype
  geno_row <- maxis[match(geno, maxis$TAG), geno_colnames]
  
  geno_info[n, geno_colnames] <- geno_row
  
  # is the genotype in the mini gardens?
  geno_info[n, 'in_minis'] <- geno %in% mod$frame$genotype

}

geno_info$Rval_resp <- geno.r.resp
geno_info$Rval_cond <- geno.r.cond

# there are a lot of missing values due to low sample size for some genotypes. Look at using residuals instead? or some other measure of accuracy?

# are predictions better for genotypes that were included in the minis (training dataset)?
# average R values are similar, but more variance in accuracy for non-mini genotypes, and most of the negative R values are non-mini genotypes
par(mfrow = c(1,2), mar = c(8,4,4,3))
boxplot(geno_info$Rval_resp ~ geno_info$in_minis)
abline(h = 0, col = 'blue')
boxplot(geno_info$Rval_cond ~ geno_info$in_minis)
abline(h = 0, col = 'blue')
```

```
plot(density(geno_info[geno_info$in_minis == F,'Rval_resp'], na.rm = T))

# is there variance across transects?
# not too much variance across transects, including WY which was not in minis
# lost of variation in accuracy for crowsnest conditional model
par(mfrow = c(1,2), mar = c(8,4,4,3))
```

```
boxplot(geno_info$Rval_resp ~ geno_info$TRANSECT, las = 2, xlab = '')
abline(h = 0, col = 'blue')
boxplot(geno_info$Rval_cond ~ geno_info$TRANSECT, las = 2, xlab = '')
abline(h = 0, col = 'blue')
```

```
# plot cor against genetic info
# genotypes with low R values are both balsam and tricho

par(mfrow = c(1,2))

plot(geno_info$k2_tricho, geno_info$Rval_resp)
abline(h = 0, col = 'blue')
plot(geno_info$k2_tricho, geno_info$Rval_cond)
abline(h = 0, col = 'blue')
```

```
plot(geno_info$genetic_PC1, geno_info$Rval_resp)
abline(h = 0, col = 'blue')
plot(geno_info$genetic_PC1, geno_info$Rval_cond)
abline(h = 0, col = 'blue')
```

```
plot(geno_info$genetic_PC2, geno_info$Rval_resp)
abline(h = 0, col = 'blue')
plot(geno_info$genetic_PC2, geno_info$Rval_cond)
abline(h = 0, col = 'blue')
```

```
plot(geno_info$genetic_PC3, geno_info$Rval_resp)
abline(h = 0, col = 'blue')
plot(geno_info$genetic_PC3, geno_info$Rval_cond)
abline(h = 0, col = 'blue')
```
