## Supplementary figures and images for "Variation in responses to temperature in admixed *Populus* genotypes predicts geographic shifts in regions where hybrids are favored"

### Figure S12

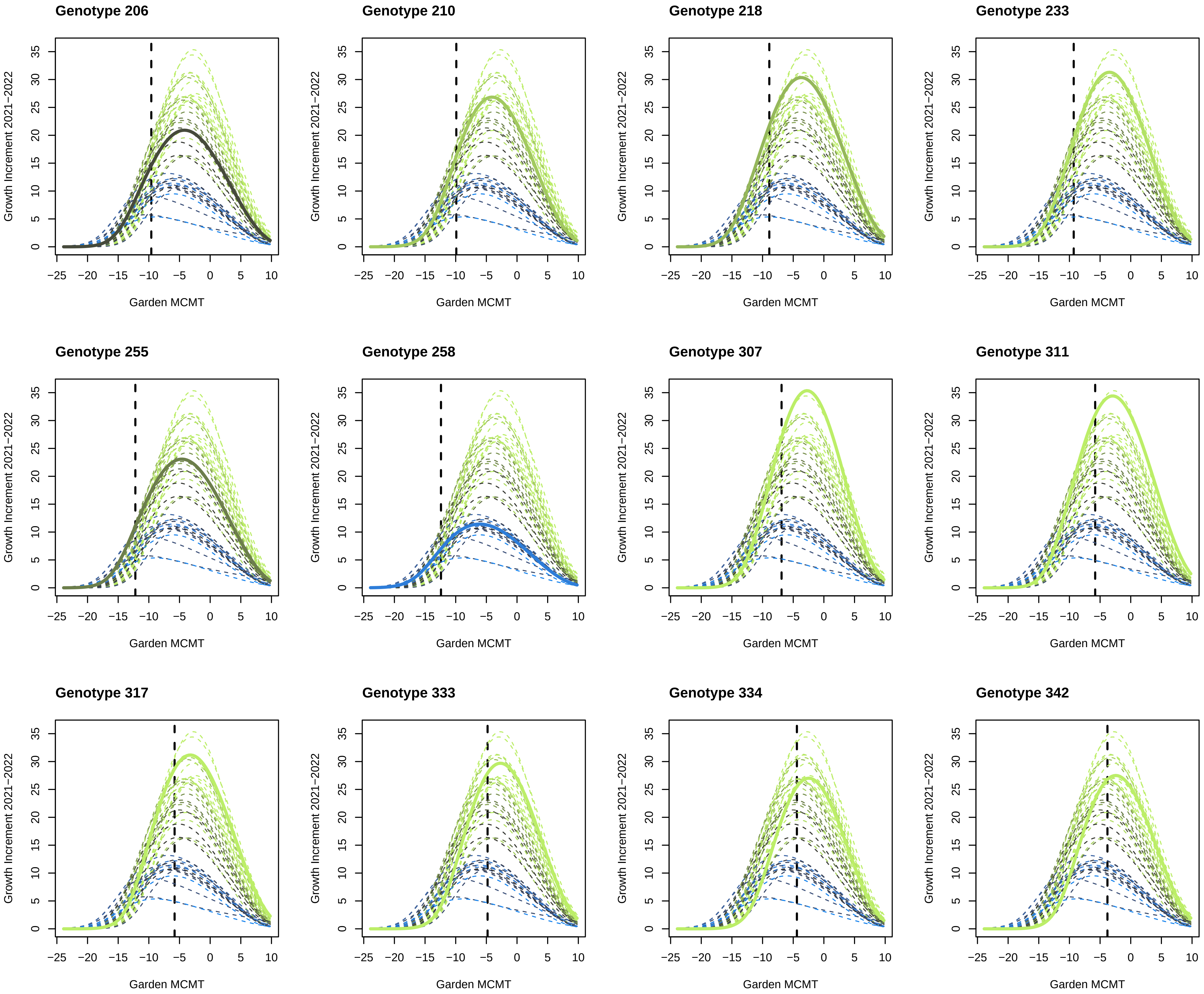

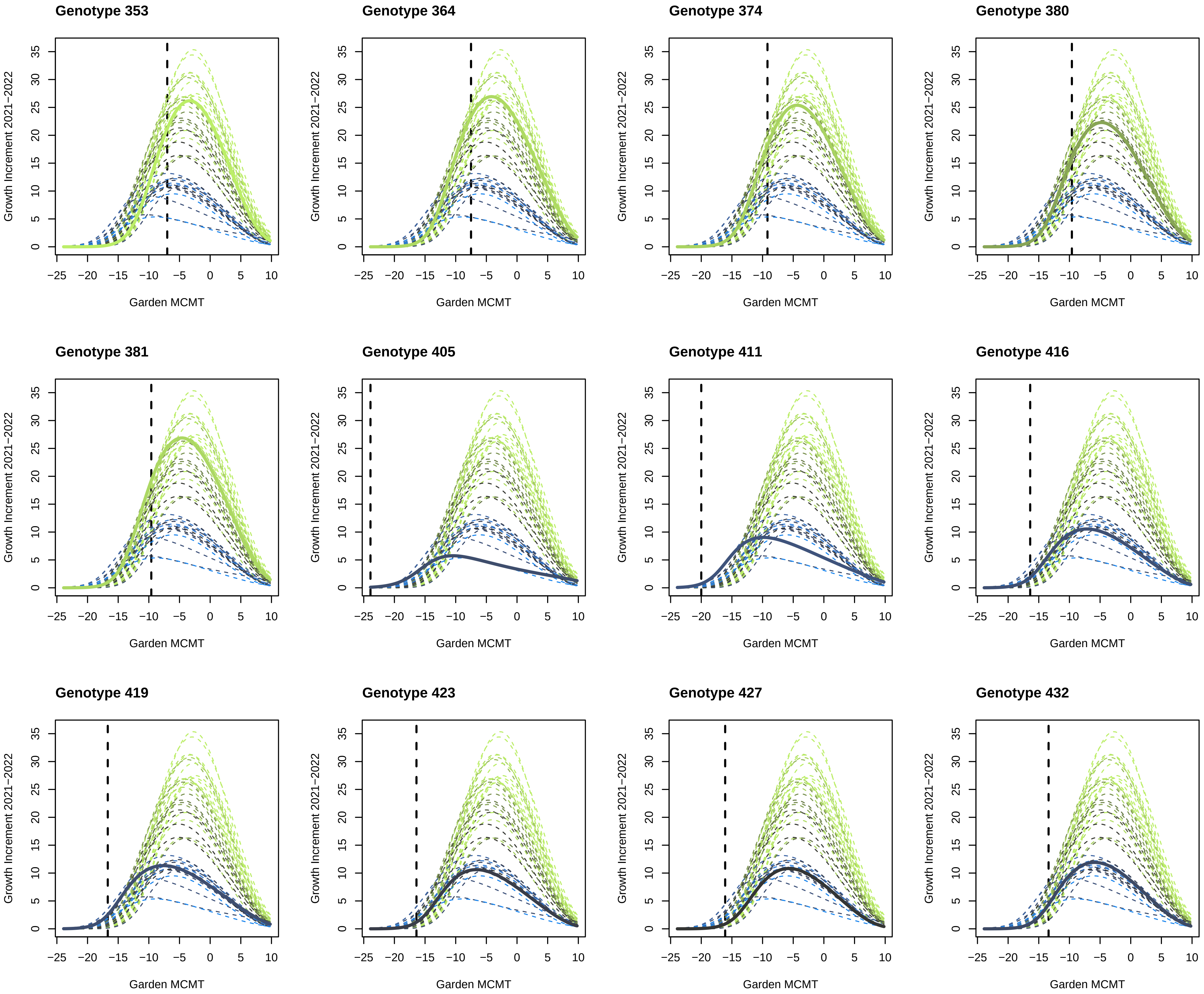

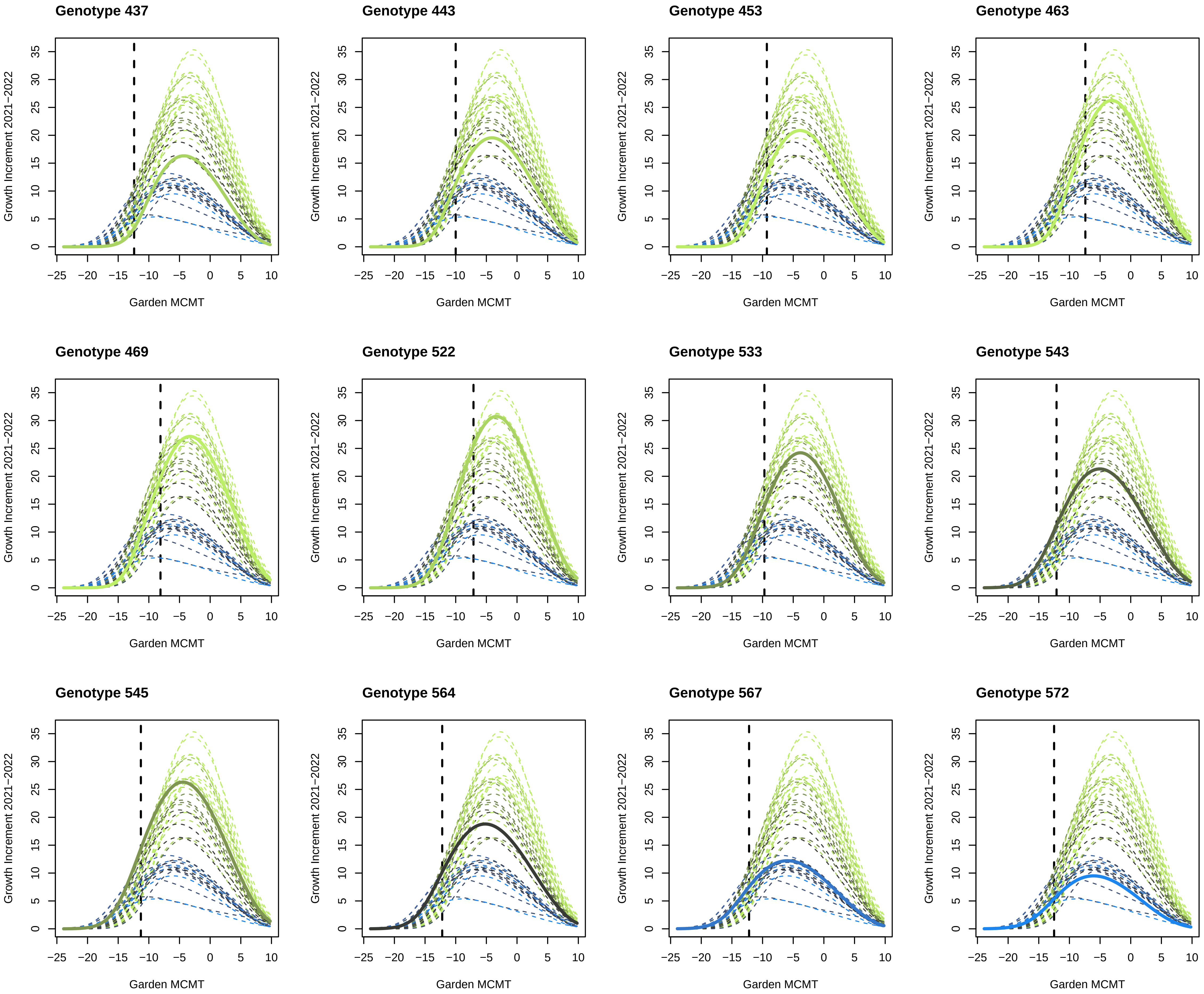

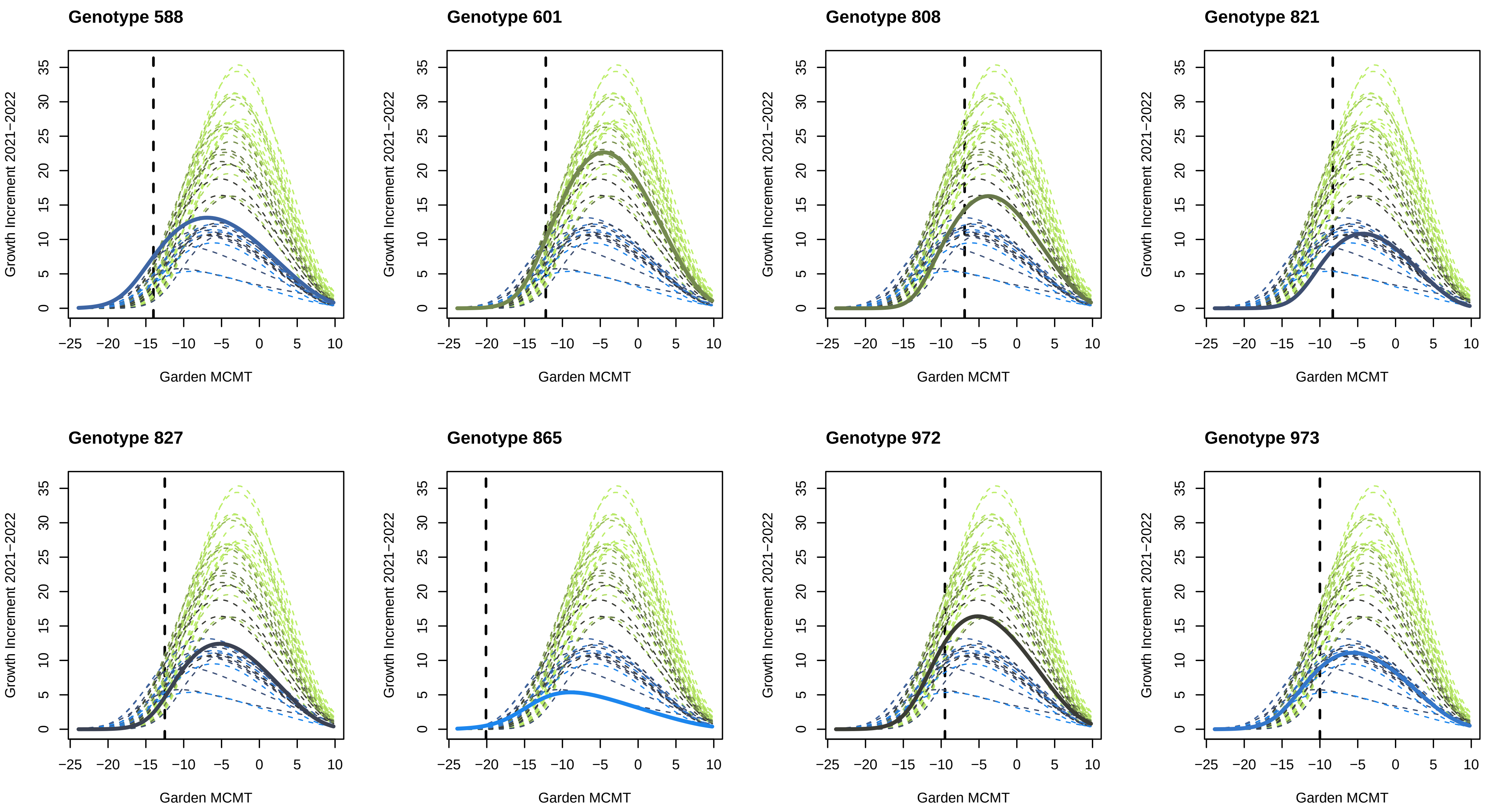
